# The nunatak and *tabula rasa* hypotheses may be compatible: the European phylogeography of a riparian earthworm

**DOI:** 10.1101/2024.01.26.576623

**Authors:** Irene de Sosa, Daniel F Marchán, Christer Erséus, Emmanuel Lapied, Misel Jelic, Aleksandra Jabłońska, Timea Szederjesi, Ana Almodóvar, Marta Novo, Darío Díaz Cosín

## Abstract

The *tabula rasa* hypothesis of postglacial immigration supports the notion that species now found in northern European areas must have been recently recolonized from historical refugia. Until the 1960s, however, there was almost complete consensus that disjunctions and endemism in the North Atlantic region of Europe could not be explained without in situ survival during glacial periods (the nunatak hypothesis). Although some earthworms can survive in permafrost and tolerate cold conditions, it is generally believed that most earthworms were eradicated from northern latitudes during the Last Glacial Maximum. To test which hypothesis explains the phylogeography of the riparian and parthenogenetic earthworm *Eiseniella tetraedra*, we collected 1,640 specimens from 19 different countries in Europe. We examined three molecular markers (COI, 16S and 28S) and their morphology. Eleven lineages were found, nested in five clades. Clade I was more prevalent in cold biogeographical regions such as the continental, the Atlantic or even the Arctic, while clade II was prevalent in Mediterranean regions. We investigated their potential niches through Species Distribution Models, which agreed with the distribution trends. The presence of restricted clades in the Iberian and Scandinavian peninsulas, as well as in Eastern Europe, suggests that these three regions served as refugia during the Last Glacial Maximum. Thus, both hypotheses were necessary to explain the actual distribution of this shore-dwelling earthworm.

## 1. Introduction

Nowadays, earthworms include about 6,500 species described, of which 3,000-3,500 are valid (Csuzdi, 2012). The family Lumbricidae (Rafinesque-Schmaltz, 1815) includes about 300 species (Csuzdi, 2012) and originated in the Lower Cretaceous in the Holarctic region (Dominguez et al., 2015). Earthworms belonging to this family are the most abundant invertebrates in the soil of temperate regions and account for 90% of invertebrate biomass (Edwards, 2004). Earthworms are not able to live in permafrost for long periods of time (Holmstrup et al., 1991). Therefore, it is generally assumed that earthworms were eradicated from northern latitudes during the Last Glacial Maximum (LGM) (Tiunov et al., 2006), so that species now found in northern European areas must have been recently recolonized from historical refugia such as the southern European peninsulas. This basic expansion-contraction model is known as the “*tabula rasa*” hypothesis. In contrast, the nunatak hypothesis suggests that some ice-free refuge existed in northern Europe, such as mountains rising above inland ice or coastal ice-free refuges, where the biota could survive. When the ice melted, plants and animals recolonized previously ice-covered areas from these northern refugia. There is now strong geological evidence suggesting that some nunataks and ice-free coastal shelves existed within the maximum limits of the last glacial period 25,000– 10,000 years ago (Hansen et al., 2006). Although nunatak hypothesis seems unlikely for earthworms, it was proposed for Fennoscandia (Fridolin 1936, Stöp-Bowitz 1969) and Greenland (Hansen et al. 2006). Moreover, some hardy Lumbricidae species could survive the last glacial period in ice-free refugia in association with some arctic plants (Julin, 1949; Stöp-Bowitz, 1969). *Dendrobaena octaedra* (Savigny, 1826) overwinters in frozen ground either as adults or in cocoons. In Greenland, it exhibited high genetic diversity, which, combined with its high frost tolerance, suggests its survival in ice-refuge in Greenland (Hansen et al., 2006), following the nunatak hypothesis. Phylogeographic studies can help us determine the history of diversification and dispersal of a species and earthworms with high genetic diversity are good models for these studies (Fernández et al., 2013; Shekhovtsov et al., 2020; de Sosa et al., 2023a). A phylogeographical study covering the whole Europe, might therefore throw light on these hypotheses.

*Eiseniella tetraedra* (Savigny, 1826) is a parthenogenetic and tetraploid (Casellato, 1897) earthworm with a riparian lifestyle that inhabits margins of freshwater. It has a worldwide distribution and is therefore referred as a cosmopolitan earthworm (Blakemore, 2006). *E. tetraedra* showed a high genetic diversity in the Iberian Peninsula with eight different lineages nested in two clades. This genetic diversity was distributed according to three environmental factors: temperature, precipitations and pH. Thus, three linages were found in the northern half of the Iberian Peninsula (namely B, C and D), three lineages were distributed throughout the Peninsula (A,E and F) and two of them were restricted geographically and appeared in only one (H) or two (G) populations (de Sosa et al., 2023a). Jadvikar et al. (2020) found only six lineages in Iran, probably introduced by human activity, while de Sosa et al. (2017) found only one lineage in Scandinavia, which could be due to a limited number of samples. *E. tetraedra* was found to the north of the 65 parallel (Haraldsen and Engelstad 1998), and was detected as far as the northern coast of the Scandinavian Peninsula (Terhivuo 1988). It was also found in Iceland (Blakemore, 2007). According to these presences, *E. tetraedra* seems to be a cold tolerant earthworm. However, de Sosa et al. (2022) showed that *E. tetraedra* adults are not good candidates for freezing conditions due to the downregulation of the respiratory chain.

Ecological Niche Modeling (ENM) with MaxEnt (Phillips et al., 2006) has facilitated ecological inference in soil due to its high power when only presence data are included. It has been implemented in several groups such as termites (Maynard et al., 2015), beetles (Crawford and Hoagland, 2010), millipedes (Marek et al., 2012) and earthworms (Marchán et al., 2015; Marchán et al., 2016). Moreover, macroecological preferences in soil communities were studied by Decaëns (2010). The macroecological preferences of *E. tetraedra* have been studied (Si-Moussi, 2020) and soil texture was the most discriminating factor for this species, but nothing is known about its clades. Marchán et al. (2016) found ecological divergence among four different clades of the earthworm family Hormogastridae, which showed different environmental responses and preferred different habitats.

The present study investigated the genetic variation and phylogeographic relationships of a large number of populations of *E. tetraedra* collected from nineteen countries in Europe. The aim of this study was: i) to examine the genetic diversity of *E. tetraedra* in Europe; ii) to infer its phylogeography in this continent and iv) to test macroecological preferences for clades.

## 2. Material and methods

### 2.1. Earthworm sampling and morphological analyses

We collected 1,640 specimens from 19 different countries: Belgium, Bulgaria, Czech Republic, Finland, France, Germany, Greece, Israel, Italy, Netherlands, Norway, Poland, Portugal, Russia, Slovakia, Spain, Sweden, Turkey and United Kingdom (Supplementary File 1). Earthworms were collected by hand-sorting and fixed in 96% ethanol and stored at −20°C. Whenever possible, we selected ten individuals per locality, and a portion of the posterior body section was excised and carefully cleaned under a stereomicroscope to remove gut and soil particles. Tegument samples were stored in ethanol and preserved at −20°C until DNA extraction. Morphological characters were studied on 1,193 specimens, focusing on: length, dry weight, number of segments, position of clitellum and *tubercula pubertatis*, position of male pores, number and position of seminal vesicles, spermathecae and spermiducal funnels.

### 2.2. DNA extraction, gene amplification and sequencing

Total genomic DNA was extracted using the Speedtools Tissue DNA Kit (Biotools). We amplified a fragment of cytochrome *c* oxidase subunit I (COI) in all specimens, and we chose two specimens per site and lineage (see Results) for amplification of a fragment containing 16S rRNA + tRNAs Leu, Ala and Ser (16S), and a fragment of 28S rRNA (28S) (see Supplementary File 1). For COI (632 bp) primer sequences and polymerase chain reactions (PCR) followed Pop et al. (2003). For 16S-tRNAs (775 bp) and 28S (804 bp) primer sequences and PCR conditions followed Fernández et al. (2015). All PCRs were specific, resolved via 1% agarose gel electrophoresis and were visualised with GelRed stain (Biotium). PCR products were purified using ExoSAP-IT reagent (ThermoFisher Scientific) and sequenced by Macrogen Spain Inc.

### 2.3. Phylogenetic analyses and genetic variability

Sequences of each gene were aligned in MAFFT v.7 (Katoh and Standley, 2003) using default settings and concatenated with BioEdit v7.0.9 (Hall, 1999). Phylogenetic analyses with the concatenated sequence of the three genes (2,213 bp) included Bayesian inference (BI) using MrBayes v.3.1.2 (Ronquist and Huelsenbeck, 2003) and Maximum Likelihood (ML) with RaxML v.7.03 software (Stamatakis, 2006) both implemented in Cipres Science Gateway v.3.3 (Miller et al., 2010). Phylogenetic trees obtained were visualised in FigTree v1.3.1 (Morariu et al., 2008). The best-fitting substitution model selected by jModelTest2 (Darriba et al., 2012) for each partition was GTR+Γ+I. The Bayesian analyses consisted of two parallel runs of ten million of generations, sampling 10,000 trees every 1,000^th^ generation, starting the analysis with a random tree. 20% of the trees were discarded as burn-in. ML analysis with rapid bootstrapping was performed with 1,000 replicates. Sequences of *Lumbricus rubellus* Hoffmeister, 1843, *Dendrobaena byblica* Rosa, 1893, *Eiseniona oliveirae* (Rosa, 1894), *Prosellodrilus biauriculatus* Bouché, 1972 and *Carpetania matritensis* Marchán et al., 2020, were retrieved from GenBank and used as outgroups (Supplementary File 2). In addition, all nucleotide sequences generated in this study were deposited in the GenBank (Supplementary File 2).

Uncorrected pairwise distances for COI were calculated within and between main clades (see Results). We also examined haplotype and nucleotide diversity for clades and for the three peninsulas sampled: Iberian, Italian and Scandinavian.

### 2.4. Ecological niche modeling

Presence data were aggregated according to the two principal clades (I and II) recovered in the phylogenetic analyses (see Results). The following large-scale variables were chosen as predictor variables. They are the same (or equivalent) as the variables found as the most influential for the distribution of *Eiseniella tetraedra* by Si-moussi (2020).

-*Bioclimatic variables* (downloaded from Worldclim - http://www.worldclim.org/ accessed 01/12/2020)

Min Temperature of Coldest Month (BIO6)

Temperature Annual Range (BIO7)

Precipitation of Wettest Month (BIO13)

-*Soil variables*

Lithology (PARMA), represented by the PAR-MATDOM2 (Second level code for the dominant parent material of the STU) layer obtained from the European Soil Database Raster Library 1 km _ 1 km (http://eusoils.jrc.ec.europa.eu/ESDB_Archive/ESDB_data_1k_raster_intro/ESDB_1k_raster_data_intro.ht ml accessed 01/12/2020).

Soil crusting class (CRUST) layer obtained from the European Soil Database Raster Library 1 km _ 1 km (http://eusoils.jrc.ec.europa.eu/ESDB_Archive/ESDB_data_1k_raster_intro/ESDB_1k_raster_data_intro.ht ml accessed 01/12/2020).

Topsoil available water capacity (AWC) layer obtained from the European Soil Database Raster Library 1 km

_ 1 km (http://eusoils.jrc.ec.europa.eu/ESDB_Archive/ESDB_data_1k_raster_intro/ESDB_1k_raster_data_intro.ht ml accessed 01/12/2020).

-*Biotic variables*

Vegetation and dominant land use (CLC) were represented by the CORINE 2018 Land Cover layer (version v.2020_20u1: https://land.copernicus.eu/pan-european/corine-land-cover/clc2018?tab=download).

Ecological niche models were obtained using the R package ‘SSDM’ (Schmitt et al. 2017). Presence (localities where the target clade was found) and absence (localities were the other clade - but not the target clade-was found) data were used as input. Ensemble species distribution models (ESDMs) were built by combining the algorithms (‘MAXENT’, ‘GLM’, ‘CTA’ and ‘MARS’) producing kappa values greater than 0.5, with three repetitions for each algorithm.

## 3. Results

### 3.1. Phylogenetic analysis

Bayesian and Maximum Likelihood approaches yielded trees with congruent topology for *E. tetraedra* (Figure 1). All sequences, except one (LSWED303-11 from Sweden) were nested in five different and well supported clades (labeled I to V). Clades I and II included four different lineages (B-C-D-G, A-H-E-F, respectively), whereas clades III to V comprised only one each (I, J and K). Clade IV showed the lowest genetic diversity with only three different haplotypes, while clade II was the most diverse with 35.76% of the total haplotypes.

**Figure 1.**
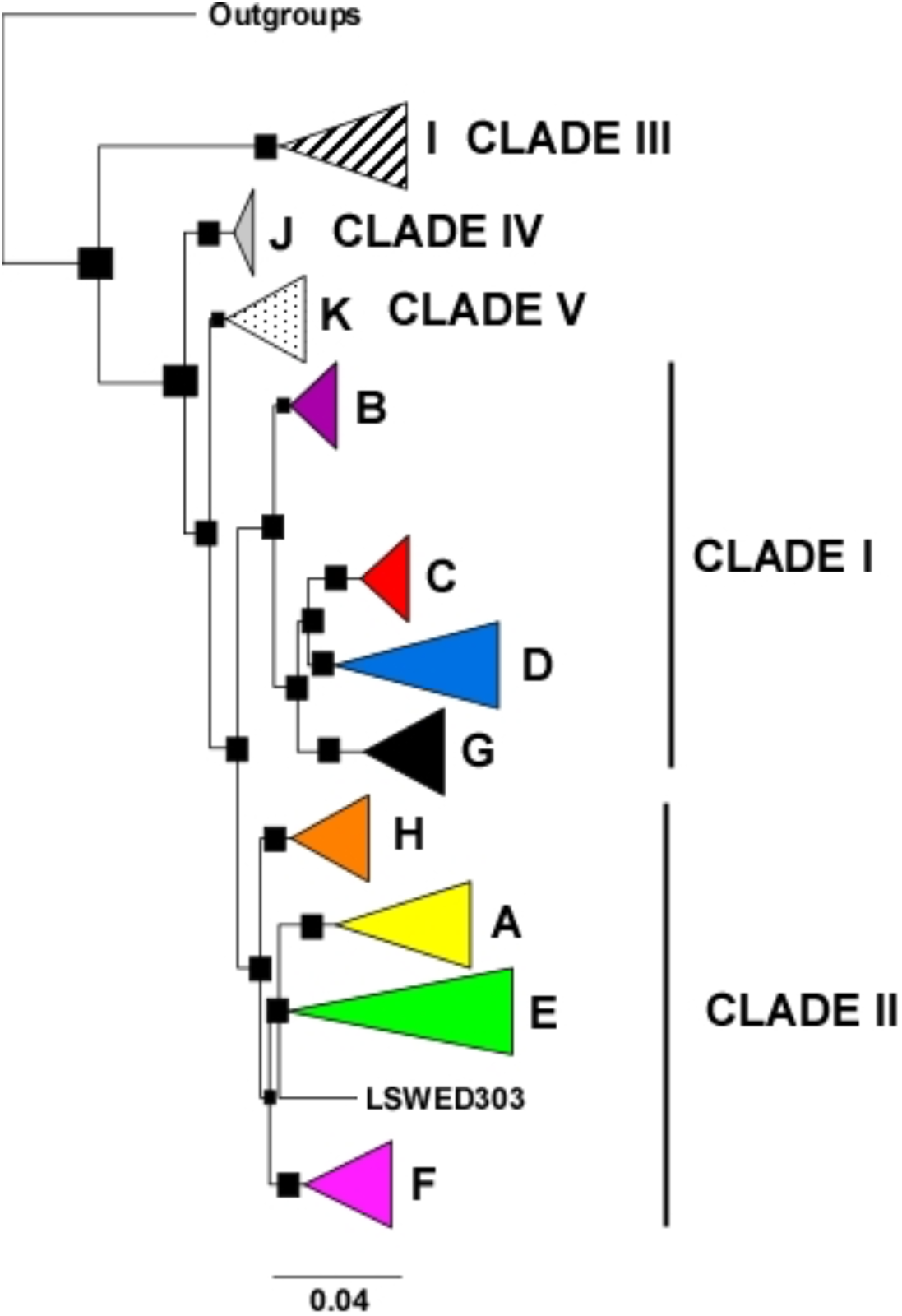
Bayesian inference (BI) of the phylogenetic tree based on concatenated sequences of COI (1,038), 16S (262) and 28S (378). Posterior probability/bootstrap support values (of Maximum Likelihood Analysis, ML) are shown as black squares when higher than 0.9/0.7 (BI/ML). The scale bar represents 0.04 substitutions per position. Colors and names of lineages (A-H) and number of clades (I and II) are the same as in de Sosa et al., 2023a and in Figures 2 and 3. LSWED303-11 correspond to a mature specimen from Frösslundabäcken Stream, Öland, Sweden.

### 3.2. Lineages distribution

The distribution of lineages in Europe is shown in Figures 2 and 3. In the Iberian Peninsula, the northern area was more diverse than the southern one, and lineages D and H were present only in this area of Europe (Figures 2 and 3). We found all lineages in the Scandinavian Peninsula (Figure 2.5), except those which were present in only a few populations (D, G, H and I). Lineage B was clearly predominant in all areas and lineage J and K occurred mostly in this countries. The Italian Peninsula is probably one of the least diverse areas (Figure 2.2). Only lineages from the clade II (A, E and F) were found. North-South differences were observed in the Italian Peninsula. Lineage A predominated in the northern area, while lineage F was more numerous in the central and southern areas. The two populations of United Kingdom showed high diversity (Figure 2.1). Only lineages A, E and F were represented at Central Europe with the exception of two individuals of lineages G in the Netherlands and J in Slovakia (Figure 2.3). The eastern regions of Europe were less represented and lineages A, F and J were found (Figure 2.1). The distribution of lineages in France is shown in Figure 2.4. In northern France, lineage B was predominant, while South France showed greater diversity, especially in the Pyrenees. Therefore, clades I and II were widespread in Europe, although clade I was most common in cold areas and clade II in temperate ones. In contrast, clades III, IV and V had a restricted distribution, with clade III occurring mainly in Eastern Europe and clades IV and V found mainly in Scandinavian Peninsula.

**Figure 2.**
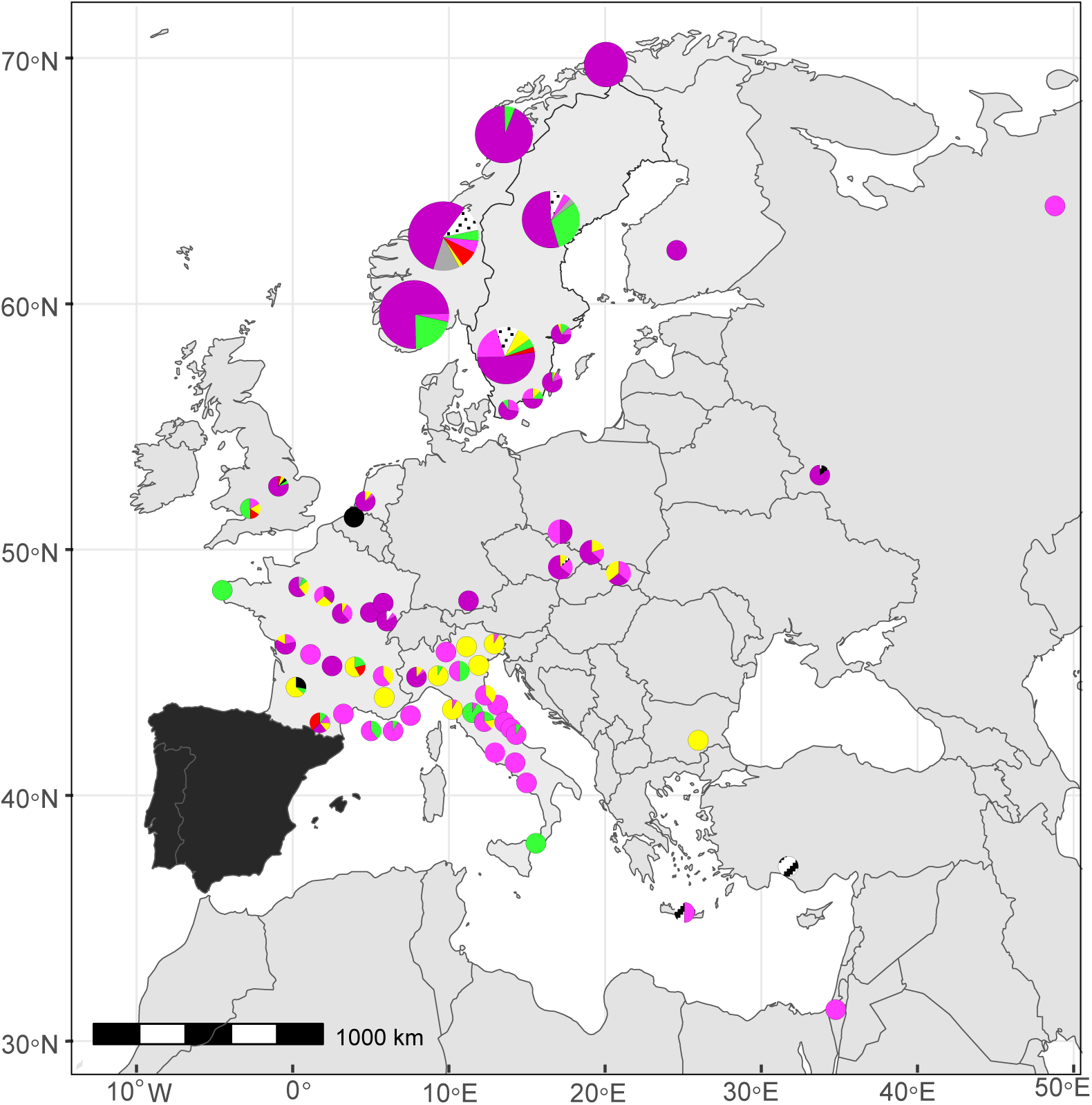
Lineages distribution in Europe. Proportion of individuals from each genetic lineage in each locality is represented in pie charts. 1: The pie charts shows distribution of lineages in distinct parts of Europe. 2: Lineages distribution in the Italian Peninsula. 3: Lineages distribution in Slovakia, Poland and Czech Republic. 4: Lineages distribution in France. 5: Lineages distribution in the Scandinavian Peninsula. Many of the lineage pie charts show information from several adjacent populations. The size of each circle is proportional to the number of localities it includes. Colors used are the same as in Figure 1. Details of area 6 are showed in Figure 3.

**Figure 3.**
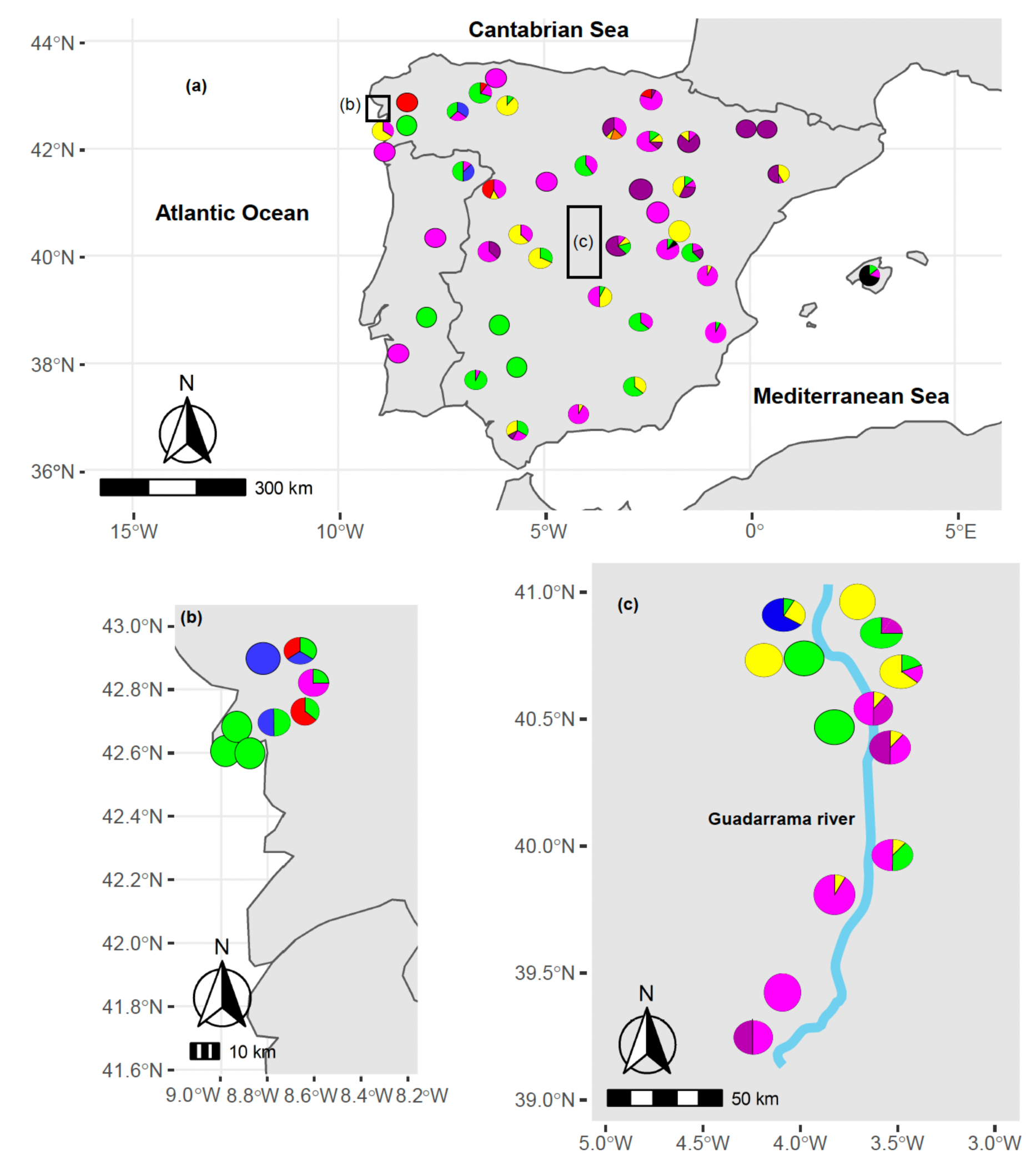
a: Lineages distribution in the Iberian Peninsula (modified from de Sosa et al., 2023a). b: Localities from a lower-scale study in Carnota, A Coruña, Spain (de Sosa et al. 2017). c: Localities from a lower-scale study in Guadarrama river basin, Madrid, Spain (de Sosa et al. 2017). Colors used are the same as in Figure 1.

### 3.3. Genetic diversity and genetic divergence

A total of 485 haplotypes were found among 1,038 sequences for the COI gene, 153 haplotypes within 262 sequences for 16S and 187 haplotypes within 378 sequences for 28S. Values of haplotype and nucleotide diversity of COI for each clade are shown in Table 1. The clade II showed the greatest diversity, while all specimens of clade IV belonged to the same COI haplotype. Haplotype diversity (H) and nucleotide diversity (π) based on COI including all the specimens within the study were 0.97 and 0.059 respectively. H and π for 16S-tRNas were 0.88 and 0.022. Finally, genetic diversity parameters for 28S were 0.33 and 0.003. Moreover, the genetic diversity (H/π) parameters in the three peninsulas studied were: 0.82/0.055 for the Iberian Peninsula, 0.91/0.06 for the Italian Peninsula and 0.65/0.06 for the Scandinavian Peninsula.

**Table 1.**
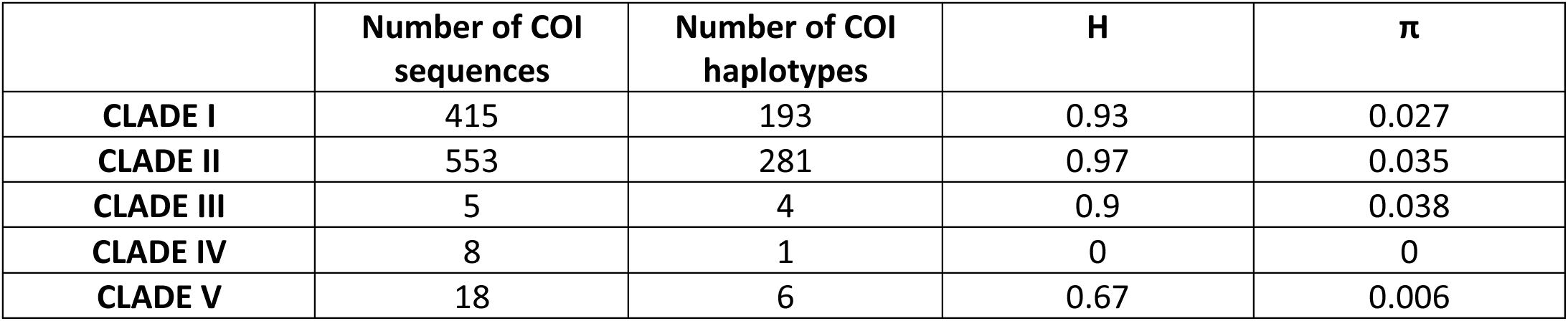
Genetic diversity parameters of each clade based on the COI gene. H: haplotype diversity. π: nucleotide diversity.

| | Number of COI sequences | Number of COI haplotypes | H | $\pi$ |
| --- | --- | --- | --- | --- |
| **CLADE I** | 415 | 193 | 0.93 | 0.027 |
| **CLADE II** | 553 | 281 | 0.97 | 0.035 |
| **CLADE III** | 5 | 4 | 0.9 | 0.038 |
| **CLADE IV** | 8 | 1 | 0 | 0 |
| **CLADE V** | 18 | 6 | 0.67 | 0.006 |

Genetic distances within clades based on COI (Table 2), ranged from 0% to 5% and showed little variability. In contrast, inter-clade distances were higher, 6.55-15.22%. The clade III showed the greatest variability with other clades, even into the ambiguous gap between intraspecific and interspecific divergence in earthworms proposed by Chang and James (2011), 9-15%.

**Table 2.** Percentage of uncorrected pairwise genetic distances based on COI retrieved for *E. tetraedra*.

|  | CLADE I | CLADE II | CLADE III | CLADE IV | CLADE V |
| --- | --- | --- | --- | --- | --- |
| **CLADE I** | 2.39 | 8.57 | 13.8 | 7.47 | 8.22 |
| **CLADE II** |  | 5 | 15.11 | 8.8 | 9.33 |
| **CLADE III** |  |  | 3.79 | 14.77 | 15.22 |
| **CLADE IV** |  |  |  | 0 | 6.55 |
| **CLADE V** |  |  |  |  | 0.58 |

### 3.4. Ecological niche modeling

The ecological niche models obtained for the Clade I and Clade II localities displayed different predictive power, with higher AUC and kappa values (0.71 vs 0.79 and 0.41 vs 0.57 respectively), higher sensitivity and specificity (0.72 vs 0.79 and 0.71 vs 0.80 respectively) and lower omission rates (0.28-0.21) for the Clade II model (Supplementary File 3).

The geographical representation of the predicted suitability values is shown in Figure 4. Highly suitable areas were more widespread for Clade II, covering most of the Mediterranean countries and Britain while being scarce in Scandinavia and other northern countries. For Clade I, highly suitable areas corresponded to countries in the same latitude as Britain and higher, being especially widespread in Scandinavia.

**Figure 4.**
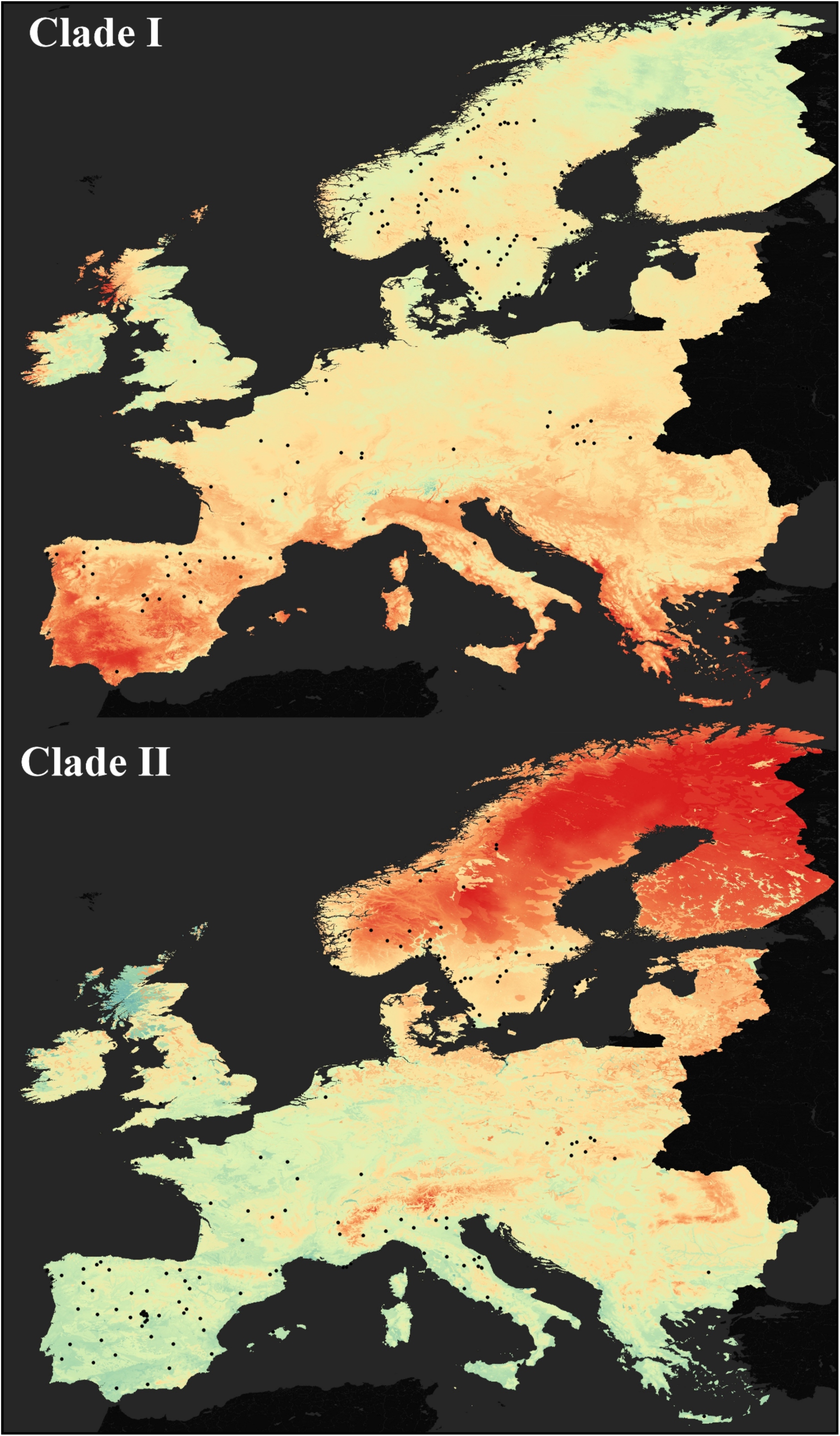
Ecological niche models obtained for the Clade I and Clade II localities (shown as black dots). Red color indicates the lowest estimated suitability and blue-green color indicate the maximum suitability.

The relative contributions of the predictor variables to each model are shown in Supplementary File 3. BIO7, BIO6, BIO13, AWC and CLC were the most influential variables for Clade I, while BIO6, PARMA, AWC and CRUST were the most influential variables for Clade II (Supplementary File 3).

### 3.5. Morphological data

Morphological data is shown in Supplementary File 2. The sexual characters belonging to the male apparatus were the most diverse (e.g. male pore, seminal vesicles and spermathecae) and no presence of sperm was found in any specimen. However, 37 earthworms had spermatophores, but empty. These individuals are nested in the same lineages or even haplotypes with other specimens that do not have spermatophores. Thus, no genetic basis for the presence of this trait can be assumed.

## 4. Discussion

Several genetic studies on earthworms have shown that some morphologically distinct species turn out to be complexes of genetically well individualized lineages (King et al., 2008; Porco et al., 2013). It is still controversial whether there are morphological or ecological differences between these lineages and whether they can be considered as cryptic species (Marchán et al., 2018; Shekhovtsov et al., 2018). Parthenogenetic lineages can arise from sexual species in a variety of ways: bacterial infections such as Wolbachia or Rickettsia (Huigens et al., 2000), spontaneous loss of sex due to mutations in genes related to mating and fertilization of eggs (Carson et al., 1982), or in genes involved in sexual forms (Simon et al., 2003) and contagious origin, with incomplete reproductive isolation between sexual individuals and pre-existing parthenogenetic lineages (Simon et al., 2003) and hybridization origin between individuals of the same or closely related species (Lorenzo-Carballa and Cordero-Rivera, 2009).

The genus *Eiseniella* Michaelsen, 1900 includes two quite distinct groups (Omodeo and Rota, 1991). One includes only the parthenogenetic and cosmopolitan species *Eiseniella tetraedra* and the other includes sexual species such as *Eiseniella lacustris* Cernosvitov, 1931 or *Eiseniella ochridana* Cernosvitov, 1931, most of which occur only in Eastern Europe. This biogeographic pattern may indicate the existence of geographic parthenogenesis (Butlin, 2002), in which sexual forms are restricted to areas around the Mediterranean Sea, while parthenogens spread to the rest of the world, presumably due to a greater capacity for colonization and a high potential for adaptation to new environmental conditions. The recent inclusion of *E. neapolitana* in the genus *Norealidys* (de Sosa et al., 2023b) also highlights the need for phylogenetic studies in this group.

Although several subspecies had been defined for *E. tetraedra* based on the position of the male pores, no genetic basis for this has been found (de Sosa et al., 2017). However, eight different lineages nested in two clades have been identified for this species in the Iberian Peninsula (de Sosa et al., 2023a). In the present study, we found eleven lineages in five clades in Europe. As sexual forms of *E. tetraedra* were not ever found, this high genetic diversity in a parthenogenetic species could be due to an ancestral hybridization origin between individuals of the related species of the genus.

According to European Environmental Agency, eleven biogeographical regions in Europe could be defined: Alpine, Anatolian, Atlantic, Artic, Black Sea, Boreal, Continental, Macaronesian, Mediterranean, Pannonian and Steppic. In this study, we sampled *E. tetraedra* in all regions except four (Black Sea, Anatolian, Macaronesian and Steppic).

Clade I was widespread in Europe. However, it was clearly dominant in cold biogeographical regions such as the Alpine, Arctic, Atlantic, Boreal, Continental and Pannonian regions. To a lesser extent, clade I also occurred in the Mediterranean region, only in Iberian Peninsula. This distribution explains the ecological niche model derived for this clade, which is particularly prevalent in Scandinavian Peninsula. The most influential variables for this clade were also consistent with these results. BIO6 is related to the minimum temperature of the coldest month, which is likely to be low in clade I niches. In contrast, BIO13 corresponds to the precipitation of the wettest month, which is high, as expected. According to de Sosa et al. (2023a), temperature and precipitation were the most important factors affecting the distribution of *E. tetraedra* in Iberian Peninsula. It seems that these variables remain important in Europe as well. Nested in clade I, four linages were found. The most numerous and widespread was lineage B. It clearly dominated in the Scandinavian Peninsula and Pannonian regions, confirming the best adaptation to these zones. In contrast, lineages C, D and G were geographically restricted. Lineage C occurred only in Atlantic regions, mainly in the Iberian Peninsula. The presence of this lineage in United Kingdom, Norway and France may be due to human transport, which has been previously reported in this species (Gates, 1977; Javidkar et al., 2020). Lineage D occurred only at Iberian Peninsula, and mainly in the Atlantic region. We found only a single occurrence in the Mediterranean region, which could also be attributed to human transport. Finally, lineage G occurred in the Atlantic and in one Mediterranean region, although it was scattered in the southern half of Europe (United Kingdom, Netherlands, France and Balearic Islands).

Like clade I, clade II was widespread in Europe and included four lineages. It was predominant in Mediterranean regions, but also occurred in Alpine, Atlantic, Boreal, Continental and Pannonian regions. This dominance explains its distribution in potential niches, which includes all Mediterranean regions and is scarce in the Scandinavian Peninsula and northern countries. Considering its potential distribution, it seems reasonable that the most influential variable for this clade was the minimum temperature of the coldest month. de Sosa et al. (2023a) found that pH was one of the most important factors affecting the distribution of lineages in this species. In agreement with this, lithology also appeared to be an influential variable for the clade II. Lineages A, E, F and H were nested in the clade II. All lineages occurred predominantly in the Mediterranean region in Europe. However, lineage A also occurred in Atlantic, Pannonian and Continental regions; lineage F in Pannonian, Continental (only in France) and to a lesser extent in Boreal and Atlantic regions; lineage E in Atlantic, Continental and Boreal regions. Finally, lineage H, which occurs only in one place on Iberian Peninsula, belonged to the Mediterranean region.

The parthenogenetic and cosmopolitan earthworms *Aporrectodea trapezoides* Dugés, 1828 and *Aporrectodea rosea* Dugés, 1828 appeared to be divided into two distinct clades: one present in the Eurosiberian region and the other in the Mediterranean region (Fernández et al., 2012; Fernández et al., 2015). Although clades I and II of *E. tetraedra* were not restricted to one region; the trend seems similar to those species. While paleographic events seem to be of great importance for the earthworm’s present-day distribution due to its low vagility (Fernández et al., 2013; Novo et al., 2011), the ability of *E. tetraedra* to disperse by hydrochory, possibly also by zoochory (Terhivuo and Saura, 2006) and by anthropochory (Gates, 1977; Javidkar et al., 2020) may contribute to the underlying processes being hidden in a confusing phylogeography.

The predominant presence of the clade III (lineage I) in Eastern Europe, its position on the phylogenetic tree, and its high genetic distance from the other clades of *E. tetraedra* indicate the ancestry of the clade and the possible origin of *E. tetraedra* in Eastern Europe, where most of the sexually related species of the genus were found. This hypothesis needs to be confirmed by future studies such as ancestral territory reconstruction (Fernández et al., 2016). Although all genetic distances between clades were high, only the distances between the clade III and the rest were in the ambiguous gap between intraspecific and interspecific divergence in earthworms proposed by Chang and James (2011), namely 9-15%. Blakemore (1999) described the species delimitation of parthenogenetic earthworms as a “systematic nightmare”. The biological term “species” is not applicable to parthenogenetic earthworms because of the reproductive isolation of each individual. However, the authors tried some species delimitation rules (as GMYC) with no conclusive results.

Clades IV and V included a minor number of individuals of *E. tetraedra*, mostly restricted to the Scandinavian Peninsula, although they had punctual presences in France and Italy (only one specimen in each country) probably due to human transport.

The model of glacial refugia as core areas for the survival of thermophilic and/or temperate animal and plant species during unfavourable Pleistocene environmental conditions and as sources of postglacial recolonization processes is widely accepted in biogeography (Hewitt, 2000; Willis and Whittaker, 2000). It is generally believed that the main hypothesis of the recolonization of Europe after the LGM for earthworms is the “*tabula rasa*” hypothesis. It states that earthworms became extinct in northern latitudes and recolonized these areas after the ice melted. However, there are examples in earthworms, such as *Dendrobaena octaedra*, that did not follow this pattern (Hansen et al., 2006). The nunatak hypothesis suggests that there were some ice-free refuges in northern Europe, such as mountains rising above the ice sheet or ice-free refuges on the coast, where the biota could survive. According to our results, *E. tetraedra* could follow both patterns. Although the high haplotype diversity of Italian Peninsula, 0.96, could be due to its role as a refuge during the LGM, the number of individuals from Italy was lower than on other sampled peninsulas. De Sosa et al. (2023a) found lower haplotype diversity than de Sosa et al. (2017) in the Iberian Peninsula, which expands the sampling locations. Therefore, we think that this high haplotype diversity could be explained by the smaller number of samples. The presence of restricted lineages at Iberian Peninsula and even clades at Scandinavian Peninsula and Eastern Europe suggest that the three areas served as refugia during the LGM, bringing *E. tetraedra* to the rest of Europe after the ice melted. The Iberian Peninsula was one of the most important glacial refugia of the Pleistocene in Europe (Hewitt, 1999; Hewitt, 2001) and served as a species repository for northern countries (Beebee and Rowe, 2000; Vernesi et al., 2002). Eastern Europe was also an important refuge during the LGM (Sommer and Zachos, 2009) and biota, even earthworms, survived in some ice-free refugia in northern Europe (Hansen et al., 2006).

## 5. Conclusions

*Eiseniella tetraedra* has been found to have a high genetic diversity in Europe. This diversity was classified into eleven lineages nested in five clades. Clades I and II were widely distributed in Europe, while the others had a limited distribution. Clade I was more represented in cold biogeographical regions such as the continent, the Atlantic or even the Arctic, while clade II was prevalent in Mediterranean regions. Potential niches were also consistent with distribution trends. This is consistent with the phylogeographic patterns of other cosmopolitan and parthenogenetic earthworms. The clade III is largely restricted to Eastern Europe and appears to be the original clade. Clades IV and V were mostly present in Scandinavian Peninsula. The presence of restricted clades in the Iberian and Scandinavian Peninsula and Eastern Europe, suggests that the three acted as refugia during the LGM. Thus, both the “*tabula rasa*” and nunatak hypotheses could apply to *E. tetraedra* in Europe.

## Acknowledgments

We would like to thank Nuria Sánchez for field support. IS was supported by a Predoctoral Fellowship grant by Complutense University of Madrid. DF was supported by a Juan de la Cierva Formación grant from Spanish Government (FJCI-2017-32895) and a MOPGA grant from the French Government (mopga-postdoc-3--6111272103), and MN was supported by Ramón y Cajal Fellowship (RYC2018-024654-I) of the Spanish Government. This research was funded by project CGL2013-42908-P from the Spanish Government. Several Scandinavian specimens were collected by Christer Erséus and coworkers during faunal inventories funded by the Swedish Taxonomy Initiative, Swedish Species Information Centre (SLU ArtDatabanken), and the Norwegian Taxonomy Initiative, Norwegian Biodiversity Information Centre (NTNU Artsdatabanken).

## Data Availability Statement

The data that support the findings of this study are openly available in GenBank at https://www.ncbi.nlm.nih.gov/genbank/. Reference numbers are available in Supplementary Table 6.

## References

Beebee, T. J., & Rowe, G. (2000). Microsatellite analysis of natterjack toad Bufo calamita Laurenti populations: consequences of dispersal from a Pleistocene refugium. Biological Journal of the Linnean Society, 69(3), 367–381.

Blakemore, R. J. (1999). Diversity of exotic earthworms in Australia–a status report. Proceedings of “The other, 99, 182–187.

Blakemore, R. J. (2007). Checklist of megadrile earthworms from Greenland and Iceland. Yokohama: Yokohama National University, Japan.

Blakemore, R.J. (2006). Cosmopolitan Earthworms-an Eco-taxonomic Guide to the Peregrine Species of the World (2nd ed.). VermEcology, Australia

Butlin, R. (2002). The costs and benefits of sex: new insights from old asexual lineages. Nature Reviews Genetics, 3(4), 311–317.

Carson, H. L., Chang, L. S., & Lyttle, T. W. (1982). Decay of female sexual behavior under parthenogenesis. Science, 218(4567), 68–70.

Casellato, S. 1987. On polyploidy in Oligochaetes with particular reference to Lumbricids. In On Earthworms. Modena: Selected symposia and monographs UZI, 75–87.

Chang, C. H., James, S., 2011. A critique of earthworm molecular phylogenetics. Pedobiología, 54, S3–S9.

Crawford, P. H., & Hoagland, B. W. (2010). Using species distribution models to guide conservation at the state level: the endangered American burying beetle (Nicrophorus americanus) in Oklahoma. Journal of Insect Conservation, 14(5), 511–521.

Csuzdi, C. (2012). Earthworm species, a searchable database. Opuscula Zoologica Budapest, 43(1), 97–99.

Darriba, D., Taboada, G.L., Doallo, R., Posada, D., 2012. jModelTest 2: more models, new heuristics and parallel computing. Nature Methods, 9, 772–772.

De Sosa, I., Marchan, D. F., Novo, M., Almodovar, A., Díaz Cosín, D. J., 2017a. Bless this phylogeographic mess–Comparative study of Eiseniella tetraedra (Annelida, Oligochaeta) between an Atlantic area and a continental Mediterranean area in Spain. European Journal of Soil Biology, 78, 50–56.

de Sosa, I., Verdes, A., Tilikj, N., Marchán, D. F., Planelló, R., Herrero, Ó., … & Novo, M. (2022). How to thrive in unstable environments: Gene expression profile of a riparian earthworm under abiotic stress. Science of the Total Environment, 817, 152749.

de Sosa, I., Marchán, D. F., Novo, M., Almodóvar, A., & Diaz Cosin, D. J. (2023a). Phylogeography of a riparian earthworm shows environmental factors influence genetic structure. Journal of Biogeography, 50(1), 156–168.

de Sosa, I., F. Marchán, D., Novo, M., Szederjesi, T., Jelic, M., Jabłońska, A., … & Díaz Cosín, D. J. (2023b). Guess who? Taxonomic problems in the genus Eiseniella revisited by integrated approach. Organisms Diversity & Evolution, 23(2), 295–308.

Decaëns, T. (2010). Macroecological patterns in soil communities. Global Ecology and Biogeography, 19(3), 287–302.

Domínguez, J., Aira, M., Breinholt, J. W., Stojanovic, M., James, S. W., & Pérez-Losada, M. (2015). Underground evolution: new roots for the old tree of lumbricid earthworms. Molecular phylogenetics and evolution, 83, 7–19.

Edwards, C. A. (2004). The importance of earthworms as key representatives of the soil fauna. Earthworm ecology, 2, 3–11.

Fernández, R., Almodóvar, A., Novo, M., Simancas, B., Díaz Cosín, D. J., 2012. Adding complexity to the complex: new insights into the phylogeny, diversification and origin of parthenogenesis in the Aporrectodea caliginosa species complex (Oligochaeta, Lumbricidae). Molecular phylogenetics and evolution, 64, 368–379.

Fernández, R., Almodóvar, A., Novo, M., Gutiérrez, M., Díaz Cosín, D. J., 2013. Earthworms, good indicators for palaeogeographical studies? Testing the genetic structure and demographic history in the peregrine earthworm Aporrectodea trapezoides (Dugès, 1828) in southern Europe. Soil Biology and Biochemistry, 58, 127–135.

Fernández, R., Novo, M., Marchán, D. F., Díaz Cosín, D. J., 2015. Diversification patterns in cosmopolitan earthworms: similar mode but different tempo. Molecular phylogenetics and evolution, 94, 701–708.

Fernández, R., Novo, M., Marchán, D. F., & Cosín, D. J. D. (2016). Diversification patterns in cosmopolitan earthworms: similar mode but different tempo. Molecular phylogenetics and evolution, 94, 701–708.

Fridolin, V. Yu. (1936): Animal and plant communities of the Khibiny mountains I. – Proceedings of the Kola Station AS USSR 3: 19–295

Gates, G.E., 1977. Contribution to a revision of the earthworm family Lumbricidae. XX. The genus Eiseniella in North America. Megadrilogica, 3, 71–79.

Hall, T. A., 1999. BioEdit: a user-friendlybiological sequence alignment editor and analysis program for Windows 95/98/NT. Nucleic Acids Symposium, 41, 95–98.

Hansen, P. L., Holmstrup, M., Bayley, M., & Simonsen, V. (2006). Low genetic variation for Dendrobaena octaedra from Greenland compared to populations from Europe and North America: Refuge or selection?. Pedobiologia, 50(3), 225–234.

Haraldsen, T. K. & Engelstad, F. (1998): Influence of earthworms on soil properties and crop production in Norway. – Centre for Soil and Environmental Research, Oslo, 12 pp

Hewitt, G. M. (1999). Post-glacial re-colonization of European biota. Biological journal of the Linnean Society, 68(1-2), 87–112.

Hewitt GM (2000) The genetic legacy of the Quaternary ice ages. Nature, 405, 907– 913.

Hewitt, G. M. (2001). Speciation, hybrid zones and phylogeography—or seeing genes in space and time. Molecular ecology, 10(3), 537–549.

Holmstrup, M., Ostergaard, I.K., Nielsen, A. & Hansen, B.T. (1991) The relationship between temperature and cocoon incubation time for some lumbricid earthworm species. Pedobiologia, 35, 179– 184.

Huigens, M. E., Luck, R. F., Klaassen, R. H. G., Maas, M. F. P. M., Timmermans, M. J. T. N., & Stouthamer, R. (2000). Infectious parthenogenesis. Nature, 405(6783), 178–179.

Javidkar, M., Abdoli, A., Ahmadzadeh, F., Nahavandi, Z., Yari, M., 2021. Molecular evidence reveals introduced populations of Eiseniella tetraedra (Savigny, 1826)(Annelida, Lumbricidae) with European origins from protected freshwater ecosystems of the southern Alborz Mountains. Marine and Freshwater Research, 72, 44–57.

Julin, E. (1950). *De svenska daggmaskarterna* (pp. 58-pp).

Katoh, K., Standley, D. M., 2013. MAFFT multiple sequence alignment software version 7: Improvements in performance and usability. Molecular biology and evolution, 30, 772–780.

King, R. A., Tibble, A. L., Symondson, W. O., 2008. Opening a can of worms: unprecedented sympatric cryptic diversity within British lumbricid earthworms. Molecular ecology, 17, 4684–4698.

Lorenzo-Carballa, M. O., & Cordero-Rivera, A. (2009). Thelytokous parthenogenesis in the damselfly Ischnura hastata (Odonata, Coenagrionidae): genetic mechanisms and lack of bacterial infection. Heredity, 103(5), 377–384.

Marchán, D. F., Refoyo, P., Novo, M., Fernández, R., Trigo, D., & Cosín, D. J. D. (2015). Predicting soil micro-variables and the distribution of an endogeic earthworm species through a model based on large-scale variables. Soil Biology and Biochemistry, 81, 124–127.

Marchán, D. F., Refoyo, P., Fernández, R., Novo, M., de Sosa, I., & Cosín, D. J. D. (2016). Macroecological inferences on soil fauna through comparative niche modeling: the case of Hormogastridae (Annelida, Oligochaeta). European Journal of Soil Biology, 75, 115–122.

Marchán, D. F., Fernández, R., de Sosa, I., Sánchez, N., Cosín, D. J. D., & Novo, M. (2018). Integrative systematic revision of a Mediterranean earthworm family: Hormogastridae (Annelida, Oligochaeta). Invertebrate Systematics, 32(3), 652–671.

Marek, P. E., Shear, W. A., & Bond, J. E. (2012). A redescription of the leggiest animal, the millipede Illacme plenipes, with notes on its natural history and biogeography (Diplopoda, Siphonophorida, Siphonorhinidae). ZooKeys, (241), 77.

Maynard, D. S., Crowther, T. W., King, J. R., Warren, R. J., & Bradford, M. A. (2015). Temperate forest termites: ecology, biogeography, and ecosystem impacts. Ecological Entomology, 40(3), 199–210.

Miller, M.A., Pfeiffer, W., Schwartz, T., 2010. Creating the CIPRES science gateway for inference of large phylogenetic trees. Proceedings of the Gateway Computing Environments Workshop (GCE), 1–8.

Morariu, V. I., Srinivasan, B. V., Raykar, V. C., Duraiswami, R., Davis, L. S., 2008. Automatic online tuning for fast Gaussian summation. Advances in Neural lnformation Processing Systems {NlPS}.

Novo, M., Almodóvar, A., Fernández, R., Giribet, G., & Cosín, D. J. D. (2011). Understanding the biogeography of a group of earthworms in the Mediterranean basin—The phylogenetic puzzle of Hormogastridae (Clitellata: Oligochaeta). Molecular Phylogenetics and Evolution, 61(1), 125–135.

Omodeo, P., Rota, E., 1991. Earthworms of Turkey. II. Italian Journal of Zoology, 58, 171–181.

Porco, D., Decaëns, T., Deharveng, L., James, S. W., Skarżyński, D., Erséus, C., …, Hebert, P. D., 2013. Biological invasions in soil: DNA barcoding as a monitoring tool in a multiple taxa survey targeting European earthworms and springtails in North America. Biological Invasions, 15, 899–910.

Ronquist, F., Huelsenbeck, J.P., 2003. MRBAYES 3: Bayesian phylogenetic inference under mixed models. Bioinformatics, 19, 1572–1574.

S.J. Phillips, R.P. Anderson, R.E. Schapire. Maximum entropy modeling of species geographic distributions. Ecol. Model, 190 (2006), pp. 231–259.

Schmitt, S., Pouteau, R., Justeau, D., De Boissieu, F., & Birnbaum, P. (2017). ssdm: An r package to predict distribution of species richness and composition based on stacked species distribution models. Methods in Ecology and Evolution, 8(12), 1795–1803.

Shekhovtsov, S. V., Berman, D. I., Bulakhova, N. A., Vinokurov, N. N., & Peltek, S. E. (2018). Phylogeography of Eisenia nordenskioldi nordenskioldi (Lumbricidae, Oligochaeta) from the north of Asia. Polar Biology, 41(2), 237–247.

Shekhovtsov, S. V., Rapoport, I. B., Poluboyarova, T. V., Geraskina, A. P., Golovanova, E. V., & Peltek, S. E. (2020). Morphotypes and genetic diversity of Dendrobaena schmidti (Lumbricidae, Annelida). Vavilov Journal of Genetics and Breeding, 24(1), 48.

Simon, J. C., Delmotte, F., Rispe, C., & Crease, T. (2003). Phylogenetic relationships between parthenogens and their sexual relatives: the possible routes to parthenogenesis in animals. Biological Journal of the Linnean Society, 79(1), 151–163.

Si-Moussi, S. (2010). Apports de la fouille de données à la modélisation des communautés écologiques.

Sommer, R. S., & Zachos, F. E. (2009). Fossil evidence and phylogeography of temperate species:‘glacial refugia’and post-glacial recolonization.

Stamatakis, A., 2006. RAxML-V1-HPC: maximum likelihood-based phylogenetic analyses with thousands of taxa and mixed models. Bioinformatics, 22, 2688–2690.

Stöp-Bowitz, C. (1969): A contribution to our knowledge of the systematics and zoogeography of Norwegian earthworms (Annelida, Oligochaeta: Lumbricidae). – Nytt Magasin for Zoologi 17: 169–280.

Terhivuo, J. (1988, January). The Finnish Lumbricidae (Oligochaeta) fauna and its formation. In Annales Zoologici Fennici (pp. 229–247). Finnish Academy of Sciences, Societas Scientiarum Fennica, Societas pro Fauna et Flora Fennica and Societas Biologica Fennica Vanamo.

Terhivuo, J., Saura, A. 1997. Island biogeography of North European parthenogenetic Lumbricidae: I. Clone pool affinities and morphometric differentiation of Åland populations. Ecography 20, 185–196. 10.1111/j.1600-0587.1997.tb00361.x.

Terhivuo, J., Saura, A., (2006). Dispersal and clonal diversity of North-European parthenogenetic earthworms. In Biological Invasions Belowground: Earthworms as Invasive Species (pp. 5–18). Springer, Dordrecht.

Tiunov, A. V., Hale, C. M., Holdsworth, A. R., & Vsevolodova-Perel, T. S. (2006). Invasion patterns of Lumbricidae into the previously earthworm-free areas of northeastern Europe and the western Great Lakes region of North America. In Biological Invasions Belowground: Earthworms as Invasive Species (pp. 23–34). Springer, Dordrecht.

Vernesi, C., Pecchioli, E., Caramelli, D., Tiedemann, R., Randi, E., & Bertorelle, G. (2002). The genetic structure of natural and reintroduced roe deer (Capreolus capreolus) populations in the Alps and central Italy, with reference to the mitochondrial DNA phylogeography of Europe. Molecular Ecology, 11(8), 1285–1297.

Willis, K. J., & Whittaker, R. J. (2000). The refugial debate. Science, 287(5457), 1406–1407.

