## Supplementary Material for "The nunatak and *tabula rasa* hypotheses may be compatible: the European phylogeography of a riparian earthworm"

| Locality | Longitude | Latitude | Sampling |
|---|---|---|---|
| **Burgohondo** | -4,7877 | 40,4139 | De Sosa et al. 2022 |
| **Soria** | -2,8566 | 41,7273 | De Sosa et al. 2022 |
| **Ordesa** | 0,0784 | 42,6381 | De Sosa et al. 2022 |
| **Posets-Maladeta** | 0,5282 | 42,6522 | De Sosa et al. 2022 |
| **Alba de Tormes** | -5,5179 | 40,8268 | De Sosa et al. 2022 |
| **Sanjuanejo** | -6,4928 | 40,5683 | De Sosa et al. 2022 |
| **Portalegre** | -7,9096 | 39,0246 | De Sosa et al. 2022 |
| **Casas de Don Antonio** | -6,2904 | 39,2339 | De Sosa et al. 2022 |
| **Viseu** | -7,4949 | 40,8478 | De Sosa et al. 2022 |
| **Alcacer do Sal** | -8,3372 | 38,2635 | De Sosa et al. 2022 |
| **Jabugo** | -6,7001 | 38,0791 | De Sosa et al. 2022 |
| **Ziordia** | -2,2303 | 42,87 | De Sosa et al. 2022 |
| **Miranda del Ebro** | -2,9282 | 42,6885 | De Sosa et al. 2022 |
| **Olmedillo de Roa** | -3,9333 | 41,7822 | De Sosa et al. 2022 |
| **Valladolid** | -4,7373 | 41,6674 | De Sosa et al. 2022 |
| **Río Cauxa** | -6,3229 | 43,2797 | De Sosa et al. 2022 |
| **El Castril** | -2,7769 | 37,7899 | De Sosa et al. 2022 |
| **Urda** | -3,7116 | 39,4165 | De Sosa et al. 2022 |
| **Arenas del Rey** | -3,8968 | 36,9572 | De Sosa et al. 2022 |
| **Peraleda del Zaucejo** | -5,5664 | 38,4714 | De Sosa et al. 2022 |
| **Chelva** | -0,9975 | 39,7582 | De Sosa et al. 2022 |
| **Tarazona** | -1,7275 | 41,9023 | De Sosa et al. 2022 |
| **Poveda de la Sierra** | -2,0283 | 40,6412 | De Sosa et al. 2022 |
| **El Palau d'Anglesola** | 0,8994 | 41,6566 | De Sosa et al. 2022 |
| **Tibi** | -0,5874 | 38,5283 | De Sosa et al. 2022 |
| **Arcos de Jalón** | -2,2737 | 41,2158 | De Sosa et al. 2022 |
| **Sos del Rey Católico** | -1,2194 | 42,4768 | De Sosa et al. 2022 |
| **San Blas** | -1,1774 | 40,3582 | De Sosa et al. 2022 |
| **Calahorra** | -1,9674 | 42,3002 | De Sosa et al. 2022 |
| **Molina de Aragón** | -1,8894 | 40,8427 | De Sosa et al. 2022 |
| **Sotuélamos** | -2,5723 | 39,0408 | De Sosa et al. 2022 |
| **Esporlas** | 2,5795 | 39,6657 | De Sosa et al. 2022 |
| **Bragança** | -6,754 | 41,8045 | De Sosa et al. 2022 |
| **Mallo de Luna** | -5,8873 | 42,8721 | De Sosa et al. 2022 |
| **Puenticella** | -6,5265 | 43,1383 | De Sosa et al. 2022 |
| **El Bosque** | -5,5034 | 36,7622 | De Sosa et al. 2022 |
| **Miñarzos** | -9,133 | 42,7992 | De Sosa et al. 2017 |
| **Portocubelo** | -9,1355 | 42,8041 | De Sosa et al. 2017 |
| **Mallou** | -9,099 | 42,8072 | De Sosa et al. 2017 |
| **Lavadero de Lira** | -9,1345 | 42,8001 | De Sosa et al. 2017 |
| **Locality** | **Longitude** | **Latitude** | **Sampling** |
| **Fuente de Cornido** | -9,0908 | 42,8682 | De Sosa et al. 2017 |

| | | | |
|---|---|---|---|
| **Ocruceiro** | -9,082 | 42,8515 | De Sosa et al. 2017 |
| **Cornido** | -9,0908 | 42,8682 | De Sosa et al. 2017 |
| **Chaguazoso** | -7,2164 | 42,192 | De Sosa et al. 2017 |
| **Arroyo Playa Cons** | -9,108 | 42,8072 | De Sosa et al. 2017 |
| **Forcarei** | -8,3442 | 42,5831 | De Sosa et al. 2017 |
| **Montalvo** | -8,8463 | 42,3994 | De Sosa et al. 2017 |
| **Lavadero de Oia** | -8,8726 | 42,0031 | De Sosa et al. 2017 |
| **Lavadero de Casalonga** | -8,6119 | 42,8217 | De Sosa et al. 2017 |
| **Souto** | -7,2151 | 42,9182 | De Sosa et al. 2017 |
| **Fuente de las Hondillas** | -4,1396 | 40,6974 | De Sosa et al. 2017 |
| **Alcalá de Henares** | -3,308 | 40,5111 | De Sosa et al. 2017 |
| **Las Rozas** | -3,9401 | 40,5176 | De Sosa et al. 2017 |
| **Villanueva del Pardillo** | -3,9365 | 40,4856 | De Sosa et al. 2017 |
| **Valdemorillo** | -4,0947 | 40,5159 | De Sosa et al. 2017 |
| **Río Hornillo** | -4,2295 | 40,5957 | De Sosa et al. 2017 |
| **El Soto** | no data | no data | De Sosa et al. 2017 |
| **Arroyo de los Irrios** | -4,0858 | 40,7177 | De Sosa et al. 2017 |
| **Los Molinos** | -4,0858 | 40,7177 | De Sosa et al. 2017 |
| **Parquelagos** | -3,969 | 40,5993 | De Sosa et al. 2017 |
| **Arroyomolinos** | -3,9299 | 40,2778 | De Sosa et al. 2017 |
| **Fuente de Guadarrama** | -4,1048 | 40,6732 | De Sosa et al. 2017 |
| **Rielves** | -4,1902 | 39,9634 | De Sosa et al. 2017 |
| **Casarrubios del Monte** | -4,0286 | 40,1832 | De Sosa et al. 2017 |
| **Albarreal del Tajo** | -4,1839 | 39,8984 | De Sosa et al. 2017 |

Supplementary Table 1. Localities sampled in the Iberian Peninsula.

| Locality | Longitude | Latitude |
|---|---|---|
| **Napoli** | 14,3306 | 40,8849 |
| **Caorso-delta roncaglia** | 9,8526 | 45,04356 |
| **Masio** | 8,41 | 44,8712 |
| **Pisa** | 10,3892 | 43,729 |
| **Verona** | 10,9992 | 45,4339 |
| **Roma** | 12,4657 | 41,9012 |
| **Via Boalto a Levante** | 11,545 | 45,0388 |
| **Lusserna** | 7,2433 | 44,6444 |
| **Pontecorvo** | 13,6652 | 41,4542 |
| **Milan** | 9,2006 | 45,4739 |
| **Vicenza** | 11,5511 | 45,5471 |
| **Valdarno** | 11,5606 | 43,5499 |
| **Ferriera** | 12,4508 | 43,0904 |
| **Jesi** | 13,2291 | 43,5134 |
| **San Lorenzo** | 13,1845 | 43,0538 |
| **Conca Cave** | 15,2733 | 37,9567 |
| **Amandola** | 13,3574 | 42,9842 |
| **Antillo** | 13,3674 | 42,966 |
| **Affluent of Fiume Corno, near Norcia** | 13,0797 | 42,7949 |
| **near Genga Railway Station, Genga (Ancona), Marche, Italy** | 12,9773 | 43,4029 |

Supplementary Table 2. Localities sampled in Italy by Irene de Sosa, Christer Érseus and co-workers (blue) and Misel Jelic (purple).

| Locality | Longitude | Latitude |
|---|---|---|
| **EW426** | 11,2258 | 58,8999 |
| **EW336** | 11,9602 | 57,6873 |
| **Jättadalen Nature Reserve** | 13,6986 | 58,4325 |
| **ew455** | 11,9551 | 57,6822 |
| **EW338** | 12,2706 | 57,7614 |
| **NOEW26** | 5,7543 | 58,5383 |
| **NOEW27** | 5,9183 | 58,5021 |
| **EW352** | 18,4223 | 57,7752 |
| **EW354** | 18,16 | 57,314 |
| **NOEW48** | 12,0539 | 61,5706 |
| **NOEW55** | 11,2264 | 61,5678 |
| **NOEW56** | 12,2002 | 61,3025 |
| **EW343** | 12,6142 | 59,7612 |

| | | |
|---|---|---|
| **NOEW70** | 8,2084 | 60,5307 |
| **NOEW74** | 7,6466 | 60,4119 |
| **NOEW81** | 6,1986 | 60,6388 |
| **NOEW86** | 7,29949 | 61,40259 |
| **Locality** | **Longitude** | **Latitude** |
| **NOEW97** | 7,143 | 62,1949 |
| **NOEW110** | 7,1079 | 60,8604 |
| **EW363** | 12,012 | 57,7439 |
| **EW365** | 12,79829 | 57,92529 |
| **EW364** | 12,2251 | 57,7702 |
| **EW369** | 12799 | 57931 |
| **EW458** | 11,9604 | 57,6805 |
| **EW373** | 14,7193 | 61,3272 |
| **EW376** | 17,1118 | 61,72969 |
| **EW379** | 17,0481 | 61,3001 |
| **NOEW131** | 9,1798 | 59,5788 |
| **NOEW135** | 8,486 | 59,8724 |
| **NOEW136** | 8,2297 | 59,7712 |
| **NOEW138** | 8,0122 | 59,4455 |
| **NOEW141** | 6,5263 | 59,9079 |
| **EW388** | 15,80799 | 59,132 |
| **NOEW161** | 10,2189 | 61,4274 |
| **NOEW165** | 7,7058 | 62,5667 |
| **NOEW167** | 8,5508 | 62,8917 |
| **NOEW169** | 10,9794 | 63,4639 |
| **NOEW170** | 12,1039 | 64,2318 |
| **NOEW171** | 12,91807 | 64,79248 |
| **NOEW172** | 13,4041 | 65,53308 |
| **NOEW174** | 13,703 | 66,05419 |
| **NOEW182** | 17,2609 | 68,53631 |
| **NOEW183** | 16,5625 | 68,6303 |
| **NOEW186** | 15,9844 | 69,1223 |
| **NOEW192** | 15,2939 | 67,2656 |
| **NOEW194** | 14,9989 | 67,0747 |
| **NOEW198** | 13,245 | 65,8025 |
| **NOEW200** | 13,1569 | 64,9264 |
| **NOEW214** | 10,0739 | 60,8231 |
| **EW431** | 11,9548 | 57,6796 |
| **NOEW221** | 10,7059 | 59,9281 |

| | | |
|---|---|---|
| **EW412** | 17,9874 | 59,4306 |
| **Kvillebäcken Stream at Hökälla-Ekehöjd** | 11,943 | 57,7563 |
| **EW461** | 12,5007 | 57,9332 |
| **EW463** | 13,2459 | 56,0359 |
| **EW464** | 15,282 | 56,199 |
| **EW465** | 15,105 | 56,179 |
| **EW466** | 12,4257 | 62,6453 |
| **NOEW247** | 11,8164 | 61,6568 |
| **NOEW253** | 11,521 | 58,9099 |
| **Locality** | **Longitude** | **Latitude** |
| **NOEW254** | 11,4359 | 59,0859 |
| **NOEW258** | 10,6956 | 59,542 |
| **EW475** | 16,6074 | 57,6593 |
| **EW433** | 11752 | 57732 |
| **NOEW268** | 10,016 | 59,043 |
| **NOEW278** | 6,3153 | 60,1295 |
| **NOEW288** | 8,7287 | 61,1488 |
| **EW 1** | 11,8452 | 57,6297 |
| **NOEW298** | 25,7633 | 71,1447 |
| **NOEW301** | 24,0789 | 70,2332 |
| **NOEW302** | 23,3 | 69,95 |
| **NOEW304** | 22,0121 | 69,8409 |
| **NOEW312** | 17,98608 | 69,6395 |
| **NOEW183** | 16,5625 | 68,6303 |
| **NOEW158** | 11,2569 | 60,5628 |
| **NOEW160** | 10,4783 | 61,12 |
| **NOEW328** | 9,704 | 62,5918 |
| **NOEW329** | 12,6195 | 64,6366 |
| **NOEW332** | 14,4343 | 67,2784 |
| **NOEW343** | 10,1945 | 62,9701 |
| **EW487** | 11,9591 | 57,6807 |
| **EW494** | 14,27 | 55,5321 |
| **EW496** | 12,49704 | 58,93791 |
| **EW501** | 12,4473 | 59,0002 |
| **EW502** | 12,2338 | 59,1386 |
| **EW4** | 16,0425 | 59,0853 |
| **EW503** | 15,0186 | 59,0398 |
| **NOEW351** | 10,06376 | 60,47608 |
| **NOEW358** | 6,6094 | 61,8464 |

| | | |
|---|---|---|
| **NOEW366** | 9,6369 | 60,3704 |
| **EW499** | 12,5737 | 58,8211 |
| **EW437** | 16,8764 | 56,9855 |
| **EW38** | 15,1431 | 59,2664 |
| **EW439** | 16,8539 | 56,8621 |
| **EW440** | 16,66111 | 56,8195 |
| **EW441** | 16,6468 | 56,8056 |
| **EW442** | 16,8764 | 56,9855 |
| **EW5** | 16,0872 | 59,0875 |
| **EW4** | 16,0425 | 59,0853 |
| **EW445** | 12,241 | 57,776 |
| **EW 15** | 14,7456 | 56,2997 |
| **EW 14** | 14,5028 | 56,1058 |
| **EW 29** | 14,3127 | 58,0372 |
| **Locality** | **Longitude** | **Latitude** |
| **EW 34** | 14,6339 | 58,31 |
| **EW 30** | 14,3084 | 58,0414 |
| **EW 37** | 14,7958 | 58,7306 |
| **EW 38** | 15,1431 | 59,2664 |
| **EW 47** | 14,5564 | 62,4525 |
| **EW51** | 14,4806 | 62,9831 |
| **EW 52** | 13,87 | 62,8506 |
| **EW 55** | 12,9486 | 62,2472 |
| **EW 59** | 14,1792 | 59,1169 |
| **ew64** | 11,6154 | 58,3695 |
| **EW 100** | 11,5744 | 58,4327 |
| **EW 95** | 11,1275 | 58,8903 |
| **EW 105** | 13,1247 | 57,59 |
| **EW 106** | 13,1233 | 57,5928 |
| **EW109** | 11,983 | 57,5 |
| **EW 115** | 11,9542 | 57,6831 |
| **EW 116** | 11,9567 | 57,6817 |
| **EW 124** | 12,9353 | 56,3887 |
| **EW 126** | 12,96 | 55,4797 |
| **EW 130** | 13,0752 | 56,3136 |
| **EW 133** | 12,7978 | 57,9253 |
| **EW 136** | 18,1603 | 57,3139 |
| **EW 140** | 12,453 | 57,993 |
| **EW 144** | 18,2743 | 59,496 |

| | | |
|---|---|---|
| **EW 151** | 17,6282 | 59,8515 |
| **EW419** | 18,4046 | 57,7402 |
| **EW 159** | 18,2742 | 59,4973 |
| **EW 162** | 18,5502 | 59,571 |
| **EW 164** | 16,7598 | 58,6298 |
| **EW420** | 18,5988 | 57,8409 |
| **EW213** | 11,969 | 57,683 |
| **EW 223** | 17,4782 | 62,5235 |
| Locality | **Longitude** | **Latitude** |
| **EW 225** | 17,8746 | 62,8919 |
| **EW 227** | 18,4314 | 62,8797 |
| **EW 229** | 19,6624 | 63,6038 |
| **EW 265** | 15,1348 | 65,0873 |
| **EW 270** | 14,4652 | 65,0947 |
| **EW 271** | 14,2979 | 65,0176 |
| **EW424** | 12,2845 | 57,7736 |
| **EW 303** | 18,4103 | 57,32 |
| **EW 307** | 18,285 | 57,635 |
| **EW 310** | 18,8163 | 57,8507 |
| **Locality** | **Longitude** | **Latitude** |
| **EW 317** | 11,4408 | 59,0078 |
| **EW 318** | 11,1931 | 58,9453 |
| **EW 319** | 11,2261 | 58,9003 |
| **EW 324** | 12,285 | 57,774 |
| **EW 248** | 19,0383 | 68,3484 |
| **EW 257** | 16,1578 | 65,9594 |
| **EW 263** | 16,0426 | 65,2151 |
| **EW 267** | 14,6968 | 65,0635 |
| **EW 269** | 14,5219 | 65,1194 |
| **EW 273** | 14,1278 | 64,8268 |
| **EW 274** | 14,1218 | 64,619 |
| **EW 278** | 13,2783 | 63,3153 |
| **EW 290** | 14,2116 | 58,9656 |
| **EW 284** | 16,6082 | 56,5443 |
| **EW 293** | 11,5423 | 58,0225 |
| **EW 297** | 14,1823 | 57,7529 |
| **EW332** | 16,9983 | 59,5103 |
| **Lake Alvajärvi, Jyväskylä** | 25,7109 | 62,3186 |
| **EW 187** | 15,2247 | 56,7827 |

| | | |
|---|---|---|
| **EW 191** | 11,9562 | 57,6813 |
| **NOEW54** | 10,7676 | 62,1722 |
| **NOEW111** | 7,5585 | 60,8073 |

Supplementary Table 3. Localities sampled in the Scandinavian Peninsula by Christer Érseus and co-workers.

| Locality | Longitude | Latitude |
|---|---|---|
| Brest | No data | No data |
| **FR15829.2.1, Francia** | 2,5489 | 45,5854 |
| **FR15829.2.1, Francia** | 2,5489 | 45,5854 |
| **FR14807.1.1, Francia** | 1,2914 | 45,9135 |
| **FR15808 1.1, Francia** | 5,9462 | 45,3158 |
| **FR14812.1.1, Francia** | 2,42923 | 42,6314 |
| **FR15L31.1.1, Francia** | 2,4292 | 42,6315 |
| **FR15507.1a.1, Francia** | 3,8379 | 47,5797 |
| **FR14824.1B.1, Francia** | 3,2109 | 45,9402 |
| **FR15507.1c.1, Francia** | 3,8379 | 47,5798 |
| **FR15L28.1.1, Francia** | 7,1557 | 48,0433 |
| **FR14824.1b.1, Francia** | 3,2109 | 45,9402 |
| **FR15825.1.1, Francia** | 1,0018 | 44,4063 |
| **FR15502.1.1, Francia** | -0,6405 | 46,313 |
| **FR14421.1a.1, Francia** | 3,3255 | 48,4495 |
| **FR16222.3.1, Francia** | 6,0319 | 44,7123 |
| **FR15L27.1b.1, Francia** | 6,0876 | 48,0718 |
| **FR15517.1b.1, Francia** | 1,9426 | 48,6821 |
| **FR14816.2.1, Francia** | 2,7487 | 42,8265 |
| **Porquerolles island** | 6,2023 | 42,9888 |
| **Port-Cros island** | 6,3839 | 43,0068 |
| **Cap Lardier** | 6,6045 | 43,1809 |

Supplementary Table 4. Localities sampled in France by Emmanuelle Lapied, Daniel F. Marchán (orange) and Nuria Sánchez (pink).

| Locality | Longitude | Latitude |
|---|---|---|
| **Germany, Bavaria** | 11,8823 | 48,249 |
| **Germany** | No data | No data |
| **Wadi Kelt (Quilt), Israel** | No data | No data |
| **Stara Planina, Bulgaria** | No data | No data |
| **Crete 1** | 24,85584 | 35,30808 |
| **Crete 2** | 24,83732 | 35,31051 |
| **River near Velky Folkmar, Slovakia** | 21,01145 | 48,84588 |
| **Stream near Zvolen, Slovakia** | 19,1397 | 48,5592 |
| **River Muran, Jelsava, Slovakia** | 20,23907 | 48,62386 |
| **Stream in Selec, Slovakia** | 17,99599 | 48,77677 |
| **Stream in Krnca, Slovakia** | 18,26541 | 48,53594 |

| | | |
|---|---|---|
| Stream in Pdhradie, Slovakia | 18,63131 | 48,66155 |
| Vrica Stream in Predvrecko, Slovakia | 18,72669 | 48,96505 |
| Moravia, near Brno, Blanska, River Punkva banks, Czech Republic | 16,7403 | 49,4173 |
| Jasénka (Kotrlé), Czech Republic | 18,0233 | 49,3781 |
| **Locality** | **Longitude** | **Latitude** |
| Pardubický, Czech Republic | 16,8624 | 50,1499 |
| Mala Bystricka Stream, Czech Republic | 18,05408 | 49,3948 |
| Stream Knehyne in Prosteredni Becva, Czech Republic | 18,27853 | 49,46681 |
| Zabniczanka stream, Poland | 19,19397 | 49,55338 |
| Zylica stream in Szczyrk, Poland | 18,98406 | 49,69355 |
| UA14J02.1a.1, Russia | 29,9294 | 51,3885 |
| UA14J05.2.1,Russia | 30,12828 | 51,41435 |
| UA14J06.2b.1, Russia | 29,7038 | 51,48437 |
| UA14J04.2a.1, Russia | 30,1362 | 51,35207 |
| RU12728.1.1, Russia | 53,5102 | 63,5476 |
| Derbyshire, United Kingdom | -1,4900 | 52,766 |
| Wales, United Kingdom | -3,2278 | 51,4499 |
| Yukari Dueden Selalesi waterfall, N of Antalya | No data | No data |
| Scheldt River, near Wintam, Jan Soors, Belgium | 4,3047 | 51,1073 |
| Rossum, The Netherlands | 5,2980 | 51,793 |

Supplementary Table 5. Localities sampled in several countries by Christer Érseus and co-workers (blue), Emmanuelle Lapied (purple), Aleksandra Jablonska and Misel Jelic (green), Csaba Csuzdi and co-workers (black) and Marta Novo (grey).

| Species | Accesion number COI | Accesion number 16S | Accesion number 28S |
|---|---|---|---|
| ***Eiseniella tetraedra*** | OL457985-OL458610 | | OL471766-OL471904 |
| ***Carpetania matritensis*** | GQ409661.1 | JN209218.1 | GQ409652.1 |
| ***Lumbricus rubellus*** | KM611946.1 | KJ912567.1 | KJ912213.1 |

| | | | |
|---|---|---|---|
| ***Dendrobaena byblica*** | Domínguez et al. 2015; Pérez-Losada et al. 2015 | KJ912523.1 | Domínguez et al. 2015; Pérez-Losada et al. 2015 |
| ***Iberoscolex oliveirae*** | Domínguez et al. 2015; Pérez-Losada et al. 2015 | KJ912555.1 | Domínguez et al. 2015; Pérez-Losada et al. 2015 |
| ***Prosellodrilus biariculatus*** | Provided by the authors | KJ912586.1 | JN871950.1 |

Supplementary Table 6. Accesion numbers and references of all sequences used.

| | AUC | omission.rate | sensitivity | specificity | prop.correct | Kappa | |
|---|---|---|---|---|---|---|---|
| **Clade I** | 0,71 | 0,28 | 0,72 | 0,71 | 0,72 | 0,41 | |
| **Clade II** | 0,79 | 0,21 | 0,79 | 0,80 | 0,79 | 0,57 | |
| | **awc** | **bio13** | **bio6** | **bio7** | **clc** | **crust** | **parma** |
| **Clade I** | 15,09 | 14,65 | 15,47 | 16,97 | 15,06 | 11,95 | 10,80 |
| **Clade II** | 15,78 | 7,94 | 27,41 | 9,69 | 7,38 | 13,27 | 18,54 |

Supplementary Table 7. Above: comparison of the performance of the models for clades I and II. Below: relative contribution of the predictor variables to each model.

| UCM-LT number | Length (mm) | Dry weight (mg) | Number of segments | Clitellum | Tubercula pubertatis | Male pore | Seminal vesicles | Spermatechae | Spermiducal funnels | Observations |
|---|---|---|---|---|---|---|---|---|---|---|
| **30000** | 35 | 143 | 83 | 22-27 | 23-24-25 | 13 | 9-10-11-12 | not present | not present | |
| **30001** | 34 | 140 | 56 | 22-27 | 23-24-25 | 13 | 9-10-11-12 | not present | not present | |
| **30002** | 47 | 220 | 90 | 22-27 | 23-24-25 | 13 | 9-10-11-12 | not present | very small | |
| **30003** | 34.5 | 141 | 82 | 22-27 | 23-24-25 | 13 | 9-10-11-12 | not present | very small | |
| **30004** | 31 | 99 | 54 | 22-26 | 23-24-25 | 13 | 9-10-11-12 | not present | not present | |
| **30005** | 28 | 100 | 81 | 22-27 | 23-24-25 | 13 | 9-10-11-12 | not present | very small | |
| **30006** | 33 | 115 | 83 | 22-1n27 | 23-24-25-1n26 | 13 | 10-11--12 | not present | maybe on 10 without sperm | |
| **30007** | 16.5 | 40 | 46 | 22-26 | 23-24-25 | 13 | 9-10-11-12 | not present | very small | |
| **30008** | 15 | 56 | 52 | 23-26 | 23-24-25 | 13 | 8-10-11-12 | not present | maybe on 10 without sperm | |
| **30009** | 27 | 78 | 89 | | | 13 | | | | semi-mature |
| **30010** | 25 | 70 | 92 | | 23-24-25 | | | | | semi-mature |
| **30011** | 16 | 27 | 88 | | | | | | | inmature |
| **30012** | 18 | 15 | 80 | | | | | | | inmature |
| **30013** | 14 | 9 | 74 | | | | | | | inmature |
| **30014** | 11 | 10 | 87 | | | | | | | inmature |
| **30015** | 12.5 | 14 | 82 | | | | | | | inmature |
| **30016** | 17.5 | 80 | 89 | 22-26 | 23-24-25 | 13 | 10-11--12 | not present | not present | the earthworm was broken |
| **30017** | 15.5 | 44 | 92 | 22-26 | 23-24-25 | 13 | 9-10-11-12 | not present | very small | the earthworm was broken |
| **30018** | 20 | 45 | 79 | | | | | | | inmature |

| | | | | | | | | | | |
|---|---|---|---|---|---|---|---|---|---|---|
| **30019** | 21.5 | 49 | 85 | | | | | | | inmature |
| **30020** | 24 | 92 | 76 | 22-26 | 23-24-25 | 13 | 9-10-11-12 | very small | very small | |
| **30021** | 22 | 85 | 85 | 22-27 | 23-24-25-1n26 | 13 | 9-10-11-12 | not present | maybe on 10 without sperm | |
| **30022** | 21 | 63 | 72 | 1n22-1n27 | 23-1n26 | 13 | 9-10-11-12 | without sperm | not present | |
| **30023** | 24 | 78 | 88 | 22-27 | 23-24-25 | 13 | 9-10-11-12 | not present | not present | |
| UCM-LT number | **Length (mm)** | **Dry weight (mg)** | **Number of segments** | **Clitellum** | **Tubercula pubertatis** | **Male pore** | **Seminal vesicles** | **Spermatechae** | **Spermiducal funnels** | **Observations** |
| **30024** | 23.5 | 92 | 62 | 22-27 | 23-24-25 | 13 | 8-10-11-12 | very small | very small | |
| **30025** | 24 | 68 | 83 | 22-26 | 23-24-25-26 | 13 | 10-11--12 | not present | maybe on 10 without sperm | spermatophore without sperm between 11 and 12 |
| **30026** | 26.5 | 99 | 85 | 22-27 | 23-24-25-26 | 13 | 10-11--12 | not present | without sperm on left side, not present on right side | |
| **30027** | 19 | 53 | 84 | 23-26 | 23-24-25 | 13 | 10-11--12 | not present | not present | spermatophore without sperm on 11 |
| **30028** | 21 | 49 | 65 | | 23-24-25 | 13 | | | | semi-mature |
| **30029** | 21.5 | 50 | 84 | 22-27 | 23-24-25 | 13 | 9-10-11-12 | not present | very small | |
| **30030** | 19 | 44 | | | | | 11 -- 12 | not present | very small | the earthworm was broken |
| **30031** | 21 | 73 | 65 | 22-27 | 23-24-25-1n26 | 13 | left side: 11-12 right side:10-11--12 | not present | without sperm | |
| **30032** | 22 | 68 | 65 | 22-27 | 1n22-23-24-25 | 13 | left side: 11-12 right side: 10-11--12 | not present | without sperm | |
| **30033** | 19 | 42 | 69 | 22-26 | 23-24-25 | 13 | 9-10-11-12 | not present | without sperm | |

| | | | | | | | | | | |
|---|---|---|---|---|---|---|---|---|---|---|
| **30034** | | | | | | | | | | |
| **30035** | 15 | 26 | 65 | 22-27 | | | | | | inmature |
| **30036** | 20 | 56 | 71 | 20-25 | 21-22-23-24 | 11 | 8-9--10 | not present | without sperm | |
| **30037** | 21.5 | 54 | 82 | 18-24 | 20-21-22-1n23 | 9 | left side: 8-7--6 right side: 8-6 | not present | without sperm | |
| **30038** | 23.5 | 73 | 74 | 22-28 | 23-24-25-26 | 13 | 9-10-11-12 | not present | without sperm | |
| **30039** | 27 | 86 | 78 | 22-27 | 23-24-25-1n26 | 13 | 9-10-11-12 | not present | very small | |
| **30040** | 22 | 86 | 80 | 22-27 | 23-24-25 | 13 | 9-10-11-12 | not present | very small on left side | |
| **30041** | 26 | 85 | 86 | 22-27 | 23-24-25 | 13 | 9-10-11-12 | not present | very small | |
| **30042** | 19 | 53 | 88 | 22-27 | 23-24-25-1n26 | 13 | 11--12 | not present | without sperm | |
| **30043** | 24 | 81 | 80 | 22-27 | 23-24-25 | 13 | 9-10-11-12 | not present | not present | |
| **UCM-LT number** | **Length (mm)** | **Dry weight (mg)** | **Number of segments** | **Clitellum** | **Tubercula pubertatis** | **Male pore** | **Seminal vesicles** | **Spermatechae** | **Spermiducal funnels** | **Observations** |
| **30044** | 27 | 91 | 88 | 22-27 | 23-24-25 | 13 | 10--11--12 | not present | maybe on 10 without sperm | |
| **30045** | 23.5 | 80 | 85 | 23-27 | 23-24-25-1n26 | 13 | 10-11--12 | without sperm | without sperm | |
| **30046** | 27 | 86 | 87 | 22-27 | 23-24-25 | 13 | 9--12 | without sperm | without sperm | |
| **30047** | 23 | 62 | 80 | 22-27 | 23-24-25 | 13 | 9-10-11-12 | without sperm | without sperm | |
| **30048** | 26 | 90 | 85 | 22-27 | 23-24-25 | 13 | 9-10-11-12 | without sperm | without sperm | |
| **30049** | 24.5 | 71 | 91 | 22-27 | 23-24-25 | 13 | 10--11-12 | without sperm | without sperm | |
| **30050** | 25.5 | 85 | 83 | 22-27 | 23-24-25 | 13 | 10--11--12 | without sperm | without sperm | |
| **30051** | 24 | 80 | 84 | 22-27 | 23-24-25 | 13 | 10--11--12 | not present | without sperm | |

| | | | | | | | | | | |
|---|---|---|---|---|---|---|---|---|---|---|
| **30052** | 25 | 79 | 82 | 22-27 | 23-24-25-1n26 | 13 | 9-10-11-12 | without sperm | without sperm | |
| **30053** | 22.5 | 65 | 81 | 20-26 | 23-24-25 (right), 20 to 26 left | 13 | 9-10-11-12 | without sperm | not present | |
| **30054** | 25.5 | 85 | 88 | 22-27 | 23-24-25-1n26 | 13 | 9-10--12 | without sperm | very small | |
| **30055** | 25 | 70 | 84 | | | | | | | inmature |
| **30056** | 21 | 38 | 88 | | | | | | | inmature |
| **30057** | 22.5 | 43 | 84 | | | | | | | inmature |
| **30058** | 20.5 | 60 | 83 | | | | | | | inmature |
| **30059** | 22 | 43 | 86 | | | | | | | inmature |
| **30060** | 25 | 71 | 83 | | | | | | | inmature |
| **30061** | 22.5 | 59 | 79 | | | | | | | inmature |
| **30062** | 20.5 | 49 | 81 | | | | | | | inmature |
| **30063** | 19.5 | 57 | 80 | | | | | | | inmature |
| **30064** | 16 | 28 | 80 | | | | | | | inmature |
| **60065** | 17 | 25 | 76 | | | | | | | inmature |
| **30066** | 22 | 49 | 82 | | | | | | | inmature |
| **30067** | 20.5 | 45 | 89 | | | | | | | inmature |
| **UCM-LT number** | **Length (mm)** | **Dry weight (mg)** | **Number of segments** | **Clitellum** | **Tubercula pubertatis** | **Male pore** | **Seminal vesicles** | **Spermatechae** | **Spermiducal funnels** | **Observations** |
| **30068** | 14.5 | 24 | 68 | | | | | | | inmature |
| **30069** | 19.5 | 29 | 87 | | | | | | | inmature |
| **30070** | 16 | 30 | 77 | | | | | | | inmature |

| | | | | | | | | | | |
|---|---|---|---|---|---|---|---|---|---|---|
| **30071** | 17 | 27 | 88 | | | | | | | inmature |
| **30072** | 24 | 108 | 82 | 22-28 | 23-24-25 | 13 | 9--10-12 | not present | maybe on 10 without sperm | |
| **30073** | 23 | 67 | 83 | 22-27 | 23-24-25 | 13 | 9-10-11-12 | one on the left side without sperm | very small | |
| **30074** | 28.5 | 50 | 55 | 22-26 | 23-24-25 | 15 | 9--11-12 | one on the left side without sperm | very small | |
| **30075** | 17 | 21 | 83 | | | | | | | inmature |
| **30076** | 17 | 24 | 82 | | | | | | | inmature |
| **30077** | 20 | 33 | 87 | | | | | | | inmature |
| **30078** | | | | | | | | | | the earthworm was broken, inmature |
| **30079** | 12 | 15 | 52 | | | | | | | inmature |
| **30080** | 14.5 | 17 | 83 | | | | | | | inmature |
| **30081** | 13 | 12 | 62 | | | | | | | inmature |
| **30082** | 17.5 | 33 | 63 | | | | | | | inmature |
| **30083** | 25 | 63 | 85 | 22-26 | 23-24-25-1n26 | 13 | 11 -- 12 | not present | very small | |
| **30084** | 23 | 51 | 91 | 22-27 | 23-24-25-1n26 | 15 | 9 -- 12 | without sperm | very small | |
| **30085** | 18 | 42 | 87 | | | | | | | inmature |
| **30086** | 14 | 15 | 90 | | | | | | | inmature |
| **30087** | 11.5 | 11 | 89 | | | | | | | inmature |
| **30088** | | | | | | | | | | inmature |
| **30089** | 19 | 31 | 87 | | 23-24-25 | | | | | semi-mature |
| **30090** | 15 | 12 | 88 | | | | | | | inmature |

| UCM-LT number | Length (mm) | Dry weight (mg) | Number of segments | Clitellum | Tubercula pubertatis | Male pore | Seminal vesicles | Spermatechae | Spermiducal funnels | Observations |
|---|---|---|---|---|---|---|---|---|---|---|
| **30091** | | | | 22-27 | 23-24-25 | 13 | 9-10-11-12 | not present | not present | the earthworm was broken |
| **30092** | 29.5 | 106 | 86 | 22-27 | 23-24-25 | 13 | 9-10-11-12 | without sperm | without sperm | |
| **30093** | | | | 22-27 | 23-24-25 | 13 | 9--11--12 | not present | not present | the earthworm was broken |
| **30994** | 33 | 94 | 79 | 22-27 | 23-24-25 | 13 | 9-10-11-12 | without sperm | very small | |
| **30095** | 20 | 49 | 68 | 22-27 | 23-24-25-1n26 | 13 | 8-10-11-12 | not present | not present | |
| **30096** | 34.5 | 113 | 88 | 22-26 | 23-24-25-1n26 | 13 | 11 -- 12 | not present | not present | |
| **30097** | 27.5 | 66 | 83 | 22-27 | 23-24-25-26 | 13 | 11 -- 12 | without sperm | not present | |
| **30098** | 31.5 | 115 | 82 | 22-1n28 | 23-24-25 | 13 | 9--11--12 | not present | not present | |
| **30099** | 29.5 | 82 | 87 | 22-27 | 23-24-25-1n26 | 13 | 9-10-11-12 | not present | very small | |
| **30100** | 30 | 100 | 73 | 22-28 | 23-24-25-26 | 13 | 10--11--12 | without sperm | not present | |
| **30101** | 17 | 26 | 86 | | | | | | | |
| **30102** | 20 | 46 | 78 | | | | | | | |
| **30103** | 18 | 28 | 81 | | | | | | | |
| **30104** | 19 | 57 | 69 | 22-27 | 23-24-25 | 13 | 9--11--12 | without sperm | not present | |
| **30105** | 19.5 | 36 | 87 | | | 13 | | | | semi-mature |
| **30106** | | | | | | | | | | inmature the earthworm was broken |
| **30107** | 10 | 8 | 90 | | | | | | | inmature |
| **30108** | 19 | 33 | 103 | | | | | | | inmature |

| | | | | | | | | | | |
|---|---|---|---|---|---|---|---|---|---|---|
| **30109** | 16 | 19 | 69 | | | | | | | inmature |
| **30110** | 10 | 5 | 90 | | | | | | | inmature |
| **30111** | 35 | 176 | 85 | 22-27 | 23-24-25 | 15 | 9--11-12 | not present | not present | |
| **30112** | 35 | 232 | | 22-27 | 23-24-25 | 13 | 9--11--12 | without sperm | very small | the earthworm was broken |
| **30113** | 35 | 194 | 89 | 22-27 | 23-24-25-26 | 13 | 9-10-11-12 | not present | very small | |
| **30117** | 22 | 100 | 52 | 22-26 | 23-24-25 | 15 | 9--11--12 | without sperm | very small | |
| **30118** | 19.5 | 61 | 60 | 22-27 | 23-24-25 | 15 | 9--11--12 | not present | very small | |
| **UCM-LT number** | **Length (mm)** | **Dry weight (mg)** | **Number of segments** | **Clitellum** | **Tubercula pubertatis** | **Male pore** | **Seminal vesicles** | **Spermatechae** | **Spermiducal funnels** | **Observations** |
| **30119** | 33.5 | 143 | 77 | 22-27 | 23-24-25 | 15 | 9--11--12 | without sperm | very small | |
| **30120** | 17.5 | 30 | 84 | | | | | | | inmature |
| **30121** | 26 | 129 | 77 | 22-27 | 23-24-25-1n26 | 13 | 9--11--12 | not present | very small | |
| **30122** | 26.5 | 115 | 86 | 22-27 | 23-24-25-1n26 | 13 | 9--11--12 | not present | not present | |
| **30123** | 25 | 108 | 79 | 22-27 | 23-24-25 | 13 | 9--11--12 | not present | very small | |
| **30124** | 28 | 128 | 93 | 22-27 | 23-24-25 | 13 | 9--11--12 | not present | not present | |
| **30125** | 33 | 168 | 72 | 22-27 | 23-24-25 | 13 | 9--11--12 | without sperm | very small | |
| **30126** | 29 | 150 | 84 | 22-28 | 23-24-25 | 13 | 9--11--12 | without sperm | very small | |
| **30127** | 16.5 | 74 | 46 | 22-26 | 23-24-25 | 13 | 9--11--12 | not present | maybe on 10 without sperm | |
| **30128** | 32 | 150 | 82 | 22-27 | 23-24-25 | 13 | 9--12 | without sperm | maybe on 10 without sperm | |
| **30129** | 28.5 | 114 | 85 | 22-27 | 23-24-25 | 13 | 9-10-11-12 | without sperm | maybe on 10 without sperm | |

| | | | | | | | | | | |
|---|---|---|---|---|---|---|---|---|---|---|
| **30130** | 23 | 105 | 65 | 22-26 | 23-24-25-1n26 | 13 | 9--11--12 | not present | not present | |
| **30131** | 21 | 63 | 79 | | | 13 | | | | semi-mature |
| **30132** | 20 | 47 | 82 | | | 13 | | | | semi-mature |
| **30133** | 18 | 45 | 84 | | | | | | | inmature |
| **30134** | 13.5 | 20 | 77 | | | | | | | inmature |
| **30135** | 24.5 | 86 | 89 | 22-27 | 23-24-25-1n26 | 13 | 10--11--12 | not present | not present | |
| **30136** | 23 | 44 | 83 | 22-27 | 23-24-25-26 | 13 | 10--11--12 | not present | not present | |
| **30137** | 20 | 57 | 69 | | 23-24-25 | 13 | | | | semi-mature |
| **30138** | 21 | 98 | 59 | 22-27 | 23-24-25-26 | 13 | 9--11--12 | without sperm | very small | |
| **30139** | 32 | 85 | 81 | 22-27 | 23-24-25 | 13 | 9-10-11-12 | not present | not present | |
| **30140** | 16 | 26 | 84 | | | | | | | inmature |
| **30141** | 27 | 81 | 82 | | | | | | | inmature |
| **30142** | 15 | 16 | 84 | | | | | | | inmature |
| **UCM-LT number** | **Length (mm)** | **Dry weight (mg)** | **Number of segments** | **Clitellum** | **Tubercula pubertatis** | **Male pore** | **Seminal vesicles** | **Spermatechae** | **Spermiducal funnels** | **Observations** |
| **30143** | 22.5 | 82 | 59 | 22-27 | 23-24-25-1n26 | 13 | 9--11--12 | not present | maybe on 10 without sperm | |
| **30144** | 28 | 76 | 75 | 22-27 | 23-24-25-1n26 | 13 | 8-10-11-12 | not present | not present | |
| **30145** | 36 | 145 | 83 | 22-27 | 23-24-25 | 13 | 9-10-11-12 | not present | not present | |
| **30146** | 31.5 | 90 | 84 | 22-27 | 23-24-25-1n26 | 13 | 9--11--12 | not present | not present | |
| **30147** | 15 | 17 | 41 | 22-27 | No presentes | 13 | 9--12 | without sperm | very small | |

| | | | | | | | | | | |
|---|---|---|---|---|---|---|---|---|---|---|
| **30148** | 32 | 92 | 81 | 22-27 | 1n22-23-24-25-1n26 | 13 | 9-10-11-12 | without sperm | not present | |
| **30149** | 30 | 96 | 78 | 22-27 | 23-24-25-1n26 | 13 | 9-10-11-12 | without sperm | very small | |
| **30150** | 38 | 134 | 83 | 22-27 | 23-24-25 | 13 | 9-10-11-12 | not present | very small | |
| **30151** | 29.5 | 78 | 78 | 22-27 | 23-24-25 | 13 | 9-10-11-12 | one on the right side without sperm | not present | |
| **30152** | 30 | 83 | 65 | 22-27 | 23-24-25-26 | 13 | 9-10-11-12 | not present | not present | |
| **30153** | 28.5 | 105 | 63 | 22-26 | 23-24-25 | 13 | 9--11--12 | without sperm | very small | |
| **30154** | 20 | 56 | 51 | 22-26 | 23-24-25 | 13 | 9-10-11-12 | not present | very small | |
| **30155** | 32.5 | 121 | 85 | 22-27 | | 13 | 10--11--12 | not present | not present | |
| **30156** | 28 | 93 | 72 | 22-27 | 23-24-25 | 13 | 9-10-11-12 | not present | very small | |
| **30157** | 26 | 53 | 78 | 22-27 | 23-24-25-1n26 | 13 | 9-10-11-12 | without sperm | not present | |
| **30158** | 26.5 | 67 | 56 | 22-26 | 23-24-25-1n26 | 13 | 9-10-11-12 | without sperm | not present | |
| **30159** | 36.5 | 133 | 80 | 22-26 | 23-24-25 | 13 | 9-10-11-12 | not present | not present | |
| **30160** | 24.5 | 66 | 69 | 22-27 | 26-24-25-1n26 | 13 | 9-10-11-12 | not present | not present | |
| **30161** | 23 | 69 | 55 | 22-27 | 23-24-25 | 13 | 9-10-11-12 | not present | not present | |
| **30162** | 22 | 39 | 84 | | | | | | | semi-mature |
| **30163** | 27 | 56 | 84 | 22-27 | 23-24-25 | 13 | 10--11--12 | not present | not present | |
| **30164** | 28 | 64 | 78 | 22-27 | 23-24-25-26 | 13 | 9-10-11-12 | without sperm | not present | |
| **30165** | 31 | 100 | 84 | 22-27 | 23-24-25-1n26 | 13 | 9-10-11-12 | not present | not present | |

| UCM-LT number | Length (mm) | Dry weight (mg) | Number of segments | Clitellum | Tubercula pubertatis | Male pore | Seminal vesicles | Spermatechae | Spermiducal funnels | Observations |
|---|---|---|---|---|---|---|---|---|---|---|
| **30166** | 25 | 85 | 62 | 22-26 | 23-24-25-1n26 | 13 | 9-10-11-12 | not present | very small | |
| **30167** | 29.5 | 79 | 70 | 22-27 | 23-24-25-1n26 | 13 | 9-10-11-12 | without sperm | not present | |
| **30168** | 21.5 | 53 | 68 | 22-27 | 23-24-25 | 13 | 9--11--12 | without sperm | very small | |
| **30169** | 23 | 52 | 51 | 22-26 | 23-24-25-1n26 | 13 | 9-10-11-12 | not present | very small | |
| **30170** | | | | 22-27 | 23-24-25-1n26 | 13 | 11--12 | not present | very small | the earthworm was broken |
| **30171** | 19 | 25 | 83 | | | | | | | inmature |
| **30172** | 18.5 | 20 | 82 | | | | | | | inmature |
| **30173** | 15 | 19 | 64 | | | | | | | inmature |
| **30174** | 13 | 14 | 74 | | | | | | | inmature |
| **30175** | 22 | 64 | 87 | 22-27 | 23-24-25 | 13 | 10--11--12 | without sperm | not present | |
| **30176** | 15.5 | 56 | 33 | 22-27 | 23-24-25-26 | 13 | 9-10-11-12 | without sperm | very small | |
| **30177** | 41.5 | 159 | 81 | 22-28 | 23-24-25-26 | 13 | 9-10-11-12 | not present | not present | |
| **30178** | 22.5 | 61 | 82 | 22-28 | 24-25-26 | 13 | 9-10-11-12 | without sperm | not present | |
| **30179** | 29 | 114 | 81 | 22-27 | 24-25-26 | 13 | 9-10-11-12 | without sperm | not present | |
| **30180** | 28 | 90 | 82 | 22-28 | 24-25-26 | 13 | 9--11--12 | without sperm | maybe on 10 without sperm | |
| **30181** | 28.5 | 67 | 75 | 22-27 | 23-24-25-26 | 13 | 9-10-11-12 | not present | not present | |
| **30182** | 21.5 | 46 | 86 | 22-27 | 23-24-25 | 13 | 10--11--12 | without sperm | not present | |
| **30183** | | | | 22-27 | 23-24-25-1n26 | 13 | 9-10-11-12 | without sperm | maybe on 10 without sperm | the earthworm was broken |

| | | | | | | | | | | |
|---|---|---|---|---|---|---|---|---|---|---|
| **30184** | 11 | 7 | 80 | | | | | | | inmature |
| **30185** | | | | | | | | | | inmature |
| **30186** | 11.5 | 8 | 86 | | | | | | | inmature |
| **30187** | 12 | 9 | 79 | | | | | | | inmature |
| **30188** | 18.5 | 30 | 84 | | | | | | | inmature |
| **30189** | | | | | | | | | | inmature |
| **UCM-LT number** | **Length (mm)** | **Dry weight (mg)** | **Number of segments** | **Clitellum** | **Tubercula pubertatis** | **Male pore** | **Seminal vesicles** | **Spermatechae** | **Spermiducal funnels** | **Observations** |
| **30190** | 17 | 19 | 80 | | | | | | | inmature |
| **30191** | 12 | 9 | 73 | | | | | | | inmature |
| **30192** | 24 | 106 | 82 | | 23-24-25 | 13 | | | | semi-mature |
| **30193** | 20 | 64 | 80 | 22-27 | 23-24-25-26 | 13 | 9-11--12 | without sperm | not present | |
| **30194** | 16.5 | 54 | 61 | | | 13 | | | | semi-mature |
| **30195** | 17 | 31 | 86 | | | | | | | inmature |
| **30196** | 17.5 | 48 | 85 | | | | | | | inmature |
| **30197** | 19 | 61 | 71 | | | | | | | inmature |
| **30198** | 28.5 | 106 | 76 | 22-27 | 23-24-25-26 | 13 | 9-10-11-12 | without sperm | not present | |
| **30199** | 26.5 | 134 | 73 | 22-27 | 23-24-25 | 13 | 9-10-11-12 | without sperm | not present | |
| **30200** | 27 | 101 | 84 | 22-27 | 23-24-25 | 13 | 8-9-10-11-12 | without sperm | not present | |
| **30201** | 28 | 134 | 77 | 22-27 | 23-24-25 | 13 | 9--11--12 | without sperm | not present | |
| **30202** | 28.5 | 131 | 83 | 22-27 | 23-24-25-26 | 13 | 9-10-11-12 | without sperm | very small | |
| **30203** | 26.5 | 142 | 76 | 22-27 | 23-24-25 | 13 | 9-10-11-12 | without sperm | very small | |
| **30204** | 27 | 122 | 81 | 22-27 | 23-24-25-1n26 | 13 | 9--11--12 | without sperm | not present | |

| | | | | | | | | | | |
|---|---|---|---|---|---|---|---|---|---|---|
| **30205** | 26.5 | 115 | 81 | 22-27 | 23-24-25-1n26 | 13 | 9--11--12 | without sperm | very small | |
| **30206** | 24 | 91 | 84 | 22-27 | 23-24-25 | 13 | 9--11--12 | without sperm | not present | |
| **30207** | 21 | 74 | 65 | 22-27 | 23-24-25-1n26 | 13 | 10--11--12 | without sperm | not present | |
| **30208** | 25 | 106 | 80 | 22-27 | 23-24-25-1n26 | 13 | 10--11--12 | without sperm | not present | |
| **30209** | 23 | 117 | 59 | 22-27 | 23-24-25 | 13 | 9-10-11-12 | without sperm | not present | |
| **30210** | | | | 22-27 | 23-24-25 | 13 | 9--11--12 | without sperm | very small | the earthworm was broken |
| **30211** | | | | 22-26 | 23-24-25 | 13 | 8-10-11-12 | without sperm | not present | the earthworm was broken |
| **30212** | 25.5 | 116 | 82 | | 23-24-25 | 13 | | | | semi-mature |
| **30213** | 25 | 96 | 80 | | 23-24-25 | 13 | | | | semi-mature |
| **UCM-LT number** | **Length (mm)** | **Dry weight (mg)** | **Number of segments** | **Clitellum** | **Tubercula pubertatis** | **Male pore** | **Seminal vesicles** | **Spermatechae** | **Spermiducal funnels** | **Observations** |
| **30214** | 24 | 99 | 81 | | 23-24-25-1n26 | 13 | | | | semi-mature |
| **30215** | 21.5 | 90 | 80 | | 23-24-25-1n26 | 13 | | | | semi-mature |
| **30216** | 24 | 95 | 83 | | 23-24-25-1n26 | 13 | | | | semi-mature |
| **30217** | 21 | 86 | 66 | | 23-24-25-1n26 | 13 | | | | semi-mature |
| **30218** | 20.5 | 85 | 63 | | 23-24-25-1n26 | 13 | | | | semi-mature |
| **30219** | 17 | 59 | 60 | | 23-24-25-1n26 | 13 | | | | semi-mature |

| | | | | | | | | | | |
|---|---|---|---|---|---|---|---|---|---|---|
| **30220** | 20 | 62 | 77 | | 23-24-25-1n26 | 13 | | | | semi-mature |
| **30221** | 23.5 | 93 | 81 | | 23-24-25 | 13 | | | | semi-mature |
| **30222** | 17 | 66 | 75 | | 23-24-25-1n26 | 13 | | | | semi-mature |
| **30223** | | | | | 23-24-25 | 13 | | | | semi-mature |
| **30224** | 21.5 | 68 | | | | 13 | | | | semi-mature and the earthworm was broken |
| **30225** | 22 | 72 | 81 | | | 13 | | | | semi-mature |
| **30226** | 12.5 | 44 | 65 | | | | | | | inmature |
| **30227** | 19.5 | 74 | 69 | | | | | | | inmature |
| **30228** | 16.5 | 38 | 84 | | | | | | | inmature |
| **30229** | 21.5 | 60 | 96 | | | | | | | inmature |
| **30230** | 18.5 | 36 | 76 | | | | | | | inmature |
| **30231** | 17.5 | 41 | 83 | | | | | | | inmature |
| **30232** | 24.5 | 87 | 81 | 22-26 | 23-24-25 | 13 | 9--11--12 | not present | not present | |
| **30233** | 21 | 47 | 83 | | 24-25 | 13 | | | | semi-mature |
| **30234** | 21 | 41 | 84 | 22-26 | 23-24-25 | 13 | 11 -- 12 | not present | not present | |
| **30235** | 19.5 | 35 | 75 | 22-25 | 1n22-23-24-1n25 | 13 | 11 -- 12 | not present | not present | |
| **30236** | | | | 22-27 | 24-25-26 | 13 | 9-10-11-12 | not present | very small | broken |
| **UCM-LT number** | **Length (mm)** | **Dry weight (mg)** | **Number of segments** | **Clitellum** | **Tubercula pubertatis** | **Male pore** | **Seminal vesicles** | **Spermatechae** | **Spermiducal funnels** | **Observations** |
| **30237** | 22 | 82 | 64 | 22-27 | 23-24-25 | 13 | 10--11--12 | without sperm | very small | |

| | | | | | | | | | | |
|---|---|---|---|---|---|---|---|---|---|---|
| **30238** | 27 | 75 | 86 | 22-27 | 23-24-25-1n26 | 13 | 11 -- 12 | without sperm | not present | |
| **30239** | 21.5 | 84 | 58 | 22-27 | 23-24-25-1n26 | 13 | 9--11--12 | not present | not present | |
| **30240** | 20 | 69 | 63 | 22-27 | 23-24-25-1n26 | 13 | 9-10-11-12 | without sperm | not present | |
| **30241** | 22 | 104 | 65 | 22-28 | 23-24-25-26 | 13 | 9-10-11-12 | without sperm | very small. | |
| **30242** | 24 | 70 | 85 | 22-27 | 23-24-25 | 13 | 9-10-11-12 | without sperm | not present | |
| **30243** | 25 | 75 | 87 | 22-27 | 1n23-24-25-26 | 13 | 9--11--12 | without sperm | not present | |
| **30244** | 22 | 65 | 80 | 22-27 | 23-24-25 | 13 | 9-10-11-12 | without sperm | very small | |
| **30245** | 22.5 | 62 | | 22-27 | 23-24-25 | 13 | 9--11--12 | without sperm | very small | the earthworm was broken |
| **30246** | 22 | 67 | 88 | 22-28 | 24-25-26 | 13 | 9-10-11-12 | without sperm | not present | |
| **30247** | 29 | 100 | 90 | 22-28 | 23-24-25-1n26 | 13 | 9-10-11-12 | not present | not present | |
| **30248** | 19 | 71 | 66 | 22-28 | 23-24-25-1n26 | 13 | 9--11--12 | without sperm | not present | |
| **30249** | 29.5 | 79 | 89 | 22-28 | 23-24-25-1n26 | 13 | 9--11--12 | not present | not present | |
| **30250** | 28.5 | 102 | 70 | 22-28 | 23-24-25-1n26 | 13 | 9--11--12 | not present | maybe on 10 without sperm | |
| **30251** | 26 | 87 | 88 | 22-28 | 23-24-25 | 13 | 11 -- 12 | not present | not present | |
| **30252** | | | | 22-27 | 23-24-25-26 | 13 | 9-11--12 | not present | not present | the earthworm was broken |
| **30253** | 27 | 107 | 75 | 22-28 | 24-25-26 | 13 | 9--11--12 | not present | not present | |
| **30254** | 41.5 | 161 | 86 | 22-28 | 24-25-26 | 13 | 9--11--12 | not present | not present | |

| | | | | | | | | | | |
|---|---|---|---|---|---|---|---|---|---|---|
| **30255** | 50.7 | 250 | 94 | 22-27 | 23-24-25 | 13 | 9--11--12 | one on the right side without sperm | not present | |
| **30258** | 28 | 109 | 66 | 22-1n27 | 23-24-25-1n26 | 13 | 9-10-11-12 | not present | not present | |
| **30259** | 40 | 172 | 92 | 22-26 | 23-24-25 | 13 | 9-10-11-12 | one on the left side without sperm | not present | |
| **30260** | 33.5 | 117 | 88 | 22-27 | 23-24-25-26 | 13 | 9-10-11-12 | not present | not present | |
| **UCM-LT number** | **Length (mm)** | **Dry weight (mg)** | **Number of segments** | **Clitellum** | **Tubercula pubertatis** | **Male pore** | **Seminal vesicles** | **Spermatechae** | **Spermidučal funnels** | **Observations** |
| **30262** | 20.5 | 39.3 | 84 | | 23-24-25-26 | 13 | | | | semi-mature |
| **30263** | 16.5 | 22 | 80 | | | | | | | inmature |
| **30264** | 12 | 10 | 87 | | | | | | | inmature |
| **30265** | | | | 22-28 | 24-25-26 | 13 | 9-10-11-12 | not present | not present | the earthworm was broken |
| **30266** | 23 | 71 | 88 | 22-27 | 23-24-25 | 13 | 9-10-11-12 | one on the left side without sperm | maybe on 11 without sperm | |
| **30267** | 22 | 67 | 83 | 22-27 | 23-24-25 | 13 | 9--10--12 | not present | without sperm | |
| **30268** | 21.5 | 46 | 89 | | | 15 | | | | semi-mature |
| **30269** | 25 | 92 | 85 | 22-27 | 24-25-26 | 13 | 9--11--12 | one on the right side without sperm | not present | |
| **30270** | | | | 22-27 | 23-24-25-1n26 | 15 | 9--11--12 | without sperm | not present | the earthworm was broken |
| **30271** | 25 | 76 | 87 | 22-27 | 24-25-26 | 13 | 10--11--12 | not present | not present | |
| **30272** | 14 | 40 | 54 | 22-27 | 23-24-25 | 13 | 9--10--11--12 | not present | not present | |
| **30273** | 26 | 112 | 86 | 22-27 | 23-24-25-1n26 | 13 | 9-10-11-12 | not present | very small | |

| | | | | | | | | | | |
|---|---|---|---|---|---|---|---|---|---|---|
| **30274** | 16 | 34 | 62 | | 23-24-25-1n26 | 13 | | | | semi-mature |
| **30275** | 25.5 | 96 | 81 | 23-28 | 24-25-26 | 14 | 9-10-11-12 | without sperm | not present | |
| **30276** | 26.5 | 110 | 84 | 22-27 | 23-24-25-26 | 13 | 9--11--12 | without sperm | maybe on 10 without sperm | |
| **30277** | 20 | 59 | 84 | | | 13 | | | | semi-mature |
| **30278** | 18.5 | 36 | 78 | | | | | | | inmature |
| **30279** | 17 | 38 | 84 | | 24-25-26 | 13 | | | | semi-mature |
| **30280** | 34 | 114 | 89 | 22-28 | 23-24-25-26 | 13 | 12 -- 11 | not present | not present | |
| **30281** | 23.5 | 116 | 73 | 22-28 | 23-24-25-1n26 | 13 | 9--11--12 | not present | not present | |
| **30282** | 29.5 | 141 | 86 | 22-27 | 23-24-25-26 | 13 | 9--11--12 | not present | not present | |
| **30283** | 23 | 80 | 96 | 22-28 | 23-24-25-1n26 | 13 | 9-10-11-12 | not present | not present | |
| **30284** | 30 | 150 | 90 | 22-28 | 23-24-25-1n26 | 13 | 9-10-11-12 | without sperm | without sperm | |
| **UCM-LT number** | **Length (mm)** | **Dry weight (mg)** | **Number of segments** | **Clitellum** | **Tubercula pubertatis** | **Male pore** | **Seminal vesicles** | **Spermatechae** | **Spermidučal funnels** | **Observations** |
| **30285** | | | | 22-27- | 23-24-25-1n26 | 13 | 9-10-11-12 | without sperm | without sperm | the earthworm was broken |
| **30286** | 29 | 70 | 81 | 22-28 | 23-24-25-26 | 13 | 8-9-11-12 | without sperm | without sperm | |
| **30287** | 24.5 | 60 | 87 | 22-26 | 23-24-25 | 13 | 9-10-11-12 | not present | not present | |
| **30288** | 23 | 100 | 66 | 22-28 | 23-24-25-26 | 13 | 9-10-11-12 | not present | not present | |
| **30289** | 12.5 | 10 | | | | | | | | inmature and broken |
| **30290** | 22.5 | 76 | 89 | 22-27 | 23-24-25 | 13 | 9-10-11-12 | not present | not present | |

| **UCM-LT number** | **Length (mm)** | **Dry weight (mg)** | **Number of segments** | **Clitellum** | **Tubercula pubertatis** | **Male pore** | **Seminal vesicles** | **Spermatechae** | **Spermidual funnels** | **Observations** |
|---|---|---|---|---|---|---|---|---|---|---|
| **30291** | 23 | 104 | 89 | 22-27 | 23-24-25-1n26 | 13 | 9-10-11-12 | not present | not present | |
| **30292** | 31 | 122 | | 22-28 | 24-25-26 | 13 | 9-10-11-12 | not present | not present | |
| **30293** | 22.5 | 98 | 58 | 22-27 | 23-24-25 | 13 | 9-10-11-12 | without sperm | very small | |
| **30299** | 25 | 113.5 | 86 | 22-27 | 24-25-26 | 13 | 10--11--12 | not present | not present | |
| **30300** | 24 | 113.7 | 84 | 22-27 | 23-24-25-1n26 | 13 | 9-10-11-12 | not present | not present | |
| **30301** | 23 | 72.1 | 88 | 22-26 | 23-24-25 | 13 | 11---12 | not present | not present | |
| **30302** | 24.5 | 108.2 | 87 | 22-27 | 23-24-25-1n26 | 13 | 8-10-11-12 | not present | not present | |
| **30303** | 23 | 97.7 | 92 | 22-27 | 23-24-25-1n26 | 13 | 11---12 | not present | without sperm | |
| **30304** | 20 | 78.6 | 84 | 22-26 | 23-24-25 | 13 | 9-10-11-12 | not present | not present | |
| **30305** | | | | | | 13 | 9---11-12 | not present | not present | the earthworm was broken |
| **30306** | 16 | 43.3 | 74 | | | 13 | | | | semi-mature |
| **30307** | 37 | 93.5 | 81 | 21-26 | 23-24-25 | 12 | 9--11-12 | not present | not present | |
| **30308** | 22 | 45.5 | 73 | 20-24 | 22-23-24 | 11 | 9-11--12 | without sperm | without sperm | |
| **30309** | 27 | 24.4 | 45 | | | 13 | | | | semi-mature |
| **30310** | 14 | 29.2 | 71 | | | 13 | | | | semi-mature |
| **30311** | 16 | 6.9 | 38 | | | | | | | inmature |
| **30312** | 14 | 12.4 | 81 | | | | | | | inmature |
| **30313** | 17.5 | 54.6 | 72 | 22-26 | 23-24-25 | 13 | 9--11--12 | not present | not present | |

| | | | | | | | | | | |
|---|---|---|---|---|---|---|---|---|---|---|
| **30314** | 12.5 | 16.2 | 74 | | | | | | | inmature |
| **30315** | 14 | 19.2 | 91 | | | | | | | inmature |
| **30316** | 9 | 7.1 | 85 | | | | | | | inmature |
| **30317** | 20.5 | 58.5 | 79 | 22-27 | 23-24-25 | 13 | 9---11-12 | not present | not present | |
| **30318** | 19 | 41.1 | 85 | 22-27 | 23-24-25-1n26 | 13 | 9--11--12 | not present | not present | |
| **30319** | 16.5 | 35.7 | 61 | 22-27 | 23-24-25 | 13 | 11---12 | not present | not present | |
| **30320** | 16 | 50.5 | 56 | 22-27 | 23-24-25-26-27 | 13 | 9--11--12 | not present | not present | |
| **30321** | | | | | | | | | | |
| **30322** | | | | | | | | | | |
| **30323** | | | | | | | | | | |
| **30324** | | | | | | | | | | |
| **30325** | 23.5 | 97.5 | 73 | 22-27 | 23-24-25 | 13 | 9--10 | not present | not present | |
| **30326** | 30 | 172 | 76 | 22-27 | 26-24-25-26 | 13 | 9-10-11-12 | not present | not present | |
| **30327** | 31 | 83 | 91 | 22-28 | 23-25-24-25-1n27 | 13 | 9-10-11-12 | not present | not present | |
| **30328** | 24 | 117.2 | | 22-27 | 23-24-25 | 13 | 11--12 | not present | not present | |
| **30329** | 19 | 39.7 | 93 | | 23-24-25-1n26 | 13 (and 14?) | | | | semi-mature |
| **30330** | 23.5 | 66.6 | 83 | 22-27 | 23-24-25-26 | 13 | 9--11-12 | without sperm | not present | |
| **30331** | 19 | 58.7 | 76 | | | 13 | | | | semi-mature |
| **30332** | 28 | 109.1 | 78 | 22-27 | 23-24-25 | 13 | 11--12 | not present | not present | |
| **30333** | 22 | 101.8 | 60 | 22-26 | 23-24-25 | 12 | 9--11--12 | not present | not present | |
| **30334** | 27 | 138.1 | 82 | 22-26 | 23-24-25 | 13 | 11--12 | without sperm | not present | |

| | | | | | | | | | | |
|---|---|---|---|---|---|---|---|---|---|---|
| **30335** | | | | 22-28 | 23-24-25 | 13 | 9--11--12 | not present | not present | the earthworm was broken |
| **30336** | 26 | 109.7 | 80 | 22-27 | 24-25-26 | 13 | 9-10-11-12 | not present | not present | |
| **30337** | | | | 22-28 | 23-24-25 | 13 | 9-10-11-12 | not present | not present | the earthworm was broken |
| **UCM-LT number** | **Length (mm)** | **Dry weight (mg)** | **Number of segments** | **Clitellum** | **Tubercula pubertatis** | **Male pore** | **Seminal vesicles** | **Spermatechae** | **Spermidical funnels** | **Observations** |
| **30338** | | | | 22-27 | 23-24-25-26 | 13 | 8-10-11-12 | not present | not present | the earthworm was broken |
| **30339** | 24 | 101.7 | 92 | 22-27 | 24-25-26 | 13 | 9-10-11-12 | not present | not present | |
| **30340** | | | | | | | | | | the earthworm was broken, semi-mature |
| **30341** | 11 | 14.8 | 85 | | | | | | | inmature |
| **30342** | | | | 22-27 | 23-24-25-26 | 13 | 10--11--12 | not present | not present | the earthworm was broken |
| **30343** | 19 | 61.4 | 81 | | 22-23-24 | 13 | | | | semi-mature |
| **30344** | | | | | | 13 | 9-10-11-12 | without sperm | without sperm | the earthworm was broken |
| **30345** | 14.5 | 58.2 | 40 | 22-27 | 1n23-24-25-1n26 | 13 | 9-10-11-12 | without sperm | without sperm | the earthworm was broken |
| **30346** | | | | | | 13 | | | | the earthworm was broken |
| **30348** | 24 | 59.5 | 78 | 22-27 | 23-24-25-1n26 | 13 | 9-10-11-12 | not present | not present | |
| **30349** | 24.5 | 72.5 | 80 | 22-27 | 23-24-25-1n26 | 13 | 11 -- 12 | not present | not present | |
| **30350** | 21 | 47.6 | 101 | | 23-24-25 | 13 | | | | semi-mature |

| | | | | | | | | | | |
|---|---|---|---|---|---|---|---|---|---|---|
| **30351** | 32 | 106 | 78 | 22-27 | 23-24-25-1n26 | 13 | 9--11--12 | not present | not present | |
| **30352** | 20 | 73.8 | 95 | 22-27 | 23-24-25-26 | 13 | 9-10-11-12 | not present | not present | |
| **30353** | 20 | 54.3 | 97 | 22-27 | 23-24-25-26 | 13 | 9-10-11-12 | not present | not present | |
| **30354** | 18 | 31.2 | 89 | | | | | | | inmature |
| **30355** | 26.5 | 94.3 | 97 | 22-27 | 23-24-25 | 13 | 9-10-11-12 | not present | not present | |
| **30356** | 22 | 55.3 | 85 | 22-27 | 23-24-25-26 | 13 | 9-10-11-12 | not present | not present | |
| **30357** | 20 | 58.4 | 82 | | | | | | | inmature |
| **30358** | 21.5 | 34.2 | | | | | | | | inmature |
| **30360** | 24 | 102.5 | 67 | 22-28 | 24-25-26 | 13 | 9-10-11-12 | not present | not present | |
| **30361** | 19.5 | 55.5 | 78 | 21-28 | 22-23-24-25-26 | 13 | 9-10-11-12 | not present | not present | |
| **30362** | 21.5 | 61.5 | 96 | 22-27 | 23-24-25-26 | 13 | 11 -- 12 | not present | not present | |
| **UCM-LT number** | **Length (mm)** | **Dry weight (mg)** | **Number of segments** | **Clitellum** | **Tubercula pubertatis** | **Male pore** | **Seminal vesicles** | **Spermatechae** | **Spermiducal funnels** | **Observations** |
| **30363** | 23.5 | 73.4 | 77 | 21-27 | 23-24-25 | 13 | 9-10-11-12 | not present | not present | |
| **30364** | 24 | 92.3 | 97 | 22-27 | 23-24-25-26 | 13 | 9--11--12 | one on the right side without sperm | not present | |
| **30365** | 28 | 94 | 87 | 22-27 | 23-24-25-1n26 | 13 | 9--11--12 | not present | not present | |
| **30366** | 24 | 109.7 | 62 | 22-28 | 24-25-26 | 13 | 12 -- 11 | not present | not present | |
| **30367** | 28 | 103 | 97 | 22-27 | 23-24-25 | 13 | 12 -- 11 | not present | not present | |
| **30368** | 21 | 68.2 | 60 | 22-26 | 23-24-25 | 13 | 13--11--12 | not present | not present | |
| **30369** | 19.5 | 87.8 | 58 | 22-28 | 23-24-25-26 | 13 | 9-10-11-12 | one on the right side without sperm | not present | |

| | | | | | | | | | | |
|---|---|---|---|---|---|---|---|---|---|---|
| **30370** | 16.5 | 24.2 | 92 | | | | | | | inmature |
| **30371** | 19 | 54.9 | 76 | 22-27 | 23-24-25 | 13 | 9 --12 | without sperm | not present | |
| **30372** | 16 | 36.7 | 71 | | | 9 | | | | semi-mature |
| **30373** | 18 | 27.1 | 74 | | | | | | | inmature |
| **30374** | 22.5 | 71.9 | 98 | 22-28 | 23-24-25-26 | 13 | 9-10-11-12 | without sperm | without sperm | |
| **30375** | 13.5 | 13.9 | 73 | | | | | | | inmature |
| **30376** | 12 | 11.8 | 77 | | | | | | | inmature |
| **30377** | 29.5 | 99 | 59 | 22-26 | 23-24-25 | 13 | 9--11--12 | not present | not present | |
| **30378** | 20.5 | 43.1 | 92 | 22-27 | 24-25-26 | 13 | 10--11-12 | not present | without sperm | |
| **30379** | 32.5 | 118.8 | 84 | 22-27 | 23-24-25 | 13 | 9-10-11-12 | without sperm | without sperm | |
| **30380** | 22 | 102.2 | 58 | 22-27 | 23-24-25 | 13 | 12 | without sperm | not present | |
| **30381** | 25.5 | 98.7 | 93 | 22-27 | 23-24-25 | 13 | 12 -- 11 | without sperm | without sperm | |
| **30382** | 32 | 128.4 | 82 | 23-28 | 24-25-26 | 13 | 12 -- 11 | not present | not present | |
| **30383** | 23 | 97.8 | 68 | 22-28 | 23-24-25 | 13 | 9-10-11-12 | one on each side without sperm | without sperm | |
| **30384** | 35.5 | 158.8 | 86 | 21-26 | 1n22-23-24-1n25 | 12 | 12 | not present | not present | |
| **30385** | | | | 22-27 | 23-24-25 | 13 | 9-10-11-12 | without sperm | without sperm | the earthworm was broken |
| **30386** | 29 | 123.7 | 80 | 22-27 | 23-24-25 | 13 | 9--11--12 | not present | not present | |
| **UCM-LT number** | **Length (mm)** | **Dry weight (mg)** | **Number of segments** | **Clitellum** | **Tubercula pubertatis** | **Male pore** | **Seminal vesicles** | **Spermatechae** | **Spermiducal funnels** | **Observations** |
| **30387** | 24.5 | 116.1 | 83 | 22-28 | 24-25-26 | 13 | 9-10-11-12 | one on each side without sperm | without sperm | |

| | | | | | | | | | | |
|---|---|---|---|---|---|---|---|---|---|---|
| **30388** | 32.5 | 148.8 | 86 | 22-26 | 23-24-25 | 13 | 9--11--12 | one on each side without sperm | not present | |
| **30389** | 22.5 | 83.7 | 79 | 22-27 | 23-24-25-26 | 13 | 9--11--12 | without sperm | without sperm | |
| **30390** | 21.5 | 82.5 | 60 | 22-27 | 23-24-25-26 | 13 | 9 -- 12 | without sperm | without sperm | |
| **30391** | 10 | 8.8 | 83 | | | | | | | inmature |
| **30392** | 6 | 4.5 | 67 | | | | | | | inmature |
| **30399** | 34.5 | 227.2 | 71 | 22-27 | 23-24-25 | 13 | 9--11--12 | not present | not present | |
| **30401** | 40 | 289.7 | 93 | 22-28 | 23-24-25-1n26 | 13 | 9-10-11-12 | one on each side without sperm | not present | |
| **30402** | 39 | 260.6 | 86 | 22-28 | 24-25-26 | 13 | 10--11--12 | one on right side without sperm | without sperm | |
| **30403** | 30 | 150.7 | 82 | 22-27 | 1n23-24-25-1n26 | 13 | 9--11--12 | not present | not present | |
| **30404** | | | | 22-27 | 23-24-25 | 13 | 10--11-12 | not present | not present | the earhtworm was broken |
| **30405** | 30.5 | 158.4 | 83 | 22-27 | 24-25-26 | 13 | 9-10-11-12 | one on right side without sperm | without sperm | |
| **30406** | 26.5 | 101.2 | 84 | | | 13 | | | | semi-mature |
| **30407** | 23 | 81.1 | 91 | 22-26 | 23-24-25 | 13 | 9-10-11-12 | one on left side without sperm | without sperm | |
| **30408** | 38.5 | 204.3 | 82 | 22-28 | 24-25-26 | 13 | 12 | not present | without sperm | |
| **30409** | 32 | 123.1 | | 21-25 | 22-23-24 | 12 | 9-11--12 | not present | without sperm | the earthworm was broken |
| **30410** | 22 | 99 | | 22-28 | 23-24-25 | 13 | 9-10-11-12 | not present | without sperm | the earthworm was broken |
| **30411** | | | | 22-27 | 24-25-26 | 13 | 9-10-11-12 | not present | not present | the earthworm was broken |
| **30412** | 22 | 61 | 91 | | | | | | | inmature |

| UCM-LT number | Length (mm) | Dry weight (mg) | Number of segments | Clitellum | Tubercula pubertatis | Male pore | Seminal vesicles | Spermatechae | Spermidučal funnels | Observations |
|---|---|---|---|---|---|---|---|---|---|---|
| **30413** | 30 | 93.2 | 74 | 22-28 | 23-24-25 | 12 | 9-10-11-12 | without sperm | without sperm | |
| **30414** | 14 | 30.8 | 69 | | | | | | | inmature |
| **30425** | 26.5 | 119.3 | 90 | 22-28 | 23-24-25-26-27 | 13 | 9-10-11-12 | without sperm | without sperm | |
| **30426** | 19 | 87.1 | 79 | | 23-24-25 | 13 | | | | semi-mature |
| **30427** | 18.5 | 85.6 | 75 | 22-28 | 24-25-26-27 | 13 | 9--11--12 | without sperm | not present | |
| **30428** | | | | 22-27 | 24-25-26 | 13 | 9--11--12 | without sperm | not present | |
| **30429** | 13 | 74.1 | 81 | | | 13 | | | | semi-mature |
| **30434** | 25 | 77.6 | 88 | 22-27 | 23-24-25 | 13 | 9-10-11-12 | one on right side without sperm | not present | |
| **30435** | 19 | 49 | 61 | 22-27 | 23-24-25 | 13 | 9-10-11-12 | without sperm | not present | |
| **30436** | 21 | 46.2 | 70 | | 24-25-26 | 13 | | | | semi-mature |
| **30437** | 27 | 79.4 | 76 | 22-27 | 23-24-25 | 13 | 9-10-11-12 | not present | not present | |
| **30438** | 17 | 69.7 | 59 | 19-23 | 20-21-22 | 9 | 9-10-11-12 | one on left side without sperm | not present | |
| **30439** | 19 | 58.3 | 80 | 22-28 | 23-24-25 | 13 | 9-10-11-12 | without sperm | without sperm | |
| **30440** | 26 | 87.6 | 70 | 22-27 | 23-24-25 | 13 | 9--11--12 | not present | not present | |
| **30441** | 22 | 102.8 | 88 | 22-27 | 23-24-25 | 13 | 9--11--12 | without sperm | without sperm | |
| **30442** | 25 | 93.2 | 91 | 21-25 | 21-22-23-24 | 13 | 9-10-11-12 | one on right side without sperm | not present | |
| **30443** | 24.5 | 87.5 | 84 | 23-28 | 24-25-26 | 13 | 10--11-12 | not present | not present | |
| **30444** | 25 | 90.2 | 89 | 22-27 | 23-24-25 | 13 | 9--11--12 | without sperm | not present | |
| **30445** | 26 | 97.2 | 91 | 22-27 | 23-24-25 | 13 | 9-10-11-12 | without sperm | without sperm | |

| | | | | | | | | | | |
|---|---|---|---|---|---|---|---|---|---|---|
| **30446** | 27 | 94.1 | 78 | 22-28 | 23-24-25 | 13 | 10--11--12 | not present | not present | |
| **30447** | 23.5 | 58.8 | 74 | | | 13 | | | | semi-mature |
| **30448** | 24 | 91.6 | 90 | 22-27 | 23-24-25 | 13 | 9-10-11-12 | | | |
| **30449** | 22.5 | 73.8 | 79 | 22-27 | 23-24-25 | 13 | 9-10-11-12 | not present | not present | |
| **30450** | 26 | 88.6 | 66 | 22-27 | 23-24-25 | 13 | 9-10-11-12 | one on right side without sperm | not present | |
| **30451** | 22.5 | 64.4 | 77 | 22-27 | 23-24-25 | 13 | 9-10-11-12 | without sperm | without sperm | |
| **30452** | 25.5 | 101.8 | 72 | 22-28 | 23-24-25 | 12 | 9-10-11-12 | one on right side without sperm | not present | |
| **30453** | 22.5 | 51.3 | 103 | 22-28 | 23-24-25 | 13 | 9-10-11-12 | without sperm | without sperm | |
| **UCM-LT number** | **Length (mm)** | **Dry weight (mg)** | **Number of segments** | **Clitellum** | **Tubercula pubertatis** | **Male pore** | **Seminal vesicles** | **Spermatechae** | **Spermiducal funnels** | **Observations** |
| **30454** | 24.5 | 88.7 | 76 | 22-27 | 23-24-25 | 13 | 9-10-11-12 | without sperm | without sperm | |
| **30455** | 25 | 91.3 | 55 | 22-27 | 23-24-25 | 13 | 9--11--12 | without sperm | not present | |
| **30456** | 22 | 79.9 | 81 | 22-27 | 23-24-25 | 13 | 9-10-11-12 | without sperm | not present | |
| **30457** | 21 | 68.7 | 65 | 22-27 | 23-24-25 | 13 | 9-10-11-12 | not present | not present | |
| **30458** | 23 | 91.3 | 83 | 22-27 | 23-24-25 | 13 | 9-10-11-12 | without sperm | without sperm | |
| **30459** | 26 | 86.7 | 77 | 22-28 | 24-25-26 | 13 | 9-10-11-12 | without sperm | without sperm | |
| **30460** | 23.5 | 91.4 | 80 | 22-27 | 23-24-25 | 13 | 9-10-11-12 | without sperm | not present | |
| **30461** | 24 | 69.9 | 89 | 22-27 | 23-24-25 | 13 | 9-10-11-12 | not present | not present | |
| **30462** | 23.5 | 73 | 73 | 22-27 | 23-24-25 | 13 | 9-10-11-12 | without sperm | without sperm | |
| **30463** | | | | 22-27 | 23-24-25 | 13 | 9-10-11-12 | without sperm | without sperm | the earthworm was broken |
| **30464** | 21.5 | 65.3 | 82 | 23-28 | 24-25-26 | 13 | 9--11--12 | without sperm | not present | |
| **30487** | 24 | 70.2 | 68 | 22-27 | 23-24-25 | 13 | 11 -- 12 | not present | not present | |

| | | | | | | | | | | |
|---|---|---|---|---|---|---|---|---|---|---|
| **30488** | 26 | 61.9 | 62 | 22-27 | 23-24-25 | 13 | 11 -- 12 | not present | not present | |
| **30489** | 21.5 | 54.8 | 52 | 22-26 | 23-24-25 | 13 | 11 -- 12 | not present | not present | |
| **30490** | | | | 22-28 | 23-24-25 | 13 | 10--11--12 | without sperm | not present | the earthworm was broken |
| **30491** | | | | 22-28 | 24-25-26 | 13 | 10--11--12 | no presents | not present | the earthworm was broken |
| **30492** | 26 | 73.4 | 79 | 22-28 | 24-25-26 | 13 | 11 -- 12 | not present | not present | |
| **30493** | 23.5 | 55.1 | 85 | 22-28 | 24-25-26 | 13 | 11 -- 12 | not present | not present | |
| **30494** | 16 | 43.4 | 56 | 22-26 | 23-24-25 | 13 | 9-10-11-12 | without sperm | without sperm | |
| **30496** | 17.5 | 49.7 | 70 | 22-27 | 23-24-25 | 13 | 9--11-12 | not present | not present | |
| **30497** | | | | | | 13 | 11 -- 12 | not present | not present | the earthworm was broken |
| **30498** | 29 | 93.5 | 82 | 22-28 | 23-24-25 | 13 | 10--11--12 | not present | not present | |
| **30499** | | | | 22-28 | 23-24-25 | 13 | 11--12 | not present | not present | the earthworm was broken |
| **30500** | 25 | 79.1 | 80 | 22-27 | 23-24-25 | 13 | 10--11--12 | not present | not present | |
| **UCM-LT number** | **Length (mm)** | **Dry weight (mg)** | **Number of segments** | **Clitellum** | **Tubercula pubertatis** | **Male pore** | **Seminal vesicles** | **Spermatechae** | **Spermiducal funnels** | **Observations** |
| **30501** | 22 | 81.3 | 85 | | 23-24-25 | 13 | | | | semi-mature |
| **30502** | 16 | 41.4 | 77 | | | | | | | inmature |
| **30503** | | | | | | | | | | inmature, the earthworm was broken |
| **30504** | 18 | 54.1 | 68 | | | | | | | inmature |
| **30505** | 17 | 41.4 | 72 | 22-27 | 23-24-25 | 13 | 10--11--12 | not present | not present | |
| **30506** | 18 | 59.5 | 71 | 22-27 | 24-25-26 | 13 | 11 -- 12 | not present | not present | |

| | | | | | | | | | | |
|---|---|---|---|---|---|---|---|---|---|---|
| **30507** | 21 | 62.4 | 80 | 22-27 | 23-24-25 | 13 | 11 -- 12 | not present | not present | |
| **30508** | 12 | 20.1 | 69 | | | | | | | inmature |
| **30509** | 14 | 19.1 | 75 | | | | | | | inmature |
| **30510** | 11 | 6.5 | 86 | | | | | | | inmature |
| **30511** | 12.5 | 12.7 | 68 | | | | | | | inmature |
| **30512** | 14 | 19.5 | 70 | | | | | | | inmature |
| **30513** | | | | | | | | | | posterior part of the body |
| **30514** | 32 | 128.1 | 99 | 22-27 | 23-24-25 | 13 | 9--11--12 | not present | not present | |
| **30515** | 35 | 132.4 | 82 | 22-27 | 23-24-25 | 13 | 9--11--12 | not present | not present | |
| **30516** | 30 | 84.5 | 96 | 22-28 | 24-25-26 | 13 | 9,11,12 | not present | not present | |
| **30517** | 36.5 | 144.7 | 100 | 22-27 | 23-24-25-1n26 | 13 | 9-10-11-12 | not present | not present | |
| **30518** | 18.5 | 36 | 77 | | | | | | | inmature |
| **30519** | 12 | 52.1 | 56 | 22-26 | 23-24-25 | 13 | 9--11--12 | without sperm | without sperm | |
| **30520** | 26 | 129.7 | 81 | 22-27 | 23-24-25-26 | 13 | 9--11--12 | without sperm | not present | |
| **30521** | 14 | 28.4 | 82 | | | | | | | inmature |
| **30522** | | | | 22-27 | 23-24-25 | 13 | 9--11--12 | not present | not present | the earthworm was broken |
| **30523** | 16 | 26.5 | 92 | | 23-24-25 | 13 | | | | semi-mature |
| **30524** | | | | 22-27 | 23-24-25 | 12 | 9--11--12 | | | the earthworm was broken |
| **UCM-LT number** | **Length (mm)** | **Dry weight (mg)** | **Number of segments** | **Clitellum** | **Tubercula pubertatis** | **Male pore** | **Seminal vesicles** | **Spermatechae** | **Spermiducal funnels** | **Observations** |
| **30525** | 16 | 40.8 | 95 | | 23-24-25 | 13 | | | | semi-mature |

| | | | | | | | | | | |
|---|---|---|---|---|---|---|---|---|---|---|
| **30526** | | | | | | 13 | | | | the earthworm was broken semi-mature |
| **30527** | 10 | 13.2 | 74 | | | | | | | inmature |
| **30528** | 10.5 | 11.8 | 80 | | | | | | | inmature |
| **30529** | | | | | | | | | | the earthworm was broken e inmature |
| **30530** | 16 | 20.6 | 77 | | | | | | | inmature |
| **30531** | 19.5 | 40.4 | 70 | | | 13 | | | | semi-mature |
| **30532** | 21 | 69.4 | 75 | 22-27 | 23-24-25 | 14 | 9-10-11-12 | without sperm | not present | |
| **30533** | | | | | | | | | | inmature the earthworm was broken |
| **30534** | 12 | 16.3 | 72 | | | | | | | inmature |
| **30535** | 15 | 22.8 | 96 | | | | | | | inmature |
| **30536** | 10 | 12.4 | 82 | | | | | | | inmature |
| **30537** | 18 | 38.3 | 88 | | | | | | | inmature |
| **CE2518** | | | | 22-27 | 23-24-25 | 13 | 9-10-11-12 | without sperm | without sperm | |
| **ce2771** | | | | 22-29 | 24-25-26 | 13 | 9-10-11-12 | without sperm | not present | |
| **ce2772** | | | | 22-28 | 23-24-25 | 13 | 9-10-11-12 | not present | not present | |
| **ce2779** | | | | | | | | | | |
| **ce2780** | | | | | | 13 | | | | |
| **ce2864** | | | | 22-26 | 23-24-25 | 13 | 9-10-11-12 | not present | not present | |
| **ce2865** | | | | | | 13 | | | | |
| **ce2876** | | | | 22-27 | 24-25-26 | 13 | 9-10-11-12 | not present | not present | |
| **ce2877** | | | | 22-27 | 23-24-25 | 13 | 9-10-11-12 | without sperm | without sperm | |
| **ce2878** | | | | 22-27 | 23-24-25 | 13 | 9-10-11-12 | not present | not present | |

| UCM-LT number | Length (mm) | Dry weight (mg) | Number of segments | Clitellum | Tubercula pubertatis | Male pore | Seminal vesicles | Spermatechae | Spermiducal funnels | Observations |
|---|---|---|---|---|---|---|---|---|---|---|
| **ce2879** | | | | 22-27 | 23-24-25 | 11 | 9-10-11-12 | one on left side without sperm | not present | |
| **ce3813** | | | | 23-28 | 25-26-27 | 13 , 14 | 9-10-11-12 | without sperm | without sperm | |
| **ce3814** | | | | 22-28 | 24-25-26 | 13 | 9-10-11-12 | without sperm | not present | |
| **ce4070** | | | | 22-27 | 23-24-25 | 13 | 9-10-11-12 | not present | not present | |
| **ce4207** | | | | 22-27 | 23-24-25 | 13 | 9-10-11-12 | not present | not present | |
| **ce4208** | | | | 22-27 | 23-24-25 | 13 | 11 -- 12 | not present | not present | |
| **ce4209** | | | | 22-28 | 23-24-25-26-27-28 | 13 | 11 --12 | not present | not present | |
| **ce4211** | | | | 22-27 | 23-24-25 | 13 | 11 -- 12 | not present | not present | |
| **ce4213** | | | | | 23-24-25 | 13 | | | | |
| **ce4361** | | | | 22-27 | 23-24-25 | 13 | 9--11--12 | not present | not present | |
| **ce4362** | | | | 22-27 | 23-24-25 | 13 | 11 -- 12 | not present | not present | |
| **ce4395** | | | | 22-27 | 23-24-25 | 13 | 11 -- 12 | without sperm | very small | |
| **ce4396** | | | | 22-27 | 23-24-25 | 13 | 11 -- 12 | not present | not present | |
| **ce4398** | | | | 22-27 | 23-24-25 | 13 | 11 -- 12 | without sperm | without sperm | |
| **ce4399** | | | | 22-27 | 13-24-25 | 13 | 11 -- 12 | not present | not present | |
| **ce4400** | | | | 22-27 | 23-24-25 | 13 | 11 -- 12 | not present | not present | |
| **ce4401** | | | | 22-27 | 23-24-25 | 13 | 11 -- 12 | without sperm | without sperm | |
| **ce4448** | | | | 22-27 | 23-24-25 | 13 | 11 -- 12 | without sperm | without sperm | |
| **ce4454** | | | | 22-27 | 23-24-25 | 13 | 11 -- 12 | without sperm | not present | |
| **ce4455** | | | | 21-28 | 24-25-26-27 | 13 | 10--11--12 | without sperm | without sperm | |

| | | | | | | | | | | |
|---|---|---|---|---|---|---|---|---|---|---|
| **ce4456** | | | | 22-27 | 23-24-25 | 13 | left side: 11 right side: 10--11--12 | not present | not present | |
| **ce4457** | | | | 23-28 | 24-25-26 | 13 | 11 | without sperm | not present | |
| **ce4458** | | | | 22-27 | 23-24-25 | 13 | 11 --12 | without sperm | without sperm | |
| **ce4460** | | | | | 23-24-25 | 13 | | | | |
| **ce4573** | | | | 22-27 | 23-24-25 | 13 | 9--11--12 | not present | not present | |
| **ce4574** | | | | 22-27 | 23-24-25 | 13 | 11--12 | not present | not present | |
| **ce4575** | | | | 22-26 | 23-24-25 | 13 | 11--12 | not present | not present | |
| **UCM-LT number** | **Length (mm)** | **Dry weight (mg)** | **Number of segments** | **Clitellum** | **Tubercula pubertatis** | **Male pore** | **Seminal vesicles** | **Spermatechae** | **Spermidučal funnels** | **Observations** |
| **ce4587** | | | | 23-29 | 24-25-26-27 | 13 | 11--12 | without sperm | very small | |
| **ce4597** | | | | 22-27 | 23-24-25 | 13 | 9--11--12 | not present | not present | |
| **ce4725** | | | | | | 13 | | | | |
| **ce4726** | | | | 22-27 | 23-24-25 | 13 | 11--12 | without sperm | without sperm | |
| **ce4727** | | | | 21-25 | 22-23-24 | 13 | 11--12 | without sperm | not present | |
| **ce4728** | | | | 22-27 | 23-24-25 | 13 | 9-10-11-12 | not present | not present | |
| **ce4729** | | | | | | 13 | | | | |
| **ce4993** | | | | 22-27 | 23-24-25 | 13 | 11 -- 12 | not present | not present | |
| **ce5110** | | | | 22-27 | 23-24-25 | 13 | 10-11--12 | not present | very small | |
| **ce5111** | | | | 22-27 | 23-24-25 | 13 | 11--12 | not present | not present | |
| **ce5112** | | | | 22-27 | 23-24-25 | 13 | 9-10-11-12 | without sperm | without sperm | |
| **ce5248** | | | | | | 13 | | | | |
| **ce5685** | | | | | | 13 | | | | |
| **ce5711** | | | | | 23-24-25 | 13 | | | | |

| | | | | | | | | | | |
|---|---|---|---|---|---|---|---|---|---|---|
| **ce5712** | | | | 22-27 | 23-24-25 | 13 | 9-10-11-12 | not present | very small | |
| **ce5713** | | | | | | 13 | | | | |
| **ce5714** | | | | 22-27 | 23-24-25 | 13 | 9-10-11-12 | not present | not present | |
| **ce5715** | | | | 22-27 | 23-24-25 | 13 | 10--11--12 | not present | without sperm | |
| **ce5716** | | | | | 23-24-25 | 13 | | | | |
| **ce5717** | | | | 22-27 | 23-24-25 | 13 | 11--12 | not present | not present | |
| **30747** | 19.5 | 96.8 | 54 | 22-27 | 23-24-25 | 13 | 9-10-11-12 | without sperm | without sperm | spermatophore between 19 and 20 |
| **30748** | 26.5 | 148 | 79 | 22-26 | 22-23-24 | 13 | 9-10-11-12 | without sperm | without sperm | |
| **30749** | 27.5 | 136.6 | 86 | 23-28 | 24-25-26 | 13 | 9-10-11-12 | one on left side without sperm | without sperm | |
| **30750** | 30 | 133.7 | 85 | 23-28 | 24-25-26 | 13 | 9-10-11-12 | without sperm | not present | |
| **30751** | 34.5 | 192 | 90 | 22-27 | 23-24-25 | 13 | 9-10-11-12 | without sperm | not present | |
| **UCM-LT number** | **Length (mm)** | **Dry weight (mg)** | **Number of segments** | **Clitellum** | **Tubercula pubertatis** | **Male pore** | **Seminal vesicles** | **Spermatechae** | **Spermiducal funnels** | **Observations** |
| **30752** | | | | 22-27 | 23-24-25 | 13 | 11 | one on each side without sperm | not present | the earthworm was broken |
| **30753** | 32 | 150 | 84 | 22-27 | 23-24-25 | 13 | 9-10-11-12 | not present | not present | |
| **30754** | 23 | 87.5 | 107 | 22-28 | 23-24-25-26 | 13 | 9-10-11-12 | not present | not present | |
| **30755** | 30.5 | 167.1 | 90 | 22-26 | 23-24-25-26 | 13 | 11 -- 12 | not present | not present | |
| **30756** | 30 | 144.7 | 65 | 22-27 | 23-24-25 | 13 | 9-10-11-12 | not present | not present | |
| **30757** | 27 | 96.4 | 79 | 22-27 | 23-24-25 | 13 | 9-10-11-12 | not present | not present | |
| **30758** | 18 | 74.5 | 90 | | | | | | | inmature |
| **30650** | 12 | 50.3 | 88 | | | 13 | | | | semi-mature |
| **30651** | 19 | 81.7 | 80 | | 24-25-26 | 13 | | | | semi-mature |

| | | | | | | | | | | |
|---|---|---|---|---|---|---|---|---|---|---|
| **30652** | | | | | | 13 | | | | semi-mature, the earthworm was broken |
| **30653** | 15 | 77 | 91 | | | | | | | inmature |
| **30654** | 17 | 71 | 84 | | | | | | | inmature |
| **30655** | 11 | 30.8 | 86 | | | | | | | inmature |
| **30656** | 16 | 40.5 | 94 | | | | | | | inmature |
| **30657** | 12 | 20.8 | 77 | | | | | | | inmature |
| **30658** | 12.5 | 19.7 | 88 | | | | | | | inmature |
| **30663** | 15 | 24.3 | 60 | 22-27 | 23-24-25 | 13 | 9-10-11-12 | not present | without sperm | |
| **30667** | 16 | 39.1 | 84 | 22-27 | 23-24-25 | 13 | 9--11--12 | not present | not present | |
| **30668** | 18 | 44.2 | 77 | 22-27 | 23-24-25 | 13 | 11 -- 12 | not present | not present | |
| **30646** | 23 | 99.1 | 75 | 22-27 | 23-24-25 | 13 | 9--11--12 | not present | not present | |
| **30647** | 15 | 30.8 | 90 | | | | | | | inmature |
| **30648** | | | | 22-28 | 24-25-26 | 13 | 9--11--12 | without sperm | not present | the earthworm was broken |
| **30649** | 24 | 40.4 | 84 | | | 13 | | | | semi-mature |
| **30698** | 14.5 | 31.1 | 60 | 22-28 | 23-24-25-26 | 13 | 11 -- 12 | not present | not present | |
| **30699** | 25 | 59.7 | 82 | 22-28 | 23-24-25 | 13 | 9-10-11-12 | not present | not present | |
| **UCM-LT number** | **Length (mm)** | **Dry weight (mg)** | **Number of segments** | **Clitellum** | **Tubercula pubertatis** | **Male pore** | **Seminal vesicles** | **Spermatechae** | **Spermiducal funnels** | **Observations** |
| **30700** | | | | 22-27 | 23-24-25 | 13 | 9-10-11-12 | without sperm | not present | the earthworm was broken |
| **30701** | 15 | 31.3 | 67 | 22-27 | 23-24-25 | 13 | 9-10-11-12 | without sperm | not present | |
| **30702** | 10 | 22.7 | 94 | | | | | | | inmature |

| | | | | | | | | | | |
|---|---|---|---|---|---|---|---|---|---|---|
| **30703** | | | | 23-28 | 24-25-26 | 13 | 9-10-11-12 | without sperm | not present | the earthworm was broken |
| **30538** | 26 | 93.6 | 82 | 22-27 | 23-24-25 | 13 | 9-10-11-12 | without sperm | not present | |
| **30539** | 25 | 65.9 | 56 | 22-27 | 23-24-25 | 13 | 9-10-11-12 | without sperm | not present | |
| **30540** | 29.5 | 142.9 | 79 | 22-27 | 23-24-25 | 13 | 9-10-11-12 | without sperm | without sperm | |
| **30541** | 27 | 117.5 | 95 | 22-27 | 23-24-25 | 13 | 9-10-11-12 | not present | not present | |
| **30542** | 26.5 | 110.8 | 89 | 22-27 | 23-24-25 | 13 | 9-10-11-12 | not present | not present | |
| **30543** | 32 | 161.7 | 94 | 22-27 | 23-24-25 | 13 | 9-10-11-12 | not present | not present | |
| **30544** | 31 | 158.7 | 93 | 22-27 | 23-24-25 | 13 | 9-10-11-12 | without sperm | not present | |
| **30545** | | | | 22-27 | 23-24-25 | 13 | 9-10-11-12 | not present | not present | the earthworm was broken |
| **30546** | | | | 22-27 | 23-24-25 | 13 | 9-10-11-12 | not present | not present | the earthworm was broken |
| **30547** | 35 | 147.7 | 87 | 22-28 | 23-24-25-26 | 13 | 10--11-12 | not present | not present | |
| **30548** | | | | 22-27 | 23-24-25 | 15 | 9-10-11-12 | not present | not present | the earthworm was broken |
| **30599** | 19.5 | 44.3 | 60 | 22-27 | 23-24-25 | 13 | 10--11-12 | not present | not present | |
| **30719** | | | | 22-28 | 23-24-25-26 | 13 | 9-10-11-12 | not present | not present | the earthworm was broken |
| **30720** | 20 | 44.2 | 92 | 22-29 | 24-25-26 | 13 | 10--11--12 | not present | not present | |
| **30721** | 11 | 9.4 | 87 | | | | | | | |
| **30722** | 13 | 12.1 | 77 | | | | | | | |
| **30723** | | | | 22-27 | 23-24-25 | 13 | 9-10-11-12 | not present | not present | the earthworm was broken |
| **30724** | 18.5 | 34.9 | 81 | | 23-24-25 | 13 | | | | |
| **30725** | 6 | 6.9 | 74 | | | | | | | |

| | | | | | | | | | | |
|---|---|---|---|---|---|---|---|---|---|---|
| **30726** | | | | 22-27 | 23-24-25 | 13 | | | | broken |
| **UCM-LT number** | **Length (mm)** | **Dry weight (mg)** | **Number of segments** | **Clitellum** | **Tubercula pubertatis** | **Male pore** | **Seminal vesicles** | **Spermatechae** | **Spermiducal funnels** | **Observations** |
| **30727** | 11.5 | 7.8 | 83 | | | | 9-10--11-12 | not present | not present | |
| **30728** | | | | | | | | | | the earthworm was broken |
| **30729** | | | | | | 14 | | | | the earthworm was broken |
| **30788** | 13 | 8.8 | 77 | | | | | | | inmature |
| **30789** | 12 | 8.3 | 65 | | | | | | | inmature |
| **30790** | 9 | 5.1 | 60 | | | | | | | inmature |
| **30791** | 13 | 9.5 | 84 | | | | | | | inmature |
| **30792** | | | | | | | | | | inmature, the earthworm was broken |
| **30793** | | | | | | | | | | inmature, the earthworm was broken |
| **30794** | | | | | | | | | | inmature, the earthworm was broken |
| **30613** | 34 | 186 | 85 | 22-27 | 24-25-26 | 12 | 9-10-11-12 | not present | not present | |
| **30614** | 29.5 | 131.7 | 69 | 22-27 | 23-24-25 | 13 | 9-10-11-12 | not present | not present | |
| **30615** | 12 | 31.1 | 87 | | | | | | | |
| **30616** | | | | | | | | | | the earthworm was broken |
| **30617** | 23 | 68.4 | 83 | | | | | | | |
| **30618** | 20 | 33.1 | 96 | | | | | | | |

| | | | | | | | | | | |
|---|---|---|---|---|---|---|---|---|---|---|
| **30619** | | | | | | | | | | the earthworm was broken |
| **30620** | 22.5 | 83.1 | 77 | | | | | | | |
| **30621** | 19 | 75.3 | 82 | | | | | | | |
| **30622** | 9 | 10.3 | 71 | | | | | | | |
| **30623** | | | | | | | | | | the earthworm was broken |
| **30624** | 32 | 165.9 | 94 | 23-30 | 24-25-26-27 | 13 | 9--11--12 | not present | not present | |
| **30625** | 25 | 94.8 | 61 | 18-23 | 20-21-22 | 8 | not present | not present | not present | |
| **UCM-LT number** | **Length (mm)** | **Dry weight (mg)** | **Number of segments** | **Clitellum** | **Tubercula pubertatis** | **Male pore** | **Seminal vesicles** | **Spermatechae** | **Spermiducal funnels** | **Observations** |
| **30626** | 28 | 127.1 | 91 | 22-28 | 23-24-25 | 13 | 11 -- 12 | not present | not present | |
| **30627** | 21.5 | 77.4 | 69 | 22-27 | 23-24-25 | 13 | 9-10-11-12 | not present | not present | |
| **30628** | 36 | 110.7 | 81 | 22-28 | 23-24-25 | 13 | 9-10-11-12 | not present | not present | |
| **30629** | 27.5 | 102.7 | 77 | 22-27 | 23-24-25 | 13 | 10--11--12 | not present | not present | |
| **30630** | 22 | 88.9 | 66 | 22-27 | 23-24-25 | 13 | 9--11--12 | not present | not present | |
| **30631** | 27 | 107.9 | 72 | 22-27 | 23-24-25 | 13 | 9-10-11-12 | not present | not present | |
| **30632** | 29 | 110.2 | 100 | 22-28 | 23-24-25-26 | 13 | 11 -- 12 | not present | not present | |
| **30633** | 25 | 104.7 | 98 | 22-27 | 23-24-25 | 13 | 9--11--12 | not present | not present | |
| **30562** | 26 | 85.7 | 74 | 22-27 | 23-24-25 | 13 | 11 -- 12 | not present | not present | |
| **30563** | 17 | 43.3 | 73 | | 23-24-25 | 13 | | not present | not present | the earthworm was broken |
| **30564** | | | | | | 13 | | | | semi-mature, the earthworm was broken |

| | | | | | | | | | | |
|---|---|---|---|---|---|---|---|---|---|---|
| **30565** | | | | 22-27 | 23-24-25 | 13 | 9-10-11-12 | | | semi-mature, the earthworm was broken |
| **30566** | | | | | | | | without sperm | without sperm | |
| **30567** | 24.5 | 79.8 | 64 | 22-27 | 23-24-25 | 13 | 9-10-11-12 | | | |
| **30568** | 21 | 71.3 | 52 | 22-27 | 23-24-25 | 13 | 9-10-11-12 | without sperm | not present | |
| **30569** | 20 | 38.6 | 82 | | | | | | | |
| **30570** | 16 | 22.4 | 85 | | | | | | | |
| **30571** | 22 | 65 | 75 | 22-27 | 23-24-25 | 13 | 9-10-11-12 | without sperm | without sperm | |
| **30572** | | | | 22-27 | 23-24-25-26 | 13 | 9-10-11-12 | not present | not present | the earthworm was broken |
| **30573** | 20 | 27.9 | 79 | | | 13 | | | | |
| **30574** | 22 | 112.4 | 62 | 22-28 | 24-25-26 | 13 | 9-10-11-12 | not present | not present | |
| **30575** | | | | | 23-24-25 | 13 | | | | the earthworm was broken |
| **30576** | 21 | 67.3 | 68 | 22-27 | 24-25-26 | 13 | 11 -- 12 | not present | not present | |
| **30577** | 21.5 | 65.8 | 77 | | | | | | | |
| **UCM-LT number** | **Length (mm)** | **Dry weight (mg)** | **Number of segments** | **Clitellum** | **Tubercula pubertatis** | **Male pore** | **Seminal vesicles** | **Spermatechae** | **Spermiducal funnels** | **Observations** |
| **30578** | | | | 22-27 | 23-24-25 | 13 | 9 -- 10 | not present | not present | the earthworm was broken |
| **30603** | | | | | | | | | | the earthworm was broken |
| **30604** | 33 | 187.7 | 94 | | 22-27 | 13 | 9-10-11-12 | not present | not present | |
| **30605** | 39 | 157.1 | 90 | | 22-27 | 13 | 9-10-11-12 | not present | not present | |
| **30606** | 42 | 260.3 | 76 | | 22-27 | 13 | 9-10-11-12 | without sperm | without sperm | the earthworm was broken |

| | | | | | | | | | | |
|---|---|---|---|---|---|---|---|---|---|---|
| **30609** | | | | | 22-27 | 13 | 9-10-11-12 | without sperm | without sperm | the earthworm was broken |
| **30610** | | | | | | 13 | | | | |
| **30611** | 33.5 | 134.5 | 74 | | 22-27 | 13 | 11 -- 12 | not present | not present | |
| **30612** | 32 | 121.4 | 81 | | 22-27 | 13 | 10--11--12 | not present | not present | |
| **30591** | 25 | 118 | 75 | 22-27 | 23-24-25 | 13 | 9-10-11-12 | not present | not present | |
| **30592** | 31 | 122.6 | 71 | 22-27 | 23-24-25 | 13 | 10--11-12 | not present | not present | |
| **30593** | | | | | | 13 | | | | the earthworm was broken |
| **30594** | 25 | 66.7 | 87 | | 23-24-25 | 13 | | | | |
| **30595** | | | | 22-27 | 23-24-25 | 13 | 9-10-11--12 | not present | not present | the earthworm was broken |
| **30596** | 31 | 120.2 | 88 | 22-27 | 23-24-25 | 13 | 11 -- 12 | not present | not present | |
| **30597** | | | | 22-27 | 23-24-25 | 13 | 9-10-11--12 | not present | not present | the earthworm was broken |
| **30598** | 13 | 13 | 82 | | | | | | | |
| **30580** | | | | | | 13 | | | | the earthworm was broken |
| **30581** | | | | 22-27 | 23-24-25 | 13 | 9-10-11-12 | without sperm | without sperm | the earthworm was broken |
| **30582** | 18 | 88.7 | 39 | 22-27 | 23-24-25 | 13 | 9-10-11-12 | without sperm | without sperm | |
| **30583** | 22 | 82.2 | 73 | 22-27 | 23-24-25-26 | 13 | 9-10-11-12 | without sperm | without sperm | |
| **30584** | | | | 22-27 | 23-24-25 | 13 | 11 -- 12 | not present | not present | the earthworm was broken |
| **30585** | 15 | 21.9 | 93 | | | | | | | |

| UCM-LT number | Length (mm) | Dry weight (mg) | Number of segments | Clitellum | Tubercula pubertatis | Male pore | Seminal vesicles | Spermatechae | Spermiducal funnels | Observations |
|---|---|---|---|---|---|---|---|---|---|---|
| **30587** | | | | | | 13 | | | | the earthworm was broken |
| **30588** | 14.5 | 20.3 | 88 | | | | | | | |
| **30589** | 11 | 11.1 | 80 | | | | | | | |
| **30590** | 9 | 5.3 g | 77 | | | | | | | |
| **30761** | 22 | 63.1 | 93 | 22-27 | 24-25-26 | 13 | 9-10-11-12 | not present | not present | |
| **30762** | 22 | 64.8 | 87 | 22-26 | 23-24-25 | 13 | 9-10-11-12 | not present | not present | |
| **30763** | | | | | | | | | | the earthworm was broken |
| **30765** | 18 | 49.6 | 59 | 22-27 | 23-24-25 | 13 | 11 -- 12 | not present | not present | |
| **30766** | 20 | 38.2 | 88 | | | 13 | | | | |
| **30767** | 14 | 25.4 | 56 | | | | | | | |
| **30768** | 21.5 | 53.9 | 94 | | 23-24-25 | 13 | | | | |
| **30769** | | | | 22-27 | 23-24-25 | 13 | 9-10-11-12 | not present | not present | the earthworm was broken |
| **30770** | 18 | 25.4 | 74 | 22-27 | 23-24-25 | 13 | 9-10-11-12 | not present | not present | |
| **30771** | 20 | 77.8 | 90 | 22-28 | 23-24-25-26 | 13 | 10--11--12 | not present | not present | |
| **30772** | 20 | 75.4 | 87 | | 23-24-25 | 13 | | | | |
| **30773** | | | | 22-26 | 23-24-25 | 13 | 9-10-11-12 | not present | not present | the earthworm was broken |
| **30774** | 20 | 76.8 | 88 | 22-27 | 23-24-25 | 13 | 9-10-11-12 | not present | not present | |
| **30775** | 15 | 30.4 | 90 | | | | | | | |

| | | | | | | | | | | |
|---|---|---|---|---|---|---|---|---|---|---|
| **30776** | | | | | | | | | | inmature, the earthworm was broken |
| **30777** | 19 | 65.3 | 84 | | 23-24-25 | 13 | | | | |
| **30778** | | | | | 23-24-25 | 13 | | | | the earthworm was broken |
| **30843** | | | | | | | | | | inmature, the earthworm was broken |
| **30844** | 22 | 76.2 | 90 | | 23-24-25 | 13 | | | | semi-mature |
| **30845** | 19 | 32.1 | 83 | | | | | | | |
| **UCM-LT number** | **Length (mm)** | **Dry weight (mg)** | **Number of segments** | **Clitellum** | **Tubercula pubertatis** | **Male pore** | **Seminal vesicles** | **Spermatechae** | **Spermiducal funnels** | **Observations** |
| **30846** | 37 | 160.2 | 89 | 23-28 | 24-25-26 | 13 | 11 -- 12 | one on each side without sperm | without sperm | |
| **30847** | 20 | 63.7 | 77 | | 23-24-25 | 13 | | | | |
| **30848** | | | | | 23-24-25 | 13 | | | | the earthworm was broken |
| **30849** | | | | | | | | | | inmature, the earthworm was broken |
| **30850** | 18 | 30.3 | 91 | | | 13 | | | | |
| **30851** | | | | | | | | | | |
| **30854** | 20 | 59.3 | 66 | 22-27 | 23-24-25 | 13 | 9-10-11-12 | one on each side without sperm | without sperm | |
| **30856** | 4 | 30 | 57 | | | | | | | inmature |
| **30857** | 5 | 32 | 61 | | | | | | | inmature |
| **30858** | 17 | | | | | | | | | |
| **30859** | 20 | 150 | 62 | 24-27 | 25-26-27 | 13 | 11 -- 12 | not present | without sperm | |

| | | | | | | | | | | |
|---|---|---|---|---|---|---|---|---|---|---|
| **30860** | | | | | | | | | | |
| **30861** | 21 | 45.6 | 65 | 22-26 | 22-23-24-25-26 | 13 | 9-10-11-12 | not present | not present | |
| **30862** | 24 | 51.5 | 83 | 22-27 | 23-24-25 | 13 | 10--11--12 | not present | not present | |
| **30863** | | | | 22-26 | 23-24-25 | 13 | 11 -- 12 | not present | without sperm | the earthworm was broken |
| **30864** | | | | 22-26 | 23-24-25 | 13 | 11 | not present | not present | the earthworm was broken |
| **30865** | | | | 21-26 | 22-23-24 | 10 | 11 -- 12 | not present | not present | the earthworm was broken |
| **30866** | | | | | 21-22-23 | 10 | 8--9--10 | not present | without sperm | the earthworm was broken |
| **30867** | | | | 22-26 | 23-24-25 | 13 | 10--11--12 | without sperm | without sperm | the earthworm was broken |
| **12** | 14 | 12.3 | 72 | | | | | | | inmature |
| **30869** | 9 | 9.8 | 81 | | | | | | | inmature |
| **30870** | 10 | 11.5 | | | | | | | | inmature |
| **30871** | | | | | | | | | | |
| **UCM-LT number** | **Length (mm)** | **Dry weight (mg)** | **Number of segments** | **Clitellum** | **Tubercula pubertatis** | **Male pore** | **Seminal vesicles** | **Spermatechae** | **Spermiducal funnels** | **Observations** |
| **30872** | | | | | | | | | | inmature, the earthworm was broken |
| **30873** | 6 | 3.2 | 65 | | | | | | | |
| **30876** | | | | | | | | | | inmature, the earthworm was broken |
| **30877** | 21 | 43.5 | 67 | | 23-24-25 | 13 | | | | semi-mature |
| **30878** | 8 | 6.7 | 73 | | | | | | | |

| | | | | | | | | | | |
|---|---|---|---|---|---|---|---|---|---|---|
| **30879** | | | | | | | | | | inmature, the earthworm was broken |
| **30880** | | | | | | 13 | | | | semi-mature, the earthworm was broken |
| **30883** | 20 | 35.2 | 77 | | | | | | | inmature |
| **30884** | | | | 21-26 | 22-23-24 | 13 | 9-10-11-12 | without sperm | without sperm | the earthworm was broken |
| **30885** | 20 | 47.4 | 69 | 22-27 | 23-24-25-26 | 13 | 10 -- 11 | without sperm | without sperm | |
| **30886** | 10 | 11.3 | 62 | | | | | | | inmature |
| **30887** | 18 | 21.1 | 65 | | | | | | | inmature |
| **30888** | 20.5 | 60.6 | 71 | 21-26 | 22-25 | 13 | 9-10-11-12 | not present | not present | |
| **30889** | | | | | 23-24-25 | 13 | | | | semi-mature |
| **30890** | 19.5 | 33.1 | 86 | 22-28 | 24-25-26 | 13 | 9-10-11-12 | not present | not present | the earthworm was broken |
| **30891** | | | | | | | | | | the earthworm was broken |
| **30892** | 14 | 31.2 | 74 | | | | | | | inmature |
| **30894** | | | | | | 13 | | | | semi-mature, the earthworm was broken |
| **30896** | 29 | 67..7 | 87 | 22-28 | 23-24-25 | 12 | 10--11--12 | not present | without sperm | |
| **30897** | | | | | | | | | | inmature and the earthworm was broken |
| **30898** | | | | | | | | | | inmature and the earthworm was broken |
| **30899** | | | | | | | | | | inmature and the earthworm was broken |

| UCM-LT number | Length (mm) | Dry weight (mg) | Number of segments | Clitellum | Tubercula pubertatis | Male pore | Seminal vesicles | Spermatechae | Spermiducal funnels | Observations |
|---|---|---|---|---|---|---|---|---|---|---|
| **30900** | 24 | 55.9 | 82 | 22-28 | 23-24-25 | 13 | 9-10-11-12 | without sperm | without sperm | |
| **30901** | | | | 21-26 | 22-23-24-25 | 12 | 9-10-11-12 | not present | not present | the earthworm was broken |
| **30902** | 15 | 34.2 | 69 | | | | | | | inmature |
| **30903** | 29 | 79.2 | 89 | | | 13 | | | | semi-mature |
| **30904** | 25 | 56.7 | 84 | 21-27 | 23-24-25 | 13 | 10--11--12 | without sperm | without sperm | |
| **30905** | 19.5 | 34.2 | 91 | | | 13 | | | | semi-mature |
| **30906** | 22.5 | 46.3 | 97 | 21-28 | 22-23-24-25-26-1n27 | 11 | 9-10-11-12 | without sperm | without sperm | |
| **30907** | 17 | 27.1 | 87 | | | | | | | inmature |
| **30908** | | | | 22-28 | 23-24-25 | 13 | 11 -- 12 | one on each side without sperm | without sperm | the earthworm was broken |
| **30909** | | | | 20-26 | 22-23-24-25 | 13 | 9-10-11-12 | not present | not present | the earthworm was broken |
| **30910** | | | | | | | | | | the earthworm was broken |
| **30911** | 23 | 58.5 | 91 | | 23-24-25 | 13 | | | | semi-mature |
| **30912** | | | | | | | | | | |
| **30913** | 18 | 32.3 | 82 | | | | | | | inmature |
| **30914** | | | | | | | | | | |
| **30915** | 29 | 117.9 | 85 | 22-27 | 23-24-25-1n26 | 13 | 11 -- 12 | not present | not present | |
| **30916** | 26.5 | 101.6 | 90 | 22-28 | 23-24-25-1n26 | 13 | 10--11--12 | one on each side without sperm | without sperm | |

| | | | | | | | | | | |
|---|---|---|---|---|---|---|---|---|---|---|
| **30917** | 20 | 36.4 | 87 | | | 13 | | | | semi-mature |
| **30918** | 19 | 30.5 | 77 | | | | | | | inmature |
| **30919** | | | | | | | | | | |
| **30920** | 29 | 123.5 | 88 | 22-27 | 23-24-25-1n26 | 13 | 11 -- 12 | without sperm | not present | |
| **30921** | 14.5 | 19.8 | 83 | | | | | | | inmature |
| **30922** | 18 | 38.1 | 79 | | | | | | | inmature |
| **UCM-LT number** | **Length (mm)** | **Dry weight (mg)** | **Number of segments** | **Clitellum** | **Tubercula pubertatis** | **Male pore** | **Seminal vesicles** | **Spermatechae** | **Spermiducal funnels** | **Observations** |
| **30924** | | | | | | | | | | |
| **30925** | | | | | | | | | | |
| **30926** | 27 | 116.9 | 82 | 21-26 | 22-23-24 | 13 | 11 -- 12 | one on each side without sperm | not present | |
| **30927** | | | | 22-27 | 23-24-25-26 | 13 | 11 -- 12 | not present | not present | the earthworm was broken |
| **30928** | | | | | | | | | | |
| **30929** | 22 | 73.5 | 85 | 22-27 | 23-24-25 | 13 | 9-10-11-12 | one on each side without sperm | without sperm | |
| **30930** | | | | | | | | | | |
| **30931** | | | | | | | | | | |
| **30932** | 18 | 41.3 | 72 | 22-28 | 23-24-25-26 | 13 | 11 -- 12 | not present | not present | |
| **30933** | 16 | 39.5 | 57 | | | | | | | inmature |
| **30934** | 15 | 33.7 | | | | 13 | | | | semi-mature |
| **30935** | | | | | | | | | | |

| | | | | | | | | | | |
|---|---|---|---|---|---|---|---|---|---|---|
| **30936** | | | | 21-26 | 23-24-25 | 12 | 9-10-11-12 | without sperm | not present | the earthworm was broken |
| **30937** | | | | 22-28 | 23-24-25-26 | 13 | 9-10-11-12 | not present | not present | the earthworm was broken |
| **30938** | 15 | 43.1 | 53 | 22-27 | 23-24-25 | 13 | 10-- 11 | not present | not present | |
| **30939** | | | | | 23-24-25 | 13 | | | | semi-mature, the earthworm was broken |
| **30940** | | | | | | | | | | |
| **30941** | 18 | 39.8 | 82 | 21-27 | 23-24-25 | 13 | 11 -- 12 | not present | not present | |
| **30946** | 19 | 43.8 | 77 | | | | | | | |
| **30947** | | | | | | | | | | inmature, the eathworm was broken |
| **30948** | | | | | | | | | | inmature, the eathworm was broken |
| **30951** | | | | | | 13 | | | | semi-mature, the earthworm was broken |
| **30953** | 24 | 60 | 86 | 22-27 | 23-24-25 | 13 | 9-10-11-12 | one on right side without sperm | without sperm | |
| **UCM-LT number** | **Length (mm)** | **Dry weight (mg)** | **Number of segments** | **Clitellum** | **Tubercula pubertatis** | **Male pore** | **Seminal vesicles** | **Spermatechae** | **Spermiducal funnels** | **Observations** |
| **30960** | | | | | | | | | | inmature, the eathworm was broken |
| **30991** | 22.5 | 63.1 | 77 | | 23-24-25 | 13 | | | | |
| **30993** | 21.5 | 62.7 | 82 | | 23-24-25 | 13 | | | | |
| **30995** | | | | | | | | | | inmature, the earthworm was broken |
| **31002** | | | | | | | | | | |

| | | | | | | | | | | |
|---|---|---|---|---|---|---|---|---|---|---|
| **31008** | 20 | 64.3 | 77 | | 23-24-25 | 13 | | | | |
| **30818** | 21.5 | 55.3 | 93 | 22-27 | 23-24-25 | 13 | 11--12 | not present | not present | |
| **30811** | 19.5 | 43.6 | 85 | | 23-24-25 | 13 | | | | semi-mature |
| **30823** | 17 | 33.9 | 74 | | 23-24-25 | 13 | | | | semi-mature |
| **30814** | 29 | 97 | 87 | 22-27 | 23-24-25 | 13 | 11---12 | not present | not present | |
| **30820** | 14 | 16.5 | 77 | | | | | | | inmature |
| **30815** | 20 | 65.6 | 61 | 22-27 | 23-24-25 | 13 | 9--11--12 | not present | without sperm | |
| **30817** | 23 | 72.3 | 85 | | 23-24-25 | 13 | | | | semi-mature |
| **30816** | 24.5 | 84.1 | 101 | 22-28 | 24-25-26 | 13 | 11 -- 12 | not present | without sperm | |
| **30812** | 35 | 138.7 | 87 | 22-27 | 23-24-25 | 13 | 11 -- 12 | not present | not present | |
| **30813** | | | | 22-27 | 23-24-25 | 13 | 11 -- 12 | not present | not present | the earthworm was broken |
| **30838** | 23.5 | 63.7 | 91 | 22-28 | 24-25-26 | 13 | 9-10-11-12 | without sperm | without sperm | |
| **30837** | 17 | 55.6 | 58 | | 23-24-25 | 13 | | | | semi-mature |
| **30841** | | | | | | | | | | inmature, the earthworm was broken |
| **30842** | | | | | | 13 | | | | semi-mature, the earthworm was broken |
| **30836** | | | | | | | | | | inmature, the earthworm was broken |
| **30839** | 11 | 33 | 78 | | | 13 | | | | semi-mature |
| **30840** | | | | | | | | | | the earthworm was broken |
| **30796** | | | | | 23-24-25 | 13 | | | | semi-mature, broken |

| UCM-LT number | Length (mm) | Dry weight (mg) | Number of segments | Clitellum | Tubercula pubertatis | Male pore | Seminal vesicles | Spermatechae | Spermiducal funnels | Observations |
|---|---|---|---|---|---|---|---|---|---|---|
| **30797** | 18 | 28.8 | 82 | | | | | | | inmature |
| **30795** | 24 | 68.4 | 93 | 22-27 | 23-24-25 | 13 | 11--12 | without sperm | without sperm | |
| **30800** | 13 | 17.8 | 77 | | | | | | | inmature |
| **30798** | | | | 22-27 | 23-24-25 | 13 | 9--11--12 | without sperm | without sperm | the earthworm was broken |
| **30802** | | | | | 23-24-25 | 13 | | | | semi-mature, the earthworm was broken |
| **30804** | 19 | 25 | 81 | | | | | | | inmature |
| **30801** | 16 | 21.3 | 78 | | | | | | | inmature |
| **30803** | | | | | | 13 | | | | semi-mature, the earthworm was broken |
| **30808** | 15 | 19.9 | 66 | | | | | | | inmature |
| **30670** | 24 | 111.5 | 81 | 22-27 | 23-24-25 | 13 | 9--11--12 | not present | not present | spermatophore between 19 and 20 |
| **30692** | 25 | 113.8 | 93 | 22-27 | 23-24-25 | 13 | 10--11--12 | without sperm | without sperm | |
| **30687** | 18 | 100.7 | 88 | 22-27 | 23-24-25 | 13 | 9-10-11-12 | not present | not present | |
| **30696** | 30 | 114 | 89 | 22-27 | 23-24-25 | 13 | 9--11--12 | without sperm | without sperm | |
| **30694** | 12 | 76.8 | 57 | 22-27 | 23-24-25 | 13 | 9-10-11-12 | not present | not present | |
| **30688** | 21 | 74.4 | 84 | 22-27 | 23-24-25 | 13 | 9-10-11-12 | without sperm | without sperm | |
| **30686** | 20 | 86.4 | 64 | 22-27 | 23-24-25 | 13 | 9-10-11-12 | without sperm | without sperm | |
| **30693** | 26 | 109.7 | 87 | 22-28 | 24-25-26 | 13 | 9-10-11-12 | without sperm | without sperm | |
| **30685** | | | | 22-27 | 23-24-25 | 13 | 9-10-11-12 | not present | not present | the earthworm was broken |

| | | | | | | | | | | |
|---|---|---|---|---|---|---|---|---|---|---|
| **30684** | 18 | 61.2 | 65 | 22-27 | 23-24-25 | 13 | 9-10-11-12 | without sperm | without sperm | |
| **30736** | 32 | 160.4 | | 22-27 | 23-24-25 | 13 | | | | the earthworm was broken |
| **30737** | 28 | 157.4 | 94 | 22-27 | 23-24-25 | 13 | 11 -- 12 | without sperm | not present | |
| **30730** | 31 | 162.3 | 87 | 22-27 | 23-24-25 | 13 | 11 -- 12 | without sperm | without sperm | |
| **30738** | | | | 22-26 | 23-24-25 | 13 | 10--11--12 | without sperm | without sperm | the earthworm was broken |
| **30739** | 23 | 75.1 | 87 | 22-27 | 23-24-25-26 | 13 | 9-10-11-12 | without sperm | without sperm | |
| **UCM-LT number** | **Length (mm)** | **Dry weight (mg)** | **Number of segments** | **Clitellum** | **Tubercula pubertatis** | **Male pore** | **Seminal vesicles** | **Spermatechae** | **Spermiducal funnels** | **Observations** |
| **30741** | 22.5 | 68.8 | 77 | | 23-24-25 | 13 | | | | semi-mature |
| **30735** | 19.5 | 44.7 | 82 | | | | | | | inmature |
| **30734** | 23 | 82.1 g | 81 | 22-27 | 23-24-25 | 13 | 9-10-11-12 | not present | not present | |
| **30733** | 19 | 60.1 | 89 | | 23-24-25 | 13 | | | | semi-mature |
| **30732** | 13 | 15.6 | 65 | | | | | | | inmature |
| **30716** | 24 | 102.3 | 88 | 22-26 | 23-24-25 | 13 | 9-10--11-12 | without sperm | without sperm | |
| **30713** | | | | 22-27 | 23-24-25 | 13 | 9-10-11-12 | not present | not present | the earthworm was broken |
| **30714** | 22.5 | 111.1 | 79 | 22-26 | 23-24-25 | 13 | 9-10-11-12 | not present | not present | |
| **30715** | 22 | 109 | 84 | 22-27 | 23/24/25 | 13 | | | | |
| **30717** | | | | | | | | | | inmature, the earthworm was broken |
| **31023** | 16 | 30 | 74 | | | | | | | inmature |
| **31024** | 14.5 | 11.2 | 77 | | | | | | | inmature |
| **31025** | 13 | 20 | | | | | | | | |

| | | | | | | | | | | |
|---|---|---|---|---|---|---|---|---|---|---|
| **31026** | | | | | | | | | | inmature, the earthworm was broken |
| **31027** | | | | | | | | | | inmature, the earthworm was broken |
| **31028** | 8 | 11.2 | 71 | | | | | | | inmature |
| **31029** | | | | | | | | | | inmature, the earthworm was broken |
| **31030** | 13 | 20.1 | 60 | | | | | | | inmature |
| **31031** | | | | 22-28 | 23-27 | 13 | 9-10-11-12 | not present | not present | the earthworm was broken |
| **31032** | | | | 21-27 | 23-24-25 | 13 | 9-10-11-12 | not present | not present | the earthworm was broken |
| **31033** | | | | | | | | | | the earthworm was broken |
| **31034** | 20.5 | 57.6 | 63 | 22-28 | 23-24-25-1n26 | 13 | 9-10-11-12 | not present | not present | |
| **31038** | 30 | 118.8 | | | | 13 | 9-10-11-12 | not present | not present | the earthworm was broken |
| **UCM-LT number** | **Length (mm)** | **Dry weight (mg)** | **Number of segments** | **Clitellum** | **Tubercula pubertatis** | **Male pore** | **Seminal vesicles** | **Spermatechae** | **Spermiducal funnels** | **Observations** |
| **31039** | 14 | 21.1 | 60 | | | | | | | inmature |
| **31040** | 13 | 18.1 | 72 | | | | | | | inmature |
| **31041** | 27 | 110.5 | 79 | | | | | | | inmature |
| **31043** | 15 | 20.8 | 64 | | | | | | | inmature |
| **31044** | 12 | 10.5 | 70 | | | | | | | inmature |
| **31045** | 25 | 111.5 | 69 | 22-27 | 23-24-25-26 | 13 | 10--11--12 | without sperm | without sperm | |
| **31046** | 17.5 | 50.8 | 66 | 22-27 | 23-24-25 | 13 | 10--11--12 | without sperm | without sperm | |

| | | | | | | | | | | |
|---|---|---|---|---|---|---|---|---|---|---|
| **31047** | 19.5 | 30.3 | | | | | | | | inmature |
| **31048** | | | | | | | | | | |
| **31049** | 10 | 108 | 73 | | | | | | | inmature |
| **31050** | 9 | 10.9 | 71 | | | | | | | inmature |
| **31051** | 24 | 90.8 | 59 | 22-27 | 23-24-25-1n26 | 13 | 9-10-11-12 | not present | without sperm | |
| **31052** | 16 | 10.5 | 71 | | | | | | | inmature |
| **31053** | | | | | | | | | | |
| **31054** | 17.5 | 54.8 | 82 | | | | | | | inmature |
| **31055** | | | | | | | | | | inmature, the earthworm was broken |
| **31056** | 9 | 10.1 | 67 | | | | | | | inmature |
| **31057** | | | | | | | | | | inmature, the earthworm was broken |
| **31058** | 6 | 7.4 | 65 | | | | | | | inmature |
| **31059** | 4 | 6.1 | 69 | | | | | | | inmature |
| **31060** | 12 | 20.4 | 65 | | | | | | | inmature |
| **31061** | 9 | 10.2 | 63 | | | | | | | inmature |
| **31062** | 9.5 | 11.4 | 77 | | | | | | | inmature |
| **31063** | 11 | 12.5 | 75 | | | | | | | inmature |
| **31064** | 10 | 16.5 | 67 | | | | | | | inmature |
| **UCM-LT number** | **Length (mm)** | **Dry weight (mg)** | **Number of segments** | **Clitellum** | **Tubercula pubertatis** | **Male pore** | **Seminal vesicles** | **Spermatechae** | **Spermidual funnels** | **Observations** |
| **31065** | 0.9 | 5.2 | 87 | | | | | | | inmature |
| **31066** | 0.8 | 11.9 | | | | | | | | inmature |

| | | | | | | | | | | |
|---|---|---|---|---|---|---|---|---|---|---|
| **31067** | | | | 22-28 | 24-25-26-27 | 13 | 10--11--12 | not present | not present | the earthworm was broken |
| **31068** | | | | 23-28 | 24-25-26 | 13 | 11 -- 12 | not present | not present | the earthworm was broken |
| **31069** | 0.8 | 11.3 | 87 | | | | | | | inmature |
| **31070** | 11 | 12.5 | 88 | | | | | | | inmature |
| **31071** | 23 | 50.9 | 83 | | | | | | | inmature |
| **31072** | | | | | | | | | | the earthworm was broken and nmature |
| **31073** | | | | | | | | | | the earthworm was broken and inmature |
| **31074** | 16 | 25.6 | 78 | | | | | | | inmature |
| **31075** | | | | | | | | | | the earthworm was broken and inmature |
| **31076** | | | | | | | | | | |
| **31077** | | | | | | | | | | the earthworm was broken and inmature |
| **31078** | 0.7 | 3.5 | 82 | | | | | | | inmature |
| **31082** | 26 | 90.6 | 86 | 22-28 | 23-24-25 | 13 | 8-9-10-11 | without sperm | not present | |
| **31083** | 19 | 45.5 | 85 | | | | | | | inmature |
| **31084** | 21 | 50.5 | 77 | | 23-24-25 | | | | | semi-mature |
| **31085** | 8 | 10.3 | 65 | | | | | | | inmature |
| **31086** | 17 | 30.5 | 71 | | | | | | | inmature |
| **31087** | 15 | 33.6 | 74 | | | | | | | inmature |
| **31088** | | | | 23-28 | 24-25-26 | 13 | 9-10-11-12 | one on each side without sperm | without sperm | the earthworm was broken |

| | | | | | | | | | | |
|---|---|---|---|---|---|---|---|---|---|---|
| **31089** | 20 | 43.2 | 82 | | | | | | | inmature |
| **31090** | | | | | | | | | | the earthworm was broken and inmature |

| **UCM-LT number** | **Length (mm)** | **Dry weight (mg)** | **Number of segments** | **Clitellum** | **Tubercula pubertatis** | **Male pore** | **Seminal vesicles** | **Spermatechae** | **Spermiducal funnels** | **Observations** |
|---|---|---|---|---|---|---|---|---|---|---|
| **31091** | | | | 22-27 | 24-25-26 | 13 | 9-10-11-12 | not present | not present | the earthworm was broken |
| **31092** | 21 | 65.2 | 91 | 23-27 | 24-25-26 | 13 | 9-10-11-12 | without sperm | without sperm | |
| **31093** | 22 | 74.2 | 73 | 23-28 | 24-25-26 | 15 | 9-10-11-12 | without sperm | not present | |
| **31094** | 40 | 170.2 | 102 | 22-26 | 23-24-25 | 13 | 10--11--12 | not present | not present | |
| **31095** | 34 | 141.7 | 89 | 22-28 | 1n22-1n28 | 13 | 11 -- 12 | not present | without sperm | |
| **31096** | 30 | 111.4 | 93 | 22-28 | 24-25-26 | 13 | 10--11--12 | not present | not present | |
| **31097** | 26 | 95.6 | 80 | 23-27 | 24-25-26 | 13 | 9-10-11-12 | not present | not present | |
| **31098** | 21.5 | 78.4 | 51 | 22-26 | 23-24 | 13 | 10--11--12 | not present | not present | |
| **31099** | 33 | 120.3 | 88 | 22-27 | 23-24-25-26 | 13 | 11 -- 12 | not present | not present | |
| **31100** | 10 | 22.2 | 77 | | | | | | | inmature |
| **31101** | 0.8 | 24.5 | 74 | | | | | | | inmature |
| **31102** | 17 | 66.9 | 78 | | 24-25-26 | 13 | | | | semi-mature |
| **31103** | 29 | 71.8 | | | | 13 | 9-10-11-12 | not present | not present | |
| **31104** | 27 | 74.5 | 80 | 22-27 | 23-24-25 | 13 | 11 -- 12 | without sperm | not present | |
| **31105** | | | | 23-28 | 24-25-26 | 13 | 9-10-11-12 | not present | not present | the earthworm was broken |
| **31106** | 33 | 125.7 | 74 | 22-27 | 23-24-25 | 13 | 9-10-11-12 | without sperm | without sperm | |
| **31107** | 19.5 | 66.5 | 60 | 23-28 | 24-25-26 | 13 | 11 -- 12 | not present | not present | |
| **31108** | 20 | 58.4 | 87 | | | | | | | inmature |

| | | | | | | | | | | |
|---|---|---|---|---|---|---|---|---|---|---|
| **31109** | 14.5 | 10.9 | 83 | | | | | | | inmature |
| **31110** | 21 | 51 | | | | | | | | inmature |
| **31111** | 32 | 141.2 | 93 | 23-28 | 24-25-26 | 13 | 10--11--12 | not present | not present | |
| **31112** | | | | 23-28 | 24-25-26 | 13 | 10--11--12 | not present | not present | the earthworm was broken |
| **31113** | 23 | 55.3 | 94 | 22-27 | 13-24-25 | 13 | 11 -- 12 | without sperm | without sperm | the earthworm was broken and inmature |
| **31114** | | | | | | | | | | inmature |
| **31115** | 10 | 11.6 | 79 | | | | | | | inmature |
| **UCM-LT number** | **Length (mm)** | **Dry weight (mg)** | **Number of segments** | **Clitellum** | **Tubercula pubertatis** | **Male pore** | **Seminal vesicles** | **Spermatechae** | **Spermiducal funnels** | **Observations** |
| **31116** | 22 | 80.5 | 86 | 22-27 | 23-24-25 | 13 | 9-10-11-12 | not present | not present | |
| **31117** | 29.5 | 171.2 | 90 | 22-27 | 23-24-25 | 13 | 9-10-11-12 | without sperm | without sperm | |
| **31118** | 24.5 | 105.3 | 82 | 22-27 | 23-24-25 | 13 | 11 -- 12 | not present | not present | |
| **31119** | | | | | | | | | | |
| **31120** | 22 | 57.6 | 79 | | 23-24-25 | 13 | | | | semi-mature |
| **31121** | 30 | 120.6 | 90 | 22-27 | 23-24-25 | 13 | 9-10-11-12 | without sperm | without sperm | |
| **31122** | 23.5 | 50.3 | 83 | 22-27 | 23-24-25 | 13 | 9-10-11-12 | without sperm | without sperm | |
| **31123** | 13 | 20.1 | 78 | | | | | | | inmature |
| **31124** | | | | | | | | | | inmature, the earthworm was broken |
| **31125** | | | | | | | | | | the earthworm was broken |
| **31126** | | | | | | | | | | inmature, the earthworm was broken |

| | | | | | | | | | | |
|---|---|---|---|---|---|---|---|---|---|---|
| **31127** | 29 | 71.2 | 78 | 22-27 | 23-24-25 | 13 | 9-10-11-12 | not present | not present | |
| **31128** | 26 | 45.6 | 78 | 17-22 | 19-20-21 | 8 y 9 | 3-4-5-6 | without sperm | not present | maybe regenerated by the head |
| **31129** | 24 | 88.9 | 82 | | | | | | | inmature |
| **31130** | 30 | 135.2 | 92 | | 23-24-25 | 13 | | | | semi-mature |
| **31131** | 16 | 33.8 | 77 | | | | | | | inmature |
| **31132** | | | | | | | | | | |
| **31133** | | | | 22-27 | 23-24-25-1n26 | 13 | 9-10-11-12 | without sperm | without sperm | the earthworm was broken |
| **31134** | 30 | 91.7 | 88 | 22-27 | 23-24-25 | 13 | 9-10-11-12 | without sperm | without sperm | |
| **31135** | 24.5 | 63.8 | 77 | | 23-24-25 | 13 | | | | semi-mature |
| **31136** | 24 | 57.6 | 84 | | | | | | | inmature |
| **31137** | 15 | 35.2 | 91 | | | | | | | inmature |
| **31139** | 19 | 44.2 | 88 | | | | | | | inmature |
| **31140** | 28 | 109.6 | 93 | 22-27 | 23-24-25 | 13 | 9-10-11-12 | not present | not present | |
| **UCM-LT number** | **Length (mm)** | **Dry weight (mg)** | **Number of segments** | **Clitellum** | **Tubercula pubertatis** | **Male pore** | **Seminal vesicles** | **Spermatechae** | **Spermiducal funnels** | **Observations** |
| **31141** | 15 | 15.9 | 93 | | | | | | | inmature |
| **31142** | 19.5 | 36.5 | 90 | | | | | | | inmature |
| **31143** | | | | | | | | | | the earthworm was broken and inmature |
| **31144** | 23 | 57.2 | 73 | | | 13 | | | | semi-mature |
| **31145** | | | | | 24-25-26 | 13 | | | | the earthworm was broken and semi-mature |
| **31146** | 30 | 115.2 | 75 | 23-28 | 24-25-26 | 13 | 9-10-11-12 | without sperm | without sperm | |

| | | | | | | | | | | |
|---|---|---|---|---|---|---|---|---|---|---|
| **31147** | 24 | 63.8 | 70 | 23-28 | 24-25-26 | 13 | 10--11--12 | not present | without sperm | |
| **31148** | 20 | 48.3 | 81 | | | | | | | inmature |
| **31149** | 20 | 42.3 | 80 | | 24-25-26 | 13 | | | | semi-mature |
| **31169** | 27.5 | 57.6 | 82 | | 13 | | | | | semi-mature |
| **31170** | 27 | 66.8 | 87 | 22-27 | 23-24-25 | 13 | 9-10-11-12 | without sperm | without sperm | |
| **31171** | | | | | | | | | | inmature, the earthworm was broken |
| **31172** | | | | | | | | | | inmature, the earthworm was broken |
| **31173** | 18 | 25.2 | 82 | | | | | | | inmature |
| **31174** | | | | 21-25 | 22-23-24 | 12 | 9-10-11-12 | one on right side without sperm | not present | spermatophore between 19 and 20, the earthworm was broken |
| **31175** | 22 | 47.9 | 87 | 22-27 | 23-24-25 | 15 | 10--11--12 | not present | without sperm | |
| **31176** | 21.5 | 39.5 | 77 | | 23-24-25 | 13 | | | | semi-mature |
| **31177** | | | | | 23-24-25 | 13 | | | | semi-mature, the earthworm was broken |
| **31178** | 19 | 44.7 | 79 | | 23-24-25 | 13 | | | | semi-mature |
| **31179** | 20 | 42.9 | 79 | | 23-24-25 | 13 | | | | semi-mature |
| **31180** | | | | 22-27 | 23-24-25 | 13 | 11 -- 12 | without sperm | without sperm | the earthworm was broken |
| **31181** | 25 | 47.6 | 84 | 22-27 | 23-24-25 | 13 | 9-10-11-12 | without sperm | without sperm | |
| **31182** | | | | 22-27 | 23-24-25 | 13 | 9-10-11-12 | without sperm | without sperm | the earthworm was broken |
| **UCM-LT number** | **Length (mm)** | **Dry weight (mg)** | **Number of segments** | **Clitellum** | **Tubercula pubertatis** | **Male pore** | **Seminal vesicles** | **Spermatechae** | **Spermiducal funnels** | **Observations** |

| | | | | | | | | | | |
|---|---|---|---|---|---|---|---|---|---|---|
| **31183** | 25 | 78.9 | 85 | 22-27 | 23-24-25 | 13 | 9-10-11-12 | without sperm | no presente | |
| **31184** | 12 | 45.8 | 79 | | | | | | | inmature |
| **31185** | 19 | 57.2 | 61 | | | 13 | | | | semi-mature |
| **31186** | 24 | 54.6 | 81 | | | 13 | | | | semi-mature |
| **31187** | 20 | 48.5 | 73 | | 23-24-25 | 13 | | | | semi-mature |
| **31188** | 20 | 75.6 | 94 | 21-27 | 23-24-25-1n26 | 13 | 11 -- 12 | without sperm | without sperm | |
| **31189** | 17.5 | 48.2 | 87 | 22-27 | 23-24-25 | 13 | 11 -- 12 | without sperm | without sperm | |
| **31190** | 17 | 55.2 | 64 | 22-29 | 23-24-25-26 | 13 | 10--11--12 | without sperm | without sperm | |
| **31191** | 19 | 58.3 | 101 | 22-28 | 24-25-26 | 13 | 11 -- 12 | not present | without sperm | |
| **31192** | 19 | 64.1 | 88 | 22-27 | 23-24-25 | 13 | 11 -- 12 | without sperm | without sperm | |
| **31193** | 19.5 | 61.9 | 71 | 22-27 | 23-24-25 | 13 | 10--11--12 | without sperm | without sperm | |
| **31194** | 11 | 22.1 | 52 | | | | | | | inmature |
| **31195** | 11 | 24.3 | 78 | | | | | | | inmature |
| **31196** | 19.5 | 59.1 | 83 | | | | | | | inmature |
| **31197** | 9 | 23.1 | 75 | | | | | | | inmature |
| **31198** | 18 | 35.6 | 79 | 22-27 | 23-24-25 | 13 | 9-10-11-12 | without sperm | without sperm | |
| **31199** | 22 | 78 | 90 | 23-28 | 24-25-26 | 13 | 9-10-11-12 | no presente | no presente | |
| **31200** | 15 | 35.2 | 96 | 22-28 | 23-24-25 | 13 | 11 -- 12 | without sperm | without sperm | |
| **31201** | 8 | 16.4 | 36 | 22-27 | 23-24-25 | 13 | not present | without sperm | not present | |
| **31202** | 16 | 39.1 | 97 | 21-26 | 22-23-24 | 13 | 11 -- 12 | not present | not present | |
| **31203** | 11 | 33.5 | 83 | | | | | | | inmature |
| **31204** | 14.5 | 36.4 | 68 | | | | | | | inmature |
| **31205** | 13 | 37.5 | 57 | | 23-24-25 | 13 | | | | semi-mature |

| | | | | | | | | | | |
|---|---|---|---|---|---|---|---|---|---|---|
| **31206** | 28 | 87.3 | 80 | | 23-24-25 | 13 | | | | semi-mature, the earthworm was broken |
| **31207** | | | | | | | | | | inmature |
| **UCM-LT number** | **Length (mm)** | **Dry weight (mg)** | **Number of segments** | **Clitellum** | **Tubercula pubertatis** | **Male pore** | **Seminal vesicles** | **Spermatechae** | **Spermiducal funnels** | **Observations** |
| **31208** | 16 | 39.5 | 71 | 22-27 | 23-24-25 | 13 | 9-10-11-12 | without sperm | without sperm | |
| **31209** | 26.5 | 78.6 | 86 | 22-27 | 23-24-25 | 13 | 9-10-11-12 | one on each side without sperm | without sperm | |
| **31210** | 22.5 | 79.5 | 74 | 22-27 | 23-24-25 | 13 | 9-10-11-12 | not present | not present | |
| **31211** | 21.5 | 77.4 | 57 | 22-27 | 23-2425-1n26 | 13 | 11 -- 12 | not present | not present | |
| **31212** | 24 | 85.3 | 62 | 22-27 | | 13 | 10--11--12 | not present | not present | |
| **31213** | 20 | 57.2 | 83 | 20-25 | 22-23-24 | 11 | 11 -- 12 | not present | not present | |
| **31214** | 23 | 65.2 | 83 | 22-27 | 23-24-25 | 13 | 9-10-1-12 | without sperm | not present | |
| **31215** | 18 | 59.2 | 85 | 22-27 | 23-24-25 | 13 | 9-10-11-12 | not present | not present | |
| **31216** | 25 | 84.1 | 92 | 22-27 | 23-24-25 | 13 | 11 -- 12 | not present | not present | |
| **31217** | 28.5 | 95.2 | 90 | 22-27 | 23-24-25 | 14 | 9-10-11-12 | one on each side without sperm | not present | |
| **31218** | | | | | | | | | | inmature, the earthworm was broken |
| **31219** | 28 | 75.2 | 84 | | 22-23-24 | 13 | | | | semi-mature |
| **31220** | 23 | 49.3 | 91 | | | | | | | inmature |
| **31221** | 28 | 88.9 | 88 | | | 13 | | | | semi-mature |
| **31222** | 20 | 36.7 | 77 | | | | | | | inmature |
| **31223** | 25 | 59.8 | 91 | | | | | | | inmature |

| | | | | | | | | | | |
|---|---|---|---|---|---|---|---|---|---|---|
| **31224** | 21 | 36.8 | 82 | | | | | | | inmature |
| **31225** | 31 | 95.9 | 86 | | 23-24-25 | 13 | | | | semi-mature |
| **31226** | 23 | 96.7 | 91 | 22-28 | 23-24-25 | 13 | 9-10-11-12 | without sperm | without sperm | |
| **31227** | 21.5 | 75.8 | 87 | 22-28 | 23-24-25 | 14 | 10--11--12 | not present | without sperm | |
| **31228** | 25.5 | 93.2 | 87 | 22-27 | 23-24-25-26 | 13 | 9-10-11-12 | not present | not present | |
| **31229** | 42 | 193.6 | 84 | 22-27 | 23-24-25-26 | 13 | 9-10-11-12 | not present | not present | |
| **31230** | | | | | | | | | | the earthworm was broken |
| **31231** | 26 | 112.8 | 81 | 22-28 | 23-24-25-26-1n27 | 13 | 9-10-11-12 | without sperm | without sperm | |
| **UCM-LT number** | **Length (mm)** | **Dry weight (mg)** | **Number of segments** | **Clitellum** | **Tubercula pubertatis** | **Male pore** | **Seminal vesicles** | **Spermatechae** | **Spermiducal funnels** | **Observations** |
| **31232** | 21 | 75.1 | 83 | 22-28 | 23-24-25 | 13 | 9-10-11-12 | not present | not present | |
| **31233** | 19 | 43.4 | 79 | | | 15 | | | | semi-mature |
| **31234** | 21.5 | 87.9 | 65 | 22-28 | 23-24-25 | 13 | 9-10-11-12 | not present | not present | |
| **31235** | 22 | 57.3 | 89 | | | 15 | | | | semi-mature |
| **31236** | 18 | 47.2 | 77 | | | 13 | | | | semi-mature |
| **31237** | 27 | 118.5 | 86 | 22-28 | 23-24-25-1n26 | 13 | 11 -- 12 | not present | not present | |
| **31238** | 24.5 | 96.2 | 87 | 22-28 | 23-24-25 | 13 | 9-10-11-12 | not present | not present | |
| **31239** | 24.5 | 83.4 | 89 | 22-28 | 23-24-25 | 13 | 10 -- 12 | not present | not present | |
| **31240** | 25 | 87.2 | 87 | 22-28 | 23-24-25-26 | 13 | 11 -- 12 | no presentes | not present | |
| **31241** | 30 | 128.5 | 92 | 22-28 | 23-24-25-26 | 15 | 9-10-11-12 | not present | not present | |
| **31242** | 24.5 | 87.5 | 70 | 22-27 | 23-24-25 | 13 | 11 -- 12 | not present | not present | |
| **31243** | 20.5 | 74.9 | 82 | | 23-24-25 | 15 | | | | semi-mature |

| | | | | | | | | | | |
|---|---|---|---|---|---|---|---|---|---|---|
| **31244** | 16 | 47.6 | 51 | 22-27 | 23-24-25 | 13 | not present | not present | not present | |
| **31245** | 15 | 19.8 | 80 | 22-27 | 24-25-1n26 | 13 | 9-10-11-12 | not present | without sperm | |
| **31246** | 19 | 54.9 | 80 | 23-28 | 24-25-26 | 13 | 11 -- 12 | without sperm | without sperm | |
| **31247** | 20 | 36.7 | 68 | | | | | | | inmature |
| **31248** | 21.5 | 78.5 | 78 | | 23-24-25 | 13 | | | | semi-mature |
| **31249** | 21 | 47.2 | 83 | | | 13 | | | | semi-mature |
| **31250** | 24 | 75.2 | 77 | | | | | | | inmature |
| **31251** | 17 | 39.4 | 71 | | | | | | | inmature |
| **31253** | 14.5 | 47.6 | 77 | | | | | | | inmature |
| **31254** | 19 | 33.4 | 78 | | | | | | | inmature |
| **31255** | 17 | 37.2 | 63 | | | | | | | inmature |
| **31256** | 27 | 116.4 | 73 | 22-27 | 23-25 | 13 | 9-10-11-12 | not present | not present | |
| **31257** | 30 | 135.1 | 81 | 22-27 | 23-25 | 13 | left side: 11 right side: 11--12 | not present | not present | spermatophore between 15 and 16 |
| **UCM-LT number** | **Length (mm)** | **Dry weight (mg)** | **Number of segments** | **Clitellum** | **Tubercula pubertatis** | **Male pore** | **Seminal vesicles** | **Spermatechae** | **Spermiducal funnels** | **Observations** |
| **31258** | 28 | 97.4 | 85 | 22-27 | 22-25 | 13 | 10--11--12 | not present | not present | |
| **31259** | 28 | 114.2 | 82 | 23-29 | 24-27 | 13 | 11 -- 12 | not present | not present | |
| **31260** | 15 | 35.8 | 58 | | 23-25 | 13 | | | | semi-mature |
| **31262** | 17.5 | 36.7 | 89 | 22-27 | 23-25 | 13 | 9-10-11-12 | not present | not present | |
| **31263** | | | | 22-27 | 23-25 | 13 | 10--11--12 | not present | not present | the earthworm was broken |
| **31264** | 13 | 37.5 | 74 | | | | | | | inmature |
| **31265** | 18 | 59.4 | 79 | | 23-25 | 13 | | | | semi-mature |

| | | | | | | | | | | |
|---|---|---|---|---|---|---|---|---|---|---|
| **31266** | 18 | 36.5 | 77 | 22-27 | 23-25 | 13 | 11 -- 12 | not present | not present | |
| **31267** | 21 | 78.5 | 92 | 22-27 | 23-25 | 13 | 11 -- 12 | not present | not present | |
| **31268** | 18 | 39.6 | 82 | | 23-25 | 13 | | | | semi-mature |
| **31269** | 15 | 24.7 | 89 | | | | | | | inmature |
| **31270** | 18 | 47.2 | 63 | 22-27 | 23-24-25-1n26 | 13 | 10--11--12 | not present | not present | |
| **31271** | 22 | 89.6 | 76 | 22-27 | 23-24-25-1n26 | 13 | 9-10-11-12 | without sperm | not present | |
| **31272** | 21.5 | 64.7 | 83 | 22-27 | 23-24-25 | 13 | not present | not present | not present | |
| **31273** | | | | | 23-24-25 | 13 | | | | semi-mature, the earthworm was broken |
| **31274** | 17 | 58.2 | 79 | | 23-24-25 | 13 | | | | semi-mature |
| **31275** | 17 | 74.3 | 81 | 22-27 | 23-24-25 | 13 | not present | not present | not present | |
| **31276** | 15 | 47.2 | 71 | | | | | | | inmature |
| **31277** | 15 | 26.1 | 66 | | | | | | | inmature |
| **31278** | 28 | 35.4 | 79 | | | 13 | | | | semi-mature |
| **31279** | 25 | 89.4 | 88 | 22-27 | 23-24-25-1n26 | 15 | 11 -- 12 | not present | not present | |
| **31280** | | | | 22-27 | 23-24-25 | 13 | 9-10-11-12 | without sperm | without sperm | the earthworm was broken |
| **31281** | 25 | 94.3 | | | 23-24-25 | 13 | | | | semi-mature |
| **31282** | 30 | 117.6 | 92 | 22-27 | 23-24-25 | 13 | 11 -- 12 | without sperm | without sperm | |
| **UCM-LT number** | **Length (mm)** | **Dry weight (mg)** | **Number of segments** | **Clitellum** | **Tubercula pubertatis** | **Male pore** | **Seminal vesicles** | **Spermatechae** | **Spermiducal funnels** | **Observations** |
| **31283** | 26 | 86.8 | 84 | 22-27 | 23-24-25 | 15 | 11 -- 12 | not present | not present | |

| | | | | | | | | | | |
|---|---|---|---|---|---|---|---|---|---|---|
| **31284** | 18 | 54.9 | 51 | 22-27 | 24-25-26 | 13 | 11 -- 12 | not present | not present | |
| **31285** | 27 | 78.6 | 89 | 21-28 | 23-24-25-26-27 | 12 | 9-10-11-12 | not present | not present | |
| **31286** | 20 | 47.8 | 76 | | 23-24-25 | 13 | | | | semi-mature |
| **31287** | | | | | | | | | | inmature, the earthworm was broken |
| **31288** | 19 | 48.2 | 73 | | | | | | | inmature |
| **31289** | 22 | 57.5 | 92 | | | 13 | | | | semi-mature |
| **31290** | 17.5 | 35.9 | 69 | | | | | | | inmature |
| **31291** | 20 | 47.2 | 84 | | 23-24-25 | 13 | | | | semi-mature |
| **31292** | | | | 22-27 | 23-24-25 | 13 | 11 -- 12 | without sperm | without sperm | the earthworm was broken |
| **31293** | 24 | 75.21 | 89 | 22-27 | 23-24-25 | 13 | 11 -- 12 | not present | not present | |
| **31294** | 17 | 45.6 | 61 | 22-28 | 24-25-26 | 13 | 11 -- 12 | not present | not present | |
| **31295** | 16 | 33.2 | 77 | 21-17 | 23-24-25 | 13 | 11 -- 12 | not present | not present | |
| **31296** | 23 | 69.4 | 82 | 22-27 | 23-24-25 | 13 | 11 -- 12 | not present | not present | |
| **31297** | 22 | 47.8 | 85 | 22-27 | 23-24-25 | 13 | 11 -- 12 | not present | not present | |
| **31298** | 14 | 45.1 | 71 | | | | | | | inmature |
| **31299** | 27.5 | 98.4 | 84 | 22-27 | 23-24-25 | 13 | 9-10-11-12 | not present | without sperm | |
| **31300** | 24.5 | 101.6 | 82 | 22-28 | 24-25-26 | 13 | 9-10-11-12 | without sperm | without sperm | |
| **31301** | 22 | 125.8 | 75 | 22-27 | 23-24-25 | 13 | 11 -- 12 | not present | not present | |
| **31302** | 22 | 74.3 | 79 | 22-27 | 23-24-25 | 13 | 10--11--12 | not present | without sperm | |
| **31303** | 17 | 42.1 | 77 | | | 13 | | | | semi-mature |
| **31304** | 17.5 | 52.9 | 74 | 22-27 | 23-24-25 | 13 | 11 -- 12 | not present | without sperm | the earthworm was broken |

| | | | | | | | | | | |
|---|---|---|---|---|---|---|---|---|---|---|
| **31305** | | | | | | 13 | | | | semi-mature and the earthworm was broken |
| **31306** | | | | 22-27 | 23-24-25-1n26 | 13 | 10--11--12 | without sperm | without sperm | |
| **UCM-LT number** | **Length (mm)** | **Dry weight (mg)** | **Number of segments** | **Clitellum** | **Tubercula pubertatis** | **Male pore** | **Seminal vesicles** | **Spermatechae** | **Spermidular funnels** | **Observations** |
| **31307** | 15 | 12.3 | 81 | | | | | | | inmature |
| **31308** | 14 | 17.4 | 79 | | | | | | | inmature |
| **31309** | 13 | 21.9 | 78 | | | | | | | inmature |
| **31310** | 16 | 54.6 | 85 | | 13 | | | | | semi-mature |
| **31311** | 15 | 32.9 | 81 | | | | | | | inmature |
| **31312** | 16.5 | 25.8 | 81 | | | | | | | inmature |
| **31313** | 19.5 | 74.2 | 57 | 22-27 | 23-24-25 | 13 | 9-10-11-12 | without sperm | without sperm | |
| **31314** | | | | 22-27 | 23-24-25 | 13 | 10--11--12 | not present | not present | the earthworm was broken |
| **31315** | 23.5 | 71.3 | 77 | 22-27 | 23-24-25 | 13 | 10--11--12 | not present | not present | |
| **31316** | 25 | 79.5 | 92 | 22-28 | 23-24-25-26 | 13 | 11 -- 12 | not present | not present | |
| **30780** | 20 | 12.1 | 87 | 22-27 | 23-24-25 | 13 | 10--11-12 | not present | not present | |
| **30781** | 20 | 13.8 | 93 | 22-28 | 23-24-25-1n26 | 13 | 8-9-10-11 | not present | not present | |
| **30782** | 25 | 20.5 | 83 | 22-27 | 23-24-25 | 13 | 9-10-11-12 | without sperm | without sperm | |
| **30783** | 22.5 | 29.5 | 85 | 22-27 | 23-24-25 | 13 | 8-9-10-11 | without sperm | without sperm | |
| **30784** | 21 | 13.4 | 92 | 22-27 | 23-24-25 | 13 | 10--11-12 | without sperm | not present | |
| **30785** | 23.5 | 15.8 | 82 | 22-28 | 23-24-25-26 | 13 | 10--11-12 | without sperm | without sperm | |
| **30786** | 0.7 | 3.1 | 64 | | | | | | | inmature |

| | | | | | | | | | | |
|---|---|---|---|---|---|---|---|---|---|---|
| **30673** | 17 | 19.2 | 74 | | | 13 | | | | semi-mature |
| **30674** | 20 | 21 | 83 | | 23-24-25 | 13 | | | | semi-mature |
| **30675** | 12 | 10.2 | 90 | | | 13 | | | | semi-mature |
| **30676** | 15 | 6.1 | 82 | | | | | | | inmature |
| **30677** | | | | | | 13 | | | | semi-mature and the earthworm was broken |
| **30678** | 11 | 5.8 | 74 | | | | | | | inmature |
| **30679** | | | | | | | | | | the earthworm was broken and inmature |
| **30680** | 14 | 17.5 | 83 | | | 13 | | | | semi-mature, broken |
| **UCM-LT number** | **Length (mm)** | **Dry weight (mg)** | **Number of segments** | **Clitellum** | **Tubercula pubertatis** | **Male pore** | **Seminal vesicles** | **Spermatechae** | **Spermiducal funnels** | **Observations** |
| **30682** | | | | | | | | | | inmature, the earthworm was broken |
| **30683** | 14 | 8.7 | 95 | | | | | | | inmature |
| **31010** | 20 | 25.7 | 74 | 23-27 | 23-24-25 | 13 | 9-10-11-12 | not present | not present | |
| **31011** | 19 | 20.8 | 84 | 22-27 | 23-24-25 | 13 | 9 -- 10 | not present | not present | |
| **31012** | 22 | 19.8 | 85 | 22-27 | 22-23-24 | 13 | 9-10--11 | without sperm | not present | |
| **31013** | | | | | | | | | | the earthworm was broken and inmature |
| **31014** | | | | | | | | | | inmature and the earthworm was broken |
| **31015** | 9 | 3.7 mm | 83 | | | | | | | inmature |
| **31016** | 9 | 3.5 | 80 | | | | | | | inmature |
| **31017** | | | | | | | | | | |

| | | | | | | | | | | |
|---|---|---|---|---|---|---|---|---|---|---|
| **31018** | 10 | 4.1 | 74 | | | | | | | inmature |
| **31019** | | | | | | | | | | the earthworm was broken |
| **30704** | 22.5 | 131.8 | 94 | 22-27 | 23-24-25 | 13 | 11 -- 12 | not present | not present | |
| **30705** | 23 | 132.7 | 92 | 22-27 | 23-24-25 | 13 | 10--11-12 | not present | not present | |
| **30706** | | | | 22-27 | 23-24-25 | 13 | 10--11--12 | not present | not present | the earthworm was broken |
| **30707** | 29 | 151.3 | 85 | 22-27 | 23-24-25 | 13 | 11 -- 12 | not present | not present | |
| **30708** | 24 | 120.7 | 77 | 22-27 | 23-24-25 | 13 | 11 -- 12 | not present | not present | |
| **30709** | | | | 22-27 | 23-24-25 | 13 | 12 | not present | not present | the earthworm was broken |
| **30710** | 15 | 37.6 | 82 | | | | | | | inmature |
| **30711** | 26 | 100.4 | 91 | | 23-24-25 | 13 | | | | semi-mature |
| **30712** | 27 | 133.4 | 92 | 22-27 | 23-24-25 | 13 | 10--11--12 | not present | not present | |

Supplementary Table 7. Morphological traits of specimens included in the study.
